# D-SPIN constructs gene regulatory network models from multiplexed scRNA-seq data revealing organizing principles of cellular perturbation response

**DOI:** 10.1101/2023.04.19.537364

**Authors:** Jialong Jiang, Sisi Chen, Tiffany Tsou, Christopher S. McGinnis, Tahmineh Khazaei, Qin Zhu, Jong H. Park, Inna-Marie Strazhnik, Jost Vielmetter, Yingying Gong, John Hanna, Eric D. Chow, David A. Sivak, Zev J. Gartner, Matt Thomson

## Abstract

Gene regulatory networks within cells modulate the expression of the genome in response to signals and changing environmental conditions. Reconstructions of gene regulatory networks can reveal the information processing and control principles used by cells to maintain homeostasis and execute cell-state transitions. Here, we introduce a computational framework, D-SPIN, that generates quantitative models of gene regulatory networks from single-cell mRNA-seq datasets collected across thousands of distinct perturbation conditions. D-SPIN constructs probabilistic models of regulatory interactions between genes or gene-expression programs to fit the cell state distributions under different perturbations. Using large Perturb-seq and drug-response datasets, we demonstrate that D-SPIN models reveal key regulators of cell fate decisions and the coordination of distant cellular pathways in response to gene knockdown perturbations. D-SPIN also dissects gene-level drug response mechanisms in heterogeneous cell populations, elucidating how combinations of immunomodulatory drugs acting on distinct regulators induce novel cell states through additive recruitment of gene expression programs. The D-SPIN model further predicts cell state distributions under drug dosage combinations beyond the training data. D-SPIN provides a computational framework for constructing interpretable models of gene regulatory networks to reveal principles of cellular information processing and physiological control.

## Introduction

Cells simultaneously orchestrate a myriad of biological pathways to maintain homeostasis in response to diverse environmental cues [1, 2, 3]. The intricate networks of molecular interactions process information and control cell states. Understanding such networks leads to key biological and therapeutic challenges, including developmental control, novel cell state discovery, and rational intervention design [4, 5, 6, 7]. The construction of mathematical models that can predict the behavior of cells and tissues has been a central goal in computational biology [8]. Methods derived from artificial intelligence (AI), including deep neural networks and transformers, can be trained on large data sets to make predictions but lack interpretability due to the large number of parameters and the feedforward architectures that do not naturally map onto the structure of biochemical pathways [9, 10]. Building interpretable and predictive models of the information processing network could allow the design of therapeutic strategies for manipulating and programming cell states in the human body to treat diseases.

Gene regulatory network (GRN) models are mathematical models that predict genome-wide gene expression levels through networks of gene interactions mediated by different physical mechanisms, including promoter binding, post-transcriptional modifications, and protein interactions [11, 12, 13, 14]. Global reconstructions of gene regulatory networks in *E. coli*, yeast, and sea urchin embryos have revealed network modularity, recurring network motifs, and combinatorial logic through the assembly of transcription factor (TF) complexes at gene promoters [15, 16, 17, 18, 19, 20, 21]. However, in metazoans, gene network models have primarily focused on subcircuits involved in specific processes like T cell activation and embryonic stem cell differentiation, rather than global, cell-scale networks [5, 6, 7]. Furthermore, there are very few quantitative models of regulatory networks that can simulate and predict the response of a cell to genetic perturbations, therapeutics, or other signals. How gene regulatory networks globally modulate core cellular processes in parallel in response to environmental conditions remains poorly understood.

Historically, gene regulatory network reconstruction and modeling have been constrained by the number of biochemical or genetic measurements required to reconstruct networks with hundreds to thousands of interacting protein components. Classical approaches, such as pairwise biochemical binding measurements or associating genetic perturbations with impacted genes, enable the identification of gene regulators separately [22, 23]. The regulatory interactions are consolidated into pathways and integrated into diagrams of network models [24, 25, 26, 27]. However, for the global analysis of gene regulatory networks, perturbation approaches require knocking out hundreds to thousands of genes while monitoring transcription across thousands of genes. Traditionally, the experimental realization of genome-wide network reconstruction through genetic perturbation has been limited by experimental scale when genes were knocked out in bulk assays.

Developments in single-cell genomics and perturbation barcoding circumvent some of the conventional limitations of perturbation-driven network reconstruction. Perturbation multiplexing approaches, including Perturb-seq, click-tags, and MULTI-seq, allow the transcriptional state of each cell in a population to be measured across thousands of different genetic, drug-treatment, and signaling conditions [28, 29, 30, 31, 32, 33]. These experimental approaches identify the perturbation delivered to every single cell while simultaneously measuring changes in the entire transcriptome through single-cell mRNA-seq, enabling large-scale interrogation of cellular perturbation response to diverse perturbations. A core challenge is developing computational methods to integrate data from thousands of such experiments into a gene regulatory network model that can classify perturbations, map the flow of information across a regulatory network, simulate the interaction between perturbations with the network, and predict cellular responses to novel perturbations.

Here, we introduce a mathematical modeling and network inference framework, D-SPIN (Dimension-scalable Single-cell Perturbation Integration Network), which constructs regulatory network models directly from single-cell perturbation-response data. D-SPIN constructs a probabilistic graphical model on single genes or gene programs to infer regulatory interactions between genes and applied perturbations. The model can simulate the distribution of cell states from the inferred regulatory network, thus providing a circuit/pathway-level interpretation of the data that is sufficient to produce the observed gene expression. The inferred regulatory network achieves superior accuracy compared to existing methods on both synthetic datasets and large single-cell perturbation datasets with hundreds to thousands of genes. While many popular network inference methods only apply to TF regulations as they use motif or chromatin accessibility information, D-SPIN applies to general types of interactions, and the discovered interactions include post-transcriptional regulation and phosphorylation in signaling pathways. D-SPIN can accommodate a wide range of different perturbation types, including genetic perturbations, small molecules, signaling conditions, and even physiological states of health and disease.

We applied D-SPIN to construct gene regulatory network models from experimental datasets with thousands of perturbations and millions of cells, including two of the largest single-cell mRNA-seq datasets in existence: a genome-wide Perturb-seq experiment on the K562 chronic myelogenous leukemia cell line [29] and a new human immune cell drug-response experiment we performed. The integrated regulatory network models of K562 cells reveal key regulators of erythroid/myeloid fate choices and global organizing principles of homeostatic responses to perturbations. In our profiled responses of human immune cells to immunomodulatory drugs, D-SPIN identifies gene-level signatures that classify drugs with different mechanisms of action. D-SPIN further demonstrates that drug combinations can generate novel macrophage states through additive recruitment of gene expression programs, which is mediated by convergent, independent pathways regulating the anti-inflammatory M2 gene program. D-SPIN also extrapolates beyond training data and predicts the immune cell state distributions of any dosage combinations from single-drug responses and one combination experiment. Broadly, D-SPIN provides a computational framework for large-scale modeling of gene regulatory networks across cell types and physiological conditions, as well as an interpretable and generative representation of cell states for cell-scale models.

## Results

### Integrating information from perturbations reveals hidden regulatory interactions

To simulate the cell states with regulatory interaction between genes, a core challenge is uncovering hidden conditional dependence between genes. These hidden interactions do not affect the network states under wild-type conditions, resulting in the problem of alternative or indistinguishable models. As a toy model, in a simplified four-pathway network, pathways A and D constitutively inhibit genes B3 and C3 in pathways B and C, respectively (Figure 1A(i), Figures S1A and S1B, STAR Methods 2.6). In the natural state of the network, since B3 and C3 are persistently suppressed by multiple inhibitory interactions, their expression becomes uncorrelated with their regulators, such as A3, B2, C2, and D3. Consequently, these gene pairs have low correlation and mutual information and are difficult to infer their regulatory relationships (Figures S1C and S1D).

**Figure 1:**
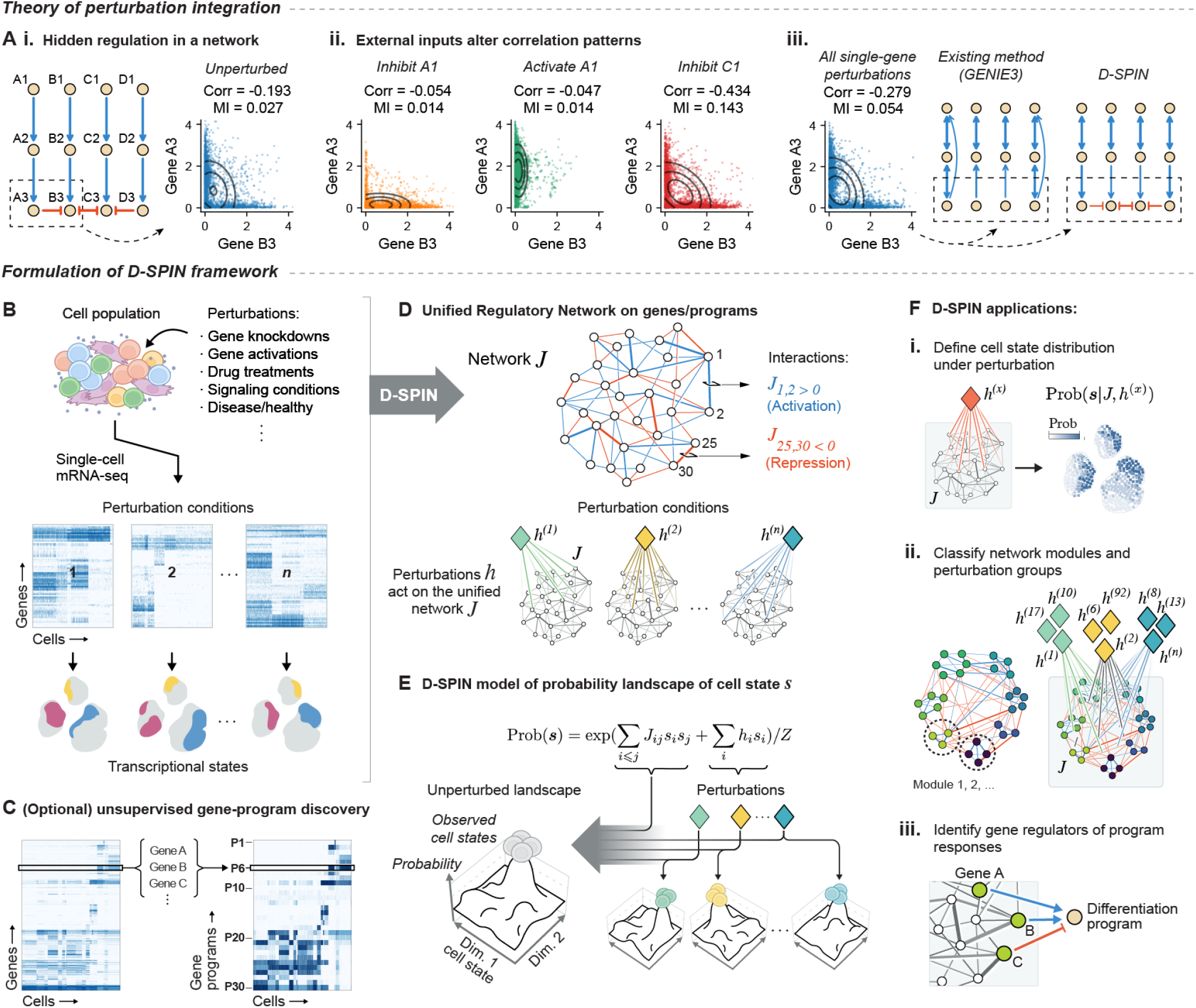
D-SPIN constructs unified gene regulatory network models from single-cell mRNA-seq data across perturbation conditions. (A)(i) A computational example of regulatory interactions hidden by other pathways. Gene A3 and B3 only have a weak negative correlation (Corr) and low mutual information (MI). (ii) External perturbation inputs alter the correlation pattern between genes, providing the possibility to decode hidden regulatory interactions, while computation methods are needed. (iii) Compared with existing top-performing methods, such as GENIE3, D-SPIN achieves more accurate network reconstruction and recovers the hidden regulations from perturbations. (B) D-SPIN accepts as input single-cell mRNA-seq data from a cell population profiled across different perturbed experimental conditions, such as genetic perturbations, drug treatments, signaling conditions, or disease/healthy conditions. (C) To enhance interpretability, apart from single-gene network constructions, D-SPIN contains an optional pipeline of unsupervised gene program discovery. (D) D-SPIN constructs a unified regulatory network ***J***, whose edges represent inferred interactions between genes or gene programs. D-SPIN also infers interactions between the network and applied perturbations. (E) Conceptually, the probabilistic model inferred by D-SPIN defines a probability landscape over all cell states ***s***. The pairwise interaction ***J*** between genes/programs defines a basis landscape which can be further modulated by sample-specific perturbation response vectors ***h***^(*n*)^ through tuning individual gene/program activities. The architecture also enables D-SPIN to scale to large datasets with millions of cells through a parallelized inference procedure. (F) D-SPIN yields a probabilistic model that can be interpreted as a gene regulatory network. The D-SPIN model can be applied to (i) estimate the distribution of transcriptional states defined by the regulatory network model under a specific condition, (ii) reveal gene regulatory network organization, such as modularity and sub-network architecture, and classify perturbation responses in the network context, and (iii) identify gene regulators of the program-level responses.

External perturbations offer an opportunity to reveal such masked interactions by exposing previously hidden dependencies between variables. Under perturbed conditions, these interactions now become impactful. In the four-pathway network, although being inhibited by another node C3, the expression level of B3 can be modulated by activating or inhibiting pathway A. Further, when the perturbation shuts down pathway C, the inhibition from A3 becomes prominent, leading to a significantly strengthened negative correlation and elevated mutual information between A3 and B3 (Figure 1A(ii)).

However, integrating the information between multiple perturbations has been a computational challenge, as the impact of perturbations can only be interpreted in the context of the network. A global joint model of network and perturbations is required to combine multiple perturbation observations into a coherent network reconstruction. Existing top-performing methods, such as GENIE3, GRNBoost2, and PIDC, focus on predicting single-gene expression state or information between gene pairs [34, 35, 36, 37]. However, none of them constructs a global network model for the observed state distribution, and no perturbation information is explicitly included. Therefore, these existing methods are not suitable for integrating information from multiple perturbation conditions and cannot recover these hidden interactions (Figure 1A(iii)).

To address the challenge, we developed the computational framework D-SPIN. D-SPIN constructs a probabilistic model that encodes the distribution of transcriptional states within a cell population across different perturbed conditions (Figures 1B-1E, STAR Methods 1.1). Mathematically, D-SPIN builds a condition-dependent spin network model, a specific form of Markov random field, to simulate gene interactions, where perturbations selectively activate certain cell states in analogy to the retrieval of associative memory of Hopfield networks (Figure 1E) [38]. Spin network models were initially introduced in physics to study collective behaviors and have been applied to complex systems in machine learning and biology, such as neural networks, bird flocks, and protein structures [39, 40, 38, 41, 42, 43].

Given all single-gene perturbations, D-SPIN recovered the correct architecture of the four-pathway model by integrating information from perturbations. In contrast, other methods cannot identify the hidden interaction of A3-B3, B3-C3, and D3-C3 when given the same perturbation data (Figure 1A(iii)). While the accuracy of network construction with D-SPIN is similar to existing methods without perturbations (Figures S1D), the unique architecture of D-SPIN extracts information from perturbations and significantly improves the inference accuracy over other methods. D-SPIN is the only method that identifies those hidden inhibitory interactions, and further assigns correct directions for most regulatory interactions (Figures S1E). D-SPIN achieves an average directed network accuracy (area under precision-recall curve, AUPRC) of 0.77 compared with 0.44 of other methods (Figure S1F).

### D-SPIN infers a unified regulatory network model from single-cell perturbation data

The application of spin network models to single-cell data has been limited due to the mathematical and computational challenges of inferring model parameters on large datasets [42, 43, 44, 45]. To enable information integration in large perturbation experiments with thousands of conditions, we leveraged a factorization of the spin network equation to split the learning problem into two parts: (1) construction of a unified gene regulatory network model and (2) inference of how each perturbation interacts with the unified network model (STAR Methods 1.2). For large networks, we further simplified the objective function by estimating conditional dependence between genes using pseudo-likelihood [46, 47, 48, 49]. These efficient computational procedures of D-SPIN enable network inference for up to thousands of genes and millions of single cells (STAR Methods 1.3).

As gene-level networks with thousands of nodes are difficult to interpret, we also designed D-SPIN to construct reduced-dimensional models by modeling interactions between gene programs, i.e., co-regulated groups of genes [50, 51, 52]. Cells regulate their transcriptional states by modulating transcription factors that impact sets of genes related to differentiation state, cellular pathways, and core physiological processes. Building upon published algorithms, D-SPIN contains an automatic pipeline for identifying gene programs from single-cell data based on unsupervised orthogonal nonnegative matrix factorization (oNMF) and phenotypic gene-program annotation (Figure 1C, STAR Methods 1.9) [53, 54, 55, 56]. D-SPIN can also accommodate pre-defined gene sets from prior biological knowledge (STAR Methods 1.10).

The program-level and gene-level networks constructed by D-SPIN provide complementary views of the regulatory architecture under perturbation responses. The program-level model provides a circuit/pathway-level representation of the data by the inferred interaction between these programs and their interaction with applied perturbations. As a probabilistic graphical model, D-SPIN can simulate the distribution of transcriptional states in a cell population under normal and perturbed conditions using these interactions (Figure 1F(i)). Therefore, D-SPIN dissects the uninterpretable cell state shifts in low-dimensional embeddings like principal component analysis (PCA) or UMAP into the action of perturbation onto specific nodes of the regulatory network, and allows making predictions for unobserved conditions. The program-level network model also reveals the internal coordination between physiological functions in response to perturbations to maintain homeostasis, classify gene programs and perturbation input into functional modules and perturbation groups (Figure 1F(ii))

At the gene level, we typically restrict the model to regulatory genes such as TFs, kinases, and phosphatases to model and analyze gene regulation. The gene-level network model highlights core signaling hubs, reveals information flow through signal transduction pathways, and provide fine-grained gene signature of perturbation responses. Further, to connect the program-level network model with gene-level models, we also developed algorithms to statistically identify key regulators from the gene-level network that control the perturbation responses of gene programs (Figure 1F(iii), STAR Methods 1.4). The program representation is constrained to assign each gene to a single program to avoid confounding interactions between programs, which may misrepresent versatile regulators controlling multiple activities. Thus, we designed the regulator discovery algorithm to operate independently of the gene program compositions, allowing D-SPIN to characterize shared regulatory control across pathways while preserving a non-overlapping program-level decomposition.

### D-SPIN constructs an accurate, interpretable, and generative model of HSC perturbation response

To assess the network model constructed by D-SPIN, we applied D-SPIN to a synthetic dataset of hematopoietic stem cell (HSC) differentiation using the network simulation and benchmarking framework BEELINE [37]. In the HSC network, there are two key regulators of cell fates: the transcriptional factor (TF) SPI1 (PU.1) controls granulocyte/monocyte differentiation, and GATA1 controls megakaryocyte/erythrocyte fates (Figure 2A). From biologically identified regulatory networks, we used the BEELINE framework to generate synthetic gene-expression profiles from the HSC network for 22 perturbation conditions, encompassing knockdown/activation of each network node individually (Figures 2B and 2C, STAR Methods 2.6). The simulated single-cell expression profiles are classified into 7 different cell state designations with the known TF markers, including monocytes (Mono), granulocytes (Gran), erythrocytes (Ery), and megakaryocytes (Mega) [57].

**Figure 2:**
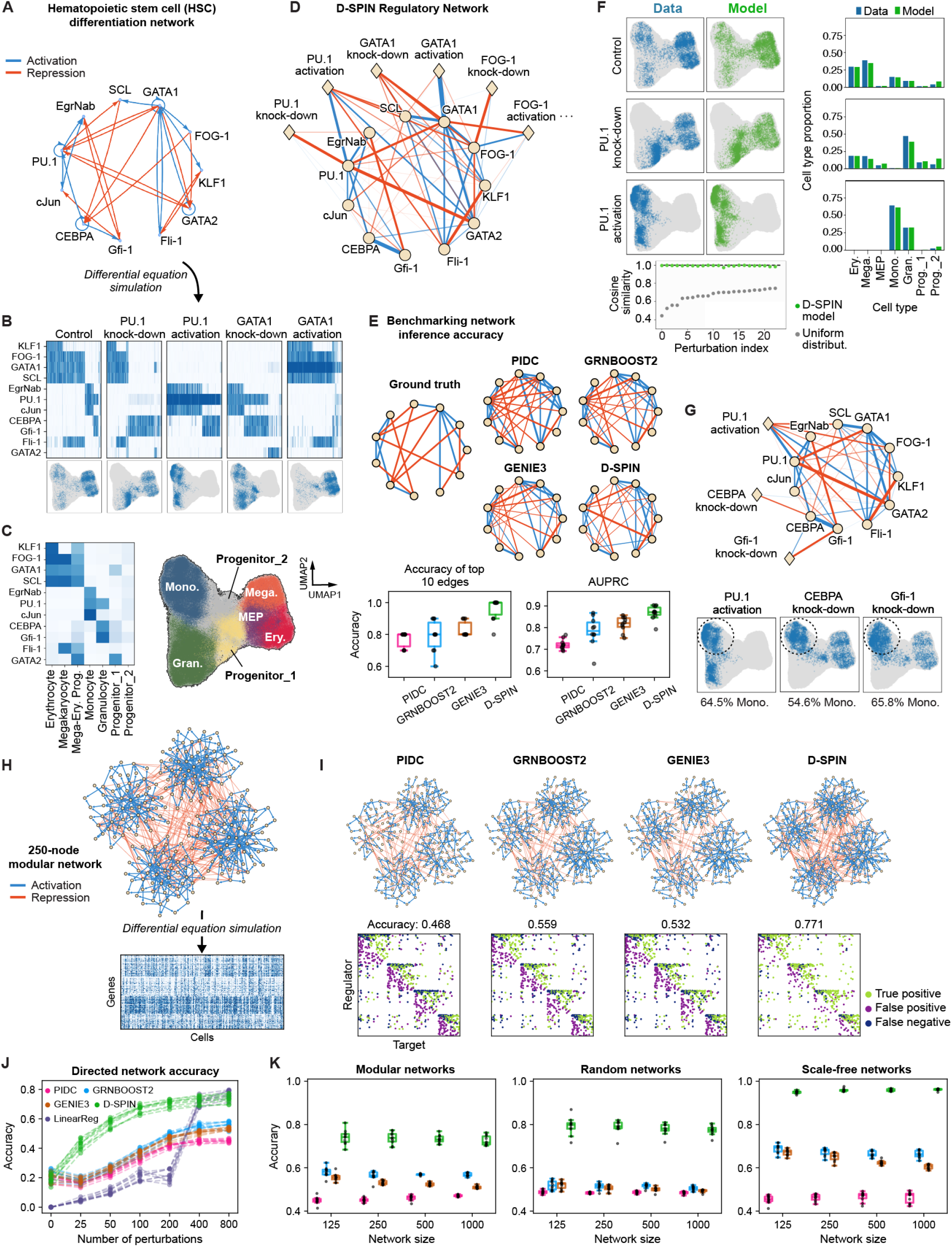
D-SPIN achieves state-of-the-art performance on network inference benchmarking tasks and reveals network-level mechanisms of cell fate modulation by TF perturbations. (A) HSC regulatory network model contains 11 TFs that interact through activations (blue) and repressions (red) to modulate the differentiation of HSCs. (B) Example simulated gene-expression profile heatmaps from the network across a series of single-gene knockdown and activation conditions. (C) Heatmap of average TF expression and UMAP embeddings for 7 clusters of cell states generated across all simulated conditions. (D) The network diagram shows the unified regulatory network model inferred by D-SPIN. Edges with diamond markers show D-SPIN-inferred perturbation vectors that infer how knockdown or activation of TFs (e.g., GATA1 knockdown) impacts the regulatory network through up- or downregulation of TFs. (E) Diagrams of the true network and inferred networks by D-SPIN and other state-of-the-art methods. The box plots quantify the accuracy of the top 10 inferred edges and AUPRC across 10 random repeats of simulations. (F) D-SPIN models generate cell state distributions highly consistent with the simulated single-cell data by applying perturbations to the underlying network, as quantified by (left) UMAP embeddings, (right) bar plots of cell type proportions, and (bottom) cosine similarity between cell type distributions. (G) (top) Network diagram and (bottom) UMAP embedding of three different perturbations that generate an increased monocyte population. Through a reasoning framework on the network, the D-SPIN model reveals network edges that mediate the response. PU.1 activation directly interacts with monocyte genes, while CEBPA and Gfi-1 knockdown act indirectly through the interaction of Gfi-1 and EgrNab. (H) (top) Network diagram of an example 250-node directed modular network with 4 mutually inhibiting modules, and (bottom) simulated single-cell data from the network for evaluating the accuracy and scalability of network inference. (I) Diagrams of the inferred directed network of different methods, as shown in (top) the subnetwork of correctly-inferred edges and (bottom) the adjacency matrices with true positives, false positives, and false negatives. (J) The plot shows the inference accuracies of each method when given different numbers of perturbations in 10 randomly generated modular networks. The network inference accuracy generally increases with the perturbation number. Compared with other methods, D-SPIN is significantly more effective at integrating information from perturbation response and achieves high accuracy. The linear regression model of perturbation response has low accuracy at small perturbation numbers, but has high accuracy similar to D-SPIN with large perturbation numbers. (K) Box plots quantify the directed network inference accuracy of three prevalent network topologies with different sizes of 125, 250, 500, and 1000 under 800 perturbations in 10 random repeats. D-SPIN achieves significantly increased accuracy compared with other network inference methods across network sizes.

The network topology inferred by D-SPIN achieves higher accuracy compared with existing network inference algorithms. We applied D-SPIN to construct a gene-level network model from the simulated data and compare the network structure with 3 different network reconstruction methods (PIDC, GRNBoost2, and GENIE3) that were top performers in the BEELINE benchmarking study [34, 35, 36, 37]. For D-SPIN, on average, 0.96 of the top 10 edges found by the model are correct across inference runs, as compared to 0.77 to 0.83 for PIDC, GRNBoost2, and GENIE3 (Figures 2D and 2E). D-SPIN achieves an average precision (AUPRC) of 0.87, compared with 0.72 to 0.82 for other methods.

Inferring directions of interactions from only stationary data has been a difficult and persistent problem [34]. D-SPIN can also infer a directed network model with the pseudolikelihood algorithm. The direction of interaction is inferred from the asymmetry between the predicting power of two factors (STAR Methods 1.3). D-SPIN achieves an average precision (AUPRC) of 0.77 on directed network inference of the HSC dataset, compared with 0.47 to 0.57 for other methods (Figure S2A). Despite the strength of D-SPIN in directed network inference, we focus our following analysis on undirected networks, as mathematically, there are fundamental difficulties in simulating stationary cell state distribution from directed graphical models with feedback regulations (STAR Methods 1.3) [58, 59, 60].

D-SPIN is a generative model that can simulate the distribution of transcriptional states of cell populations under perturbations. Most existing network inference methods are not generative, as they use either regression or information-theoretic measures to identify candidate gene interactions [37]. For the HSC datasets, D-SPIN infers a response vector for each perturbation condition that characterizes how the perturbation up- or downregulates each TF in the network to produce the observed state distribution (Figure 2D, Figure S2B). The distributions generated by the network and response vectors in the D-SPIN model are highly concordant with the data distribution across all perturbation conditions, as visualized by the UMAP embedding and cell state distribution comparison (Figure 2F, Figure S2C). Quantitatively, the cell state distributions by D-SPIN were all above 0.96 cosine similarity with the simulated data. Thus, for the HSC network, D-SPIN constructed a quantitative model of how the cell population is generated by the underlying gene regulatory network.

As an interpretable and generative model, D-SPIN can be analyzed to gain insight into how edges within a D-SPIN network model mediate the response to applied perturbations. We developed a formal framework to compute how the knockdown or activation of specific nodes impacts the cell state distribution generated by the regulatory network and to define how individual edges contribute to the perturbation response of the network (STAR Methods 2.7). For example, both PU.1 activation and the knockdown of either CEBPA or Gfi-1 led to an increased proportion of monocyte cell states, but with different inferred network mechanisms (Figure 2G). Activating PU.1 increases the activity of other monocyte genes (EgrNab and cJun) via the PU.1-EgrNab and PU.1-cJun edges. In contrast, under Gfi-1 knockdown, although Gfi-1 has both positive and negative interactions with the PU.1-EgrNab-cJun module, edge-sensitivity analysis shows that the negative interaction Gfi-1-EgrNab has a stronger impact on PU.1. The knockdown increases the activity of PU.1, EgrNab, and cJun, therefore boosting the monocyte population (Figure S2D(i)). Likewise, CEBPA knockdown reduces Gfi-1 activity through the CEBPA-Gfi-1 interaction, producing similar increased monocyte states as in Gfi-1 knockdown (Figure S2D(ii)). Overall, PU.1 activation induced increased monocyte states through the positive interactions PU.1-EgrNab and PU.1-cJun, while increased monocyte states in CEBPA or Gfi-1 knockdown were mediated by the negative edge Gfi-1-EgrNab. Therefore, similar cell-state distribution changes of increased monocyte states can be achieved by two different network-level mechanisms in the D-SPIN model.

Thus, D-SPIN, using simulated data, can infer an accurate and generative regulatory network model of a single-cell population by integrating information across perturbation conditions. The network model provides a map of how the distribution of cell states in the underlying HSC cell population is generated through interactions between internal regulators. The D-SPIN model can simulate the cell population and hypothesize network edges and network-level mechanisms that might mediate the observed cell population shifts under perturbations.

### D-SPIN achieves superior accuracy in large-scale regulatory network inference

To demonstrate the scalability of D-SPIN for building accurate network models with perturbation information, we assessed the accuracy of directed networks inferred by D-SPIN against other state-of-the-art methods on large-scale simulated networks with hundreds to a thousand nodes and various topologies. We explored three distinct network architectures: highly modular networks, Erdős–Rényi (ER) random networks, and scale-free networks (Figure 2H, Figure S2F, STAR Methods 2.8).

Each network topology captures specific characteristics that occur in real biological networks. In modular networks, activating interactions are enriched in the same module, representing groups of coexpressed genes that often perform coherent biological functions. Modular networks present challenges for network inference procedures because gene coexpression makes it difficult to define the precise regulators that control module activation and inhibition [19, 51, 61, 62]. ER networks randomly connect any pair of genes, which are considered the most general random network topology [63]. Scale-free networks contain hub nodes with a high number of interactions, and some biological networks exhibit similar statistics, such as metabolic networks [64, 65]. We generated each topology with 125, 250, 500, and 1000 nodes. Perturbation design for large networks remains an open challenge; therefore, we examined several heuristic perturbation design strategies. The most informative perturbations are combinatorial random perturbations with both activations and inhibitions (Figure S2E). Specifically, we use perturbations that are independent, normally distributed random variables on each network node (STAR Methods 2.8).

D-SPIN achieved significantly higher accuracy in directed network inference compared with other methods, including PIDC, GRNBoost2, and GENIE3 (Figure 2I, Figure S2G, STAR Methods 2.10). In the example 250-node modular network, the accuracy of the D-SPIN inferred network is 0.771, much higher than the best of the other three methods, 0.559. The accuracy improvement came from D-SPIN’s ability to effectively integrate information from perturbations. With no perturbation, the accuracy of D-SPIN is similar to that of existing methods (Figure 2J, Figure S2H). However, with the increasing number of perturbations, the average accuracy of D-SPIN continues to rise from an average of 0.191 to 0.737 with 800 normally distributed random perturbations (Figure 2J). Applying other network inference methods (PIDC, GRNBoost2, GENIE3) to the dataset of pooling all conditions together also increases the accuracy of inference, demonstrating the importance of perturbation in network inference. Nonetheless, these methods cannot integrate the information of the applied perturbation, and their accuracy is significantly lower than D-SPIN (Figure 2J, Figure S2H).

Comparison between D-SPIN and linear regression (LR) model of perturbation response reveals the source of information for network inference. The LR model computes gene regulation as the coefficients that link applied perturbations to gene expression changes. Mathematically, the LR model ignores gene covariation structure induced by gene interactions across individual cells, but efficiently builds an input-output map between the perturbations and average expression changes. In contrast, D-SPIN has an explicit model of the internal network that generates the observed responses from perturbation input. Thus, the accuracy of LR models is much lower than D-SPIN and other methods at small perturbation numbers, but significantly increases when the number of perturbations surpasses the network size (Figure 2J, Figure S2H). The similar accuracy between D-SPIN and the LR model at a large perturbation number suggests that the average perturbation responses have become the primary source of information. The performance improvement of D-SPIN over the LR model at a small perturbation number is the information D-SPIN obtained by additionally modeling the interactions between genes.

The superior accuracy of D-SPIN applies to all tested network topologies and scales up to a thousand genes. In 1000-gene networks, the directed network inferred by D-SPIN has average accuracies of 0.726 in modular networks, 0.773 in ER networks, and 0.961 in scale-free networks, compared to 0.569, 0.505, and 0.663 for the best of the other three methods (Figure 2K). The most striking example is scale-free networks, where the networks contain activating interactions from the hub nodes, and the unperturbed steady state distributions are all genes being turned on. Without perturbations, the accuracy of all methods is near 0.1. In comparison, D-SPIN achieves an accuracy higher than 0.95 by integrating information from perturbations, making almost precise network edge predictions (Figures S2G and S2H). The performance difference demonstrates the importance of combining all perturbations into the same model.

Additionally, the training algorithms of D-SPIN are highly efficient and faster than the other three methods by orders of magnitude in large-scale single-cell datasets containing hundreds of thousands of cells. We assessed the scalability of different network inference methods with the number of cells (Figure S2I, STAR Methods 2.9). D-SPIN becomes the fastest method for datasets exceeding 16,000 cells, a number that most current single-cell studies surpass. Remarkably, at 256,000 cells with 2 CPU cores, D-SPIN finished in 6 hours while all other methods could not finish within a week.

### The directed TF-target network inferred by D-SPIN corresponds well with ChIP-seq databases

D-SPIN builds regulatory network models directly from large-scale single-cell datasets, providing mechanistic insights into the hidden associations between genes and pathways. The model allows us to generate hypotheses about the cellular pathways that can motivate further experimental validations, such as identifying interactions between different pathways and nominating key regulators of biological processes. Perturb-seq is a single-cell genomics technique that enables highly parallelized genetic screens by combining the knockdown of thousands of genes using CRISPR/Cas9 with a transcriptome-scale single-cell mRNA-seq readout [28, 70]. We applied D-SPIN to one of the largest genome-wide Perturb-seq datasets of human chronic myelogenous leukemia (CML) (K562) cell lines, where 9,867 genes were knocked down individually across 2 million single cells [29].

To assess the network reconstruction quality of D-SPIN, we inferred directed gene-level regulatory networks and compared the results with independent biological measurements from ChIP-seq databases. ChIP-seq data measures the binding of transcription factors (TFs) to target gene promoters and has been used in the literature to assess the quality of inferred gene regulatory network models [37]. We created datasets of genes with active TFs and 500, 1000, and 1500 highly variable genes (HVGs), and evaluated the network inference of D-SPIN together with PIDC, GRNBoost2, GENIE3, CellOracle, and z-score on all cells in the datasets, including perturbed cells [37, 71, 72] (STAR Methods 2.11). D-SPIN achieved leading correspondence with ChIP-seq data quantified by accuracy rates (fold improvement over random guess) and AUROC (area under the receiver operating characteristic curve) across all three datasets (Figure 3A). Notably, associating gene responses with the perturbed gene by differential expression (DE) or z-score was shown to be effective in flow cytometry or bulk sequencing data [72, 73], but has limited performance in our evaluation, likely due to the high noise level in the Perturb-seq data.

**Figure 3:**
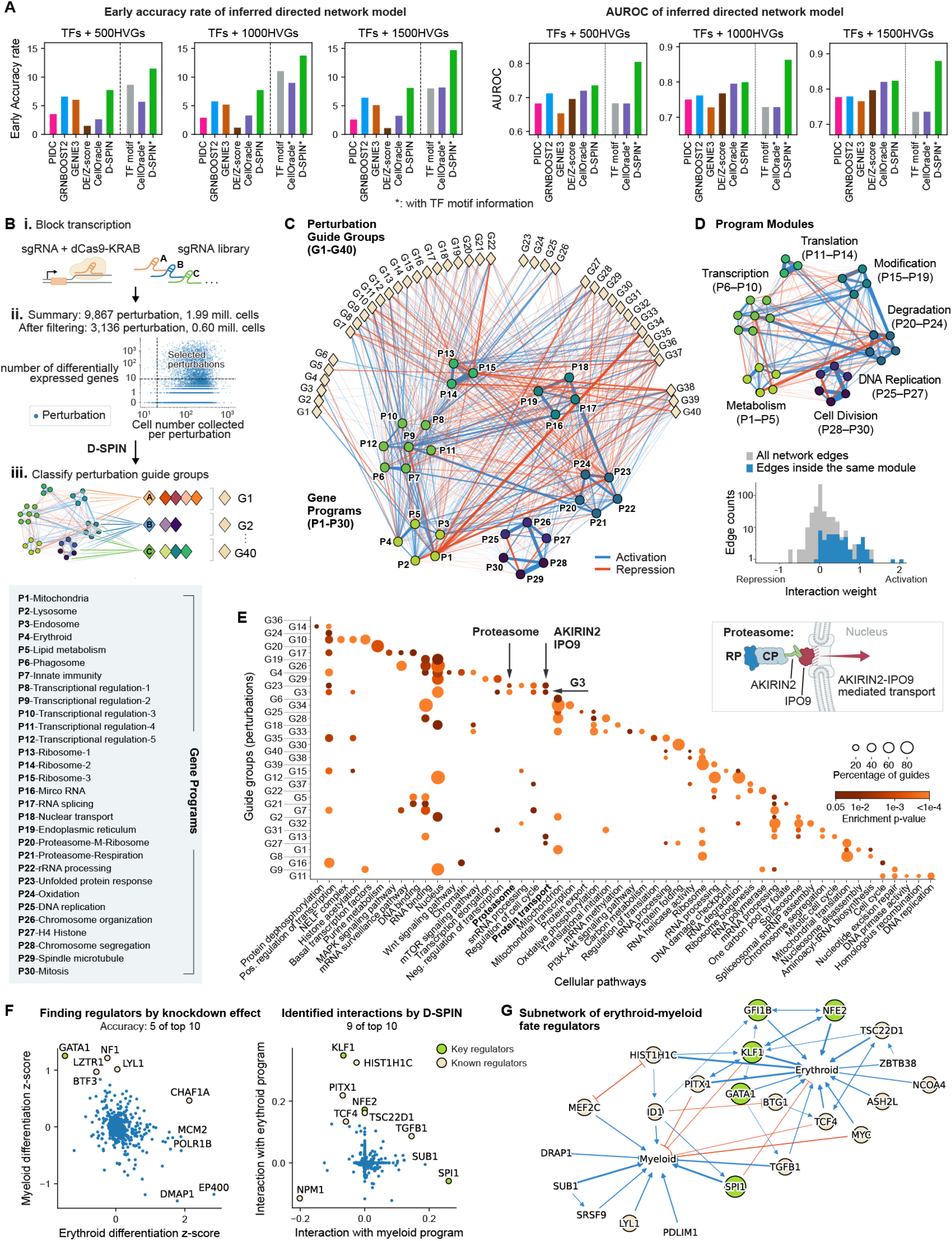
Regulatory network models of genome-wide Perturb-seq data reveal the architecture of cellular pathways and regulators of differentiation fate control. (A) Bar plots quantify the early accuracy rate and AUROC of the correspondence between inferred network and ChIP-seq databases for multiple datasets generated from the genome-wide Perturb-seq dataset [29]. Methods with the (*) symbol utilize TF-motif binding information as prior knowledge in the network inference. DE/z-score uses the differentiation expression/z-score to link responding genes to perturbed regulators. (B) Schematics of the experiments of the genome-wide Perturb-seq dataset. (i) 9,867 gene knockdown perturbations are individually delivered to ∼2M human K562 cells using guide RNAs. (ii) Scatter plots of the number of differentially expressed genes (DEGs) and the number of cells for each perturbation. 3,136 perturbations passed filtering. (iii) The D-SPIN model constructs a unified regulatory network model and infers interactions between each perturbation and the network, classifying gene-knockdown perturbations into groups. (C) The network diagram shows the inferred regulatory network model (*J* matrix) between 30 gene-expression programs (P1-P30, circles), as well as the interactions between programs and 40 groups of gene-knockdown perturbations (G1-G40, diamonds) encoded in response vectors (*h* vectors). Interactions are rendered as positive (blue) or negative (red) edges, with the thickness scaling with the strength of interactions. (left box) Gene programs are functionally annotated through gene ontology annotation tools and manual lookup [66, 67, 68]. (D) (top) The regulatory network model exhibits a modular structure with tightly connected subnetworks. The 7 network modules can be assigned physiological functions based on the gene programs they encompass. (bottom) The histogram quantifies the distribution of all network edges and edges inside the same module. Edges inside modules are mostly positive interactions and contain the majority of strong positive interactions in the network. (E) Bubble plot shows enriched cellular pathways in gene ontology and KEGG pathway from databases [66] for each guide RNA group. Gene targets in the same guide group identified by D-SPIN are involved in similar pathways or potentially reveal associations between pathways, such as (arrow pointer) protein transport genes AKIRIN2 and IPO9 that were recently found to (inset schematics) mediate the import of proteasome into the nucleus [69]. (F) Scatter plots show (left) gene knockdowns that have the strongest effect on the erythroid and myeloid differentiation programs quantified by z-score; (right) genes with the strongest interactions with the erythroid and myeloid program inferred by D-SPIN. 9 of the top 10 regulators identified by D-SPIN are known regulators associated with erythroid/myeloid differentiation. (G) Rendering of the subnetwork of activating regulators of the erythroid and myeloid programs inferred by D-SPIN. Most genes in the subnetwork are known regulators, and several are key regulators of the differentiation.

Regulatory relationships between TFs and targets are also reflected in the binding motifs, and the information is also widely leveraged for regulatory network construction in methods like SCENIC+ and CellOracle [74, 71]. As an explicit statistical framework, D-SPIN can also incorporate prior information on motif-binding information to prioritize such interactions (STAR Methods 1.5). Independent of the single-cell datasets, the motif-binding network has better correspondence with ChIP-seq datasets in the early accuracy rate than all network inference methods. Motif-binding network has a lower AUROC because it does not make predictions on edges with no known motifs, while AUROC evaluates predictions on all edges. After incorporating the motif-binding prior information, both CellOracle and D-SPIN exhibited substantial accuracy rate improvement in accuracy rate, while D-SPIN still achieved the highest correspondence with ChIP-seq data. The early accurate rate of D-SPIN achieved 11 to 15, indicating that the top-identified interactions by D-SPIN are highly reliable (Figure 3A).

### Program-level regulatory network model of genome-wide Perturb-seq data

Interpreting gene-level networks with hundreds to thousands of nodes can be complex. For interpretability, we constructed a regulatory network model with D-SPIN at the gene program level to investigate the global architecture and information processing logic of the regulatory network implied by the perturbation responses (STAR Methods 2.3). Transcriptome-scale gene expression changes under gene knockdown reflect how cells coordinate the activities of major cellular functions to maintain homeostasis under stress. To reveal the global architecture of perturbation responses, we focused on 3,136 knockdown perturbations that are each associated with more than 10 differentially expressed genes (DEGs) and 20 cells (Figure 3B, STAR Methods 2.1). We coarse-grained the transcriptional profile into 30 gene programs, a number informed by both the Bayesian information criterion (BIC) and the elbow method, to balance the model’s representative power and complexity (Figure S3B, STAR Methods 1.11) [75].

The extracted gene programs reflect both general cell biology and lineage-specific gene programs for K562 cells (Figure 3C, Figure S3A, SI Table 3). We annotated the gene programs with a combination of bioinformatic databases, including DAVID, Enrichr, and manual lookup [66, 67, 68]. The gene programs contain genes involved in core cellular processes such as transcription, translation, RNA processing, and mitosis. There are also lineage-specific programs, including an erythroid-specific program with hemoglobin (HBG1, HBG2, HGZ) and glycophorin (GYPA, GYPB, GYPE) genes, as well as two myeloid-associated programs with phagosome/actin-organization (ACTB, ACTG1, ARPC3) and immune-response (LAPTM5, VASP, RAC2) genes, respectively, which agrees with the multi-lineage potential of K562 cells.

D-SPIN generated a program-level regulatory network model that provides a wiring diagram of K562 internal regulations and responses to gene knockdown perturbations. The inferred network model is robust against sampling noise, and more than 98% of edges remain structurally stable when resampled from near-optimal solutions of the network inference problem (Figure S3C, STAR Methods 1.7). The D-SPIN network model contains a set of sub-networks or modules of programs that are enriched with positive interactions (Figure 3D, Figure S3D). Using the Leiden community detection algorithm [76] (STAR Methods 2.12), we automatically decomposed the D-SPIN network into seven core cellular functions, including transcription, translation, protein degradation, and cell-cycle-related functions.

The network contains a set of negative interactions between gene programs that are expressed in distinct cellular states. The strongest negative interaction is between gene programs expressed at different stages of mitosis. For example, the P29 Spindle microtubule has negative interactions with both P25 DNA replication and P27 Histone. The strong interactions inside the cell cycle sub-network of P25-P30 are able to reconstruct the transcription state distribution during cell cycle progression (Figures S3E and S3F). We also observe negative interactions between P4 Erythroid and P6 Phagosome, which is consistent with the presence of two mutually exclusive differentiation paths that lead to erythroid and myeloid cell fates.

In addition to the core K562 regulatory network, D-SPIN also inferred interactions between each gene program and each gene knockdown perturbation (Figure 3C and Figure S3G). Similar to the grouping of sub-network modules, we classified knockdown perturbations into a set of 40 perturbation groups that we refer to as G1-G40 (Guide RNA group 1 through 40) with unsupervised Leiden clustering of the perturbation response vectors (STAR Methods 2.12, SI Table 3). Gene knockdowns within the same guide cluster have similar perturbation responses, suggesting that these genes are involved in the same pathway or have potential interactions.

Identified clusters reflect the pathway-level organization in the K562 cell, revealing both well-known cell biology and more novel or cryptic organization of pathways (Figure 3E). Globally, each guide clusters are enriched for specific cell-biological functions, including DNA replication, the MAPK signaling pathway, and RNA degradation. As an example of interactions between pathways, we found components of the proteasome core particle (20S proteasome) are grouped into cluster G3 with genes involved in protein transport, including AKIRIN2 and IPO9 (Figure 3E). In recent studies, AKIRIN2 was found to bind directly to fully assembled 20S proteasomes to mediate their nuclear import with the nuclear import factor IPO9 [69]. Similarly, the D-SPIN guide clusters group the gene C7orf26 with the integrator subunits INTS10, INTS13, and INTS14, a key result of the original Perturb-seq study [29].

### D-SPIN network identifies key regulators of erythroid and myeloid differentiation

The program-level network provides a global view of regulatory architecture, but regulations are ultimately executed by specific genes. Identifying these key regulators can motivate experimental validations and help pinpoint potential therapeutic targets. Therefore, we also designed D-SPIN to infer interactions between gene programs and candidate regulators such as transcription factors, providing a finer-grained perspective of gene regulations (STAR Methods 2.3).

K562 cells have the potential to differentiate into erythroid or myeloid lineages. Traditional approaches to finding cell fate controllers often examine the phenotype measurements of knockdown experiments. In Perturb-seq, knocking down a fate regulator may alter the marker gene expression of the target fate or the opposite fate. However, the knockdown may also not produce an obvious phenotype due to the robustness of fate-control networks. By integrating all perturbation information into a unified network model, D-SPIN provides a more comprehensive way of tackling such problems. For comparison, we extracted an erythroid and a myeloid program from the original Perturb-seq study and analyzed potential regulators with both differential expression (DE) and D-SPIN network models [29].

The D-SPIN network uncovers many more well-known regulators of the two cell fates compared with the standard DE analysis. D-SPIN discovered erythroid fate regulators KLF1, NFE2, GFI1B, and GATA1, while DE only found GATA1; D-SPIN discovered myeloid fate regulators SPI1 (PU.1) and MEF2C [77], while DE found none of them (Figures 3F and 3G, SI Table 3, SI Data). The improvement is due to D-SPIN constructing the model with all perturbation conditions together, while DE individually examines each perturbation condition with fewer than a few hundred cells. Constructing the D-SPIN model on each perturbation separately shows that perturbing other genes also contains information about the top regulators SPI1 and KLF1, and the most informative perturbations are not knocking down these genes directly, but knocking down epigenetic regulators such as KDM5A and BRD8 (Figure S3H). Interestingly, D-SPIN also uniquely identified NPM1, which inhibits both cell fates. Recent research shows that in K562 cells, BCR-ABL1 kinase activates NPM1 to relocate KLF1 and SPI1 proteins from the nucleus to the cytoplasm, therefore silencing their cell fate control capabilities [78]. The relocation of KLF1 and SPI1 potentially explains why their knockdowns have little effect on the phenotype of cell fate program expressions.

The striking contrast between the regulators identified by D-SPIN and those indicated by traditional DE analysis underscores the advantage of constructing a unified regulatory network model. D-SPIN discovered the cell-type-specific (BCR-ABL1-dependent) post-transcriptional differentiation inhibition effect of NPM1, which cannot be found with methods using TF binding motif analysis. By constructing a network model, D-SPIN can uncover hidden regulatory interactions in single-cell perturbation datasets that are missed by traditional analyses.

### Coarse-grained D-SPIN models provide insight into global perturbation response strategies

Cells coordinate many internal processes to maintain homeostasis in response to damage and perturbation, while the corresponding control strategies remain poorly understood due to the lack of global models. The program-level D-SPIN model allows us to analyze the distributed regulation of core cellular processes in response to damage induced by gene knockdown.

While the program-level network is compact, the model can be simplified further through an automated coarse-graining strategy based on graph clustering (Figure 4A). Gene programs are grouped into sets that we call modules, and gene perturbations are grouped into major categories. We term the strategy network coarse-graining, similar to statistical physics, where correlated degrees of freedom are grouped into a single variable to aid computation and interpretation. The coarse-graining strategy produces a minimal network with 4 distinct patterns of program activation and inhibition under perturbations, which we call stress-response strategies (STAR Methods 2.13).

**Figure 4:**
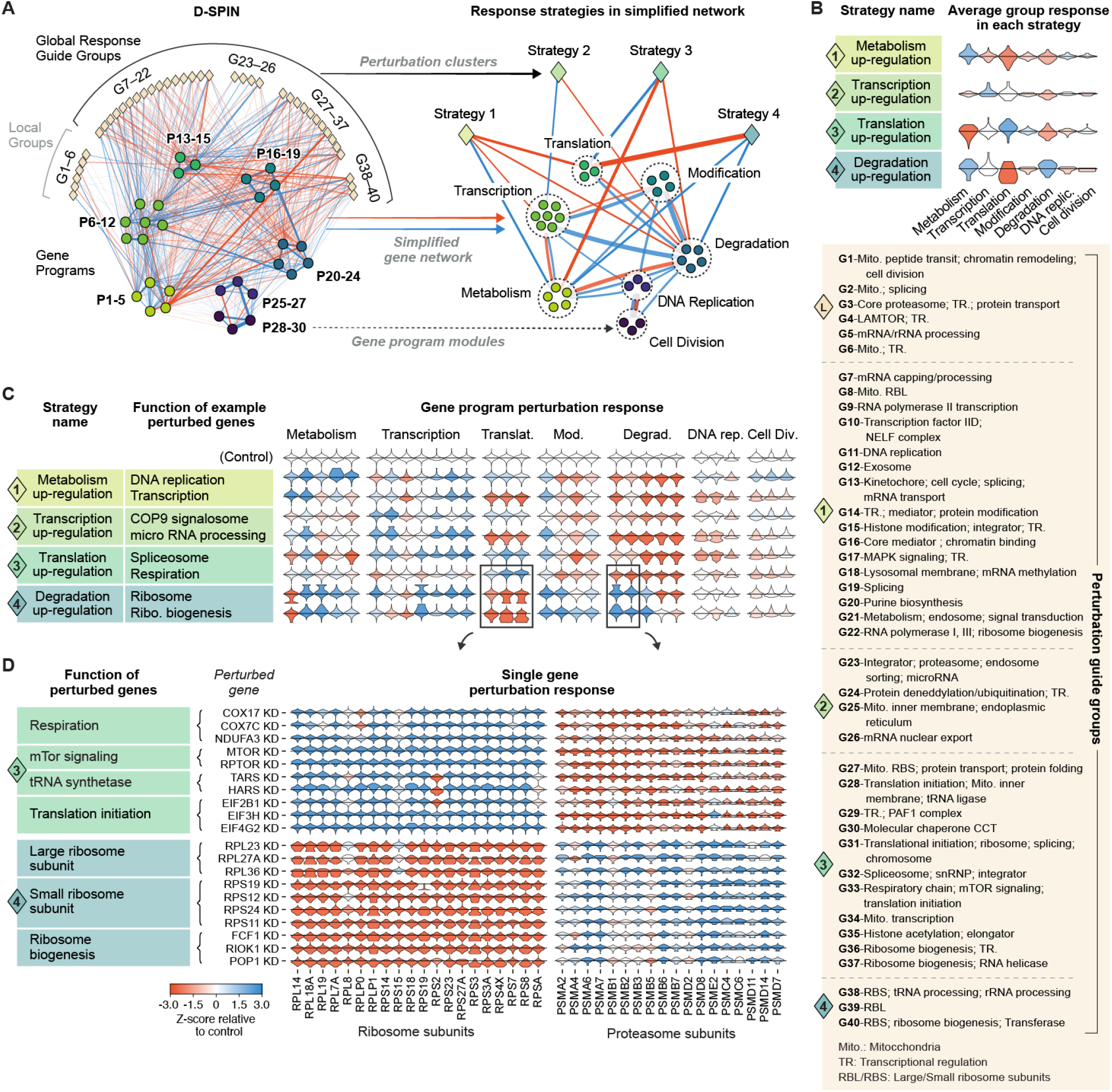
D-SPIN network model identifies global perturbation response strategies in K562 cells for distinct classes of gene knockdowns. (A) Diagram of network coarse-graining by grouping gene programs into 7 identified gene program modules and grouping guide RNA groups into 4 identified strategies. The resulting coarse-grained model enables the global analysis of cellular regulatory responses. (B) Violin plots of averaged perturbation response vectors on programs in guide groups in each response strategy. K562 cells contain 4 distinct classes of global response strategies, and we name each strategy by the upregulated characteristic biological function. (C) Violin plots show gene program expression under knockdown of example pathway components relative to the control samples treated by non-targeting guide RNAs. The upregulated biological function is typically distinct from the genes being perturbed. For example, when an RNA polymerase subunit is knocked down, the cell upregulates metabolism while downregulating translation and degradation; when a ribosome subunit is knocked down, cells upregulate protein degradation and metabolism while downregulating translation. (D) Violin plots show the single-cell gene expression distribution for ribosome and proteasome subunits under knockdown of example genes in translation upregulation and degradation upregulation strategies relative to control samples. The program-level response strategies are reflected as coherent expression changes at the single-gene level. For example, when respiration-related genes COX17 or COX7C are knocked down, all ribosome subunit genes are upregulated, and proteasome subunit genes are downregulated.

The D-SPIN-identified response strategies suggest non-trivial forms of homeostasis regulation. We found that each strategy features the upregulation of a major biological function and the downregulation of other functions, except for the fourth strategy, which upregulates both metabolism and degradation (Figure 4B). The four global response strategies are named by the characteristic program upregulation as metabolism upregulation, transcription upregulation, translation upregulation, and degradation upregulation. In general, K562 cells may be capable of mounting a wide range of homeostatic response strategies, but D-SPIN identified the four highlighted strategies under the specific context of Perturb-seq experiments.

The regulatory strategies provide insight into the information flow in the cell. The compensatory functions are typically not directly associated with the gene knockdown, suggesting long-range regulation or coupling between seemingly distinct processes (Figure 4C). For example, a large number of metabolism upregulation responses were induced by perturbations to transcription processes, including RNA polymerase(G9, G22), Transcription factor IID (G10), and mediator (G16). Consistent with the observation, transcription stress has been shown to induce an elevated ATP pool and rewired metabolism states [79]. The knockdown of mTOR signaling components leads to translation upregulation responses, reflecting the importance of mTOR in coordinating protein synthesis and energy utilization [80]. Furthermore, disruptions to different sub-processes of the same cellular process can lead to distinct response strategies. For example, perturbations to translation initiation factors (G28) lead to the upregulation of translation-related genes and the downregulation of protein degradation. Perturbations to ribosome subunits and ribosomal RNA (G28, G39, G40) induce the downregulation of translation-related genes and the upregulation of protein degradation. At the single-gene level, the response strategies are reflected as the coherent up- or downregulation of genes involved in the corresponding gene programs (Figure 4D).

These different response strategies indicate that the cell mitigates the loss of a gene by upregulating a potentially compensating cellular function. These upregulations suggest the existence of active regulatory feedback within the network to connect distinct cellular processes for homeostasis maintenance. The D-SPIN model organizes thousands of perturbations and million-level single cells into classes of regulatory strategies, providing insights into principles of information processing and homeostatic control.

### Modeling immunomodulatory drug responses in primary human immune cells

Understanding how unique cell types in an interacting community respond to a given drug or other therapeutic interventions is essential for effective drug development. Specifically, immunomodulatory drugs vary in the breadth and specificity of their biochemical targets, and a key question is to understand how differences in biochemical preferences translate into transcriptional changes, and thus differences in therapeutic responses [81, 82, 83].

We therefore developed an experimental platform for large-scale profiling of human immune cell drug responses and analyzed the single-cell data with D-SPIN. Our experimental system was designed to characterize T cell-driven hyper-activation of the immune system, as is observed in auto-immune and hyperinflammatory states. We cultured a heterogeneous population of primary donor-derived peripheral blood mononucleated cells (PBMCs) that contained T cells, B cells, myeloid cells, and NK cells. We specifically activated the T cells with anti-CD3/CD28 antibody chimera (Figure 5A, STAR Methods 1.12), which led to the dynamics of immune activation of the entire cell population (Figure 5B, Figures S4A-S4D, STAR Methods 1.14).

**Figure 5:**
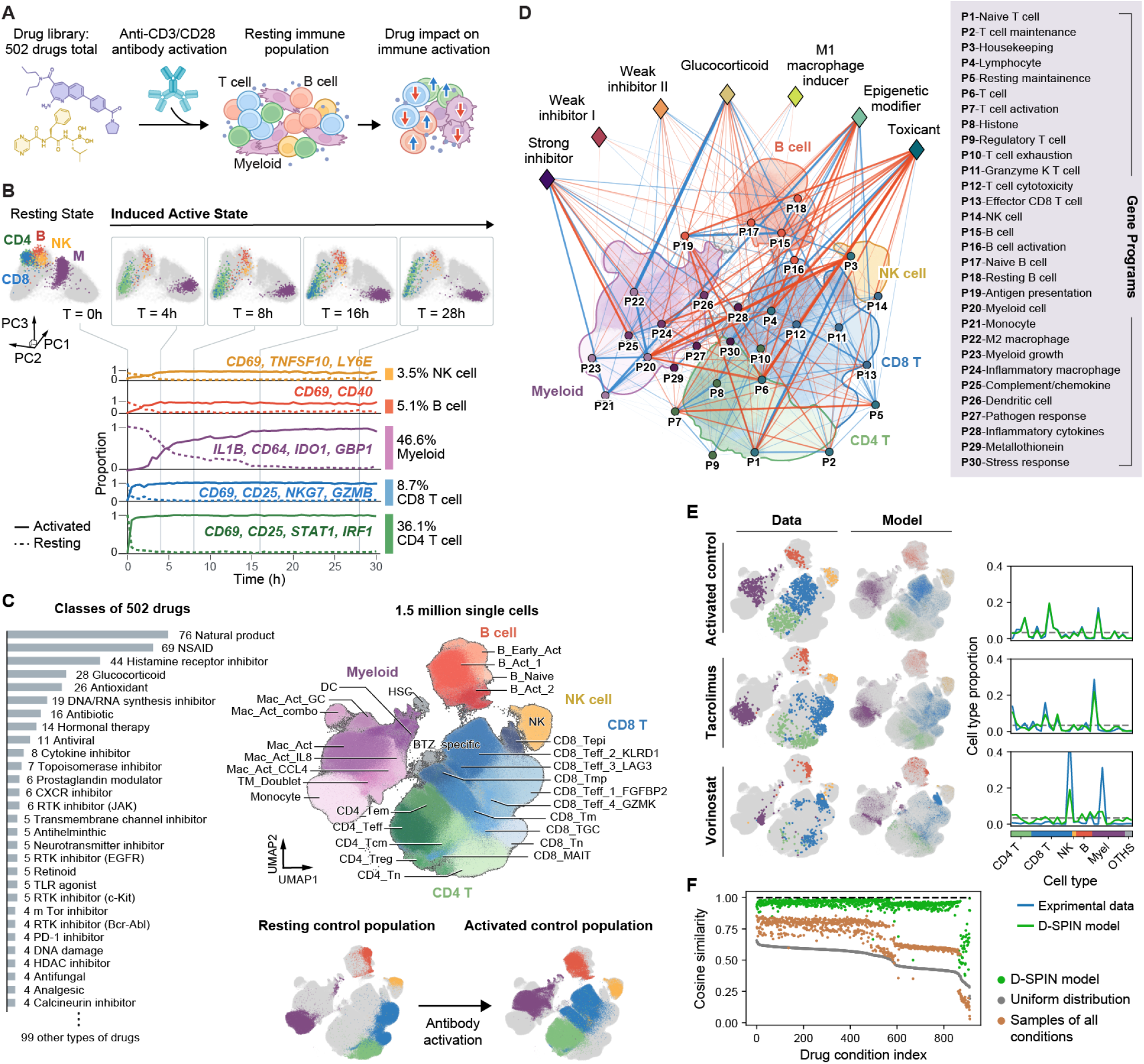
D-SPIN derives a drug-response network model from human immunomodulatory drug-response single-cell mRNA-seq profiling experiments. (A) Schematic of the experiment design for profiling drug responses on T cell-mediated immune activation. Peripheral blood mononuclear cells (PBMCs) were harvested from a human donor. The cell population was treated with anti-CD3/CD28 antibodies that specifically activate T cells and immunomodulatory drugs drawn from a library. The cell population was profiled after 24 hours of drug and antibody treatment. (B) (top) PCA projections derived from a 30-hour time-course experiment of T cell-mediated immune activation, where samples were taken every 30 minutes for single-cell mRNA-seq. (bottom) Time courses of the proportion of activated and resting cells in each cell type, and example activation gene markers. T cells reach activated states first in 2 hours, and myeloid cell activation lasts 16 hours. (C) (left) Histogram of profiled drug classes shows a variety of biochemical properties and target pathways. (right-top) UMAP embedding of 1.5 million filtered single cells obtained from the drug profiling experiments. In the profiled cell population, we identified 32 cell states in the major cell types of T, B, NK, and myeloid cells, and each cell state is curated by marker genes and gene differential expression. (right-bottom) UMAP embedding of the resting control cell population (without antibody activation) and the activated cell population. The resting and activated cell states compose the major partition on the UMAP. (D) D-SPIN-inferred regulatory network model between 30 gene programs (P1-P30, circles), as well as interactions between programs and 7 drug classes (diamonds) identified through clustering the response vectors. In the network rendering, each program is positioned on the UMAP at the cell state where it is most highly expressed. (right box) Gene programs are functionally annotated through gene ontology annotation tools and manual lookup [66, 67, 68]. (E) (left) UMAP embeddings of experimental cell state distribution and the state distribution generated by the D-SPIN model in activated control, tacrolimus treatment, and vorinostat treatment. (right) Line plots quantify the cell state distributions of experimental data and D-SPIN models. The dashed line is the uniform distribution for reference. D-SPIN models closely match the control and tacrolimus-treated samples. The model fits less well in the vorinostat-treated sample but still captures the overall distribution pattern. (F) Scatter plots show cosine similarity between experimental data and cell-state distributions generated by D-SPIN, compared with two reference null models, uniform distribution and cell type abundance by pooling all cells together. D-SPIN models have higher than 0.9 cosine similarities for 92.4% of conditions.

To profile the action of different immunomodulatory drugs on the immune activation process, we selected 502 small molecules and collected a total of 1.5 million filtered single cells. The drug library contains a diverse set of small molecules targeting pathways, including mTOR, MAPK, glucocorticoids, JAK/STAT, and histone deacetylases (HDAC) (Figure 5C, SI Data). Therapeutically, the library contains drugs used for treating autoimmune diseases (e.g., tacrolimus, budesonide, tofacitinib) as well as FDA-approved anti-cancer drugs (e.g., bosutinib, crizotinib). Drugs were added together with the activating antibody or alone to profile their effect on the resting cell population. In total, we profiled 1.5 million filtered single cells in resting and activated conditions, with over 1,200 total conditions and 31 different immune cell states, including 4 CD4 T cell states, 10 CD8 T cell states, an NK cell state, 4 B cell states, and 8 myeloid cell states. (Figure 5C, Figure S4E, STAR Methods 2.2, SI Table 5).

We constructed a program-level regulatory network with D-SPIN to dissect how small molecules interact with the network to create the altered cell population states (STAR Methods 2.4). Although the cell population contains various immune cell types and drug-specific cell states, D-SPIN took all single cells to construct a unified regulatory network model that captures all the cell types and cell states by pairwise interaction between gene programs and drug impacts on the programs. We coarse-grained the transcriptional profile into 30 gene programs, a number informed by both the BIC and the elbow method (Figures S5A-S5C, STAR Methods 1.11).

Among the discovered gene programs, there are global cell-type programs such as P6 T cells, P15 B cells, P14 NK cells, and P20 myeloid cells, as well as specific cell-state programs, including those corresponding to T cell resting and activations, anti-inflammatory M2 macrophage, and pathogen-responding M1 macrophages (Figure S5B). The program function was annotated by a combination of informatics databases and manual lookup [66, 67, 68] and subsequently validated on a single-cell atlas of human immune cells [84].

Similar to the K562 Perturb-seq response network, the drug response network has a distinct modular structure, with a set of core network modules composed of tightly interacting gene programs (Figure 5D, Figures S5D-S5F). Each module represents a group of programs that are expressed together in a cell type or subtype in the population. For example, a module containing P11 Granzyme K T cell, P12 T cell cytotoxicity, P13 Effector CD8 T cell, and P14 NK cell corresponds to the population of CD8 T cells and NK cells. Negative interactions primarily occur between programs expressed in different cell types, such as P2 T cell maintenance and P19 Antigen presentation, as antigen presentation happens in B cells and myeloid cells but not T cells (Figure 5D).

As a minimal probabilistic model, D-SPIN can simulate the drug-modulated cell state distribution with only 465 network interaction parameters and 30 parameters for each treatment condition with high fidelity. Qualitatively, we found visual similarity between D-SPIN simulated cell state distributions on the UMAP embedding and actual cell states. Quantitatively, the cosine similarity between the model and data distribution is higher than 90% for 92.4% of the samples (Figure 5E). The drugs with lower model accuracy tended to be drugs that drove the cell population into a few highly specialized states, such as the proteasome inhibitor bortezomib, where D-SPIN models only have qualitative agreement with the cell state distribution. Furthermore, D-SPIN, as a parsimonious maximum entropy model, is mathematically constrained to avoid overfitting and promote generalization [85]. We trained a separate D-SPIN model with 50% of cells in each sample and evaluated the quantitative agreement between D-SPIN-simulated cell states and data distribution in the remaining 50% of cells. D-SPIN also achieved good accuracy under various metrics, including error of mean and cross-correlation, optimal transport distance, and probabilities of most frequent states (Figures S5G and S5H, STAR Methods 2.14).

### The program-level network organizes drugs into phenotypic classes

While many of the drugs in our experiments have been previously reported as immunosuppressive, less is known about their specific effect at the transcriptome level. Our single-cell profiling and D-SPIN analysis provided a single integrated platform to compare the transcriptomic effects of drugs with different chemical mechanisms of action. In addition to the network model, D-SPIN inferred perturbation response vectors that quantify interactions between each drug and the regulatory network. We identified 70 out of 502 drugs that had statistically significant interactions with the D-SPIN-inferred network compared with control experiments (STAR Methods 2.12). D-SPIN grouped these effective drugs into 7 classes, including strong inhibitor, weak inhibitor I, weak inhibitor II, glucocorticoid, M1 macrophage inducer, epigenetic regulator, and toxicant (Figure 6A, Figures S6A and S6B).

**Figure 6:**
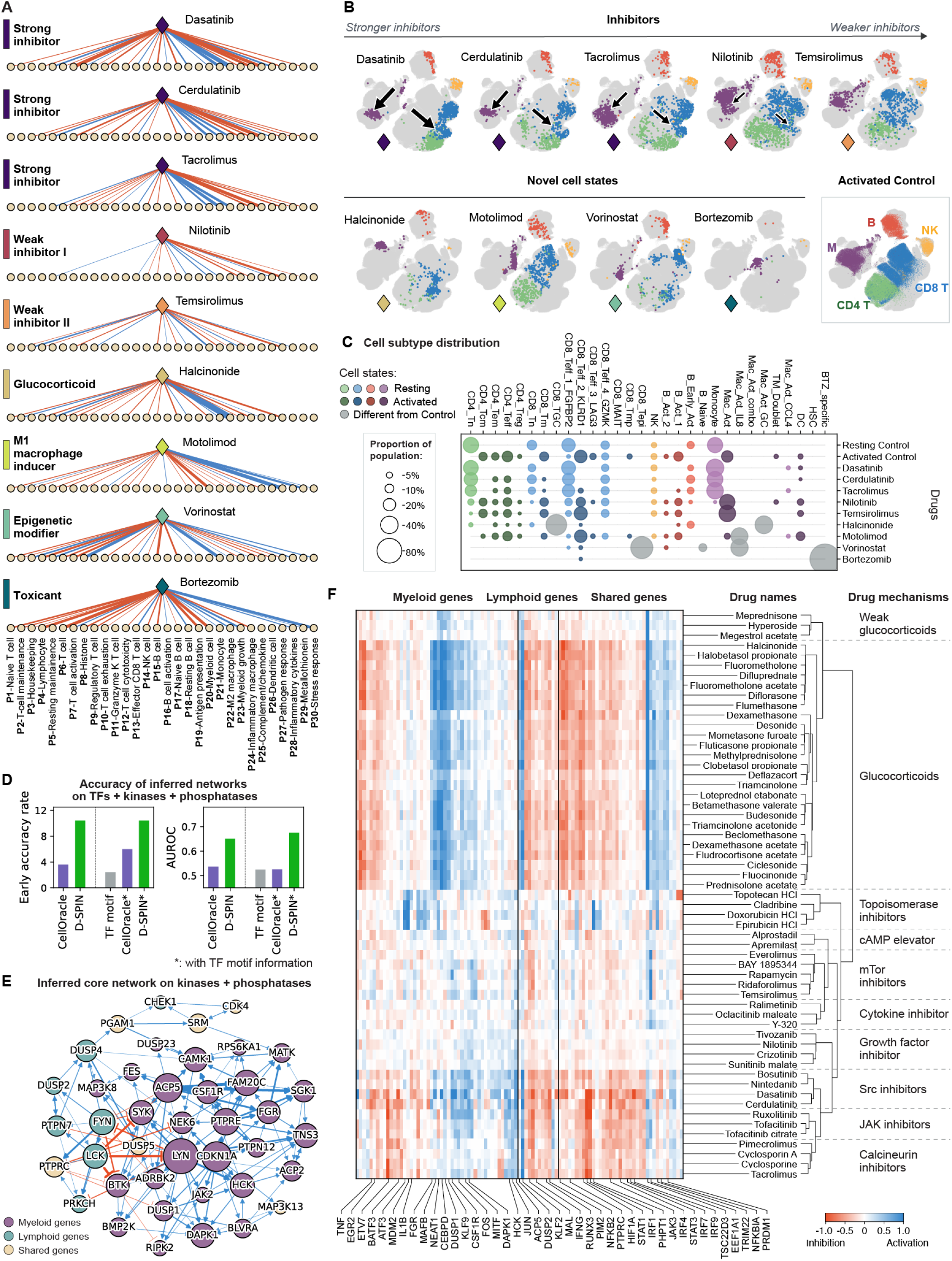
Drug classification derived from D-SPIN aligns with known drug targets and mechanisms of action. (A) Diagrams of response vectors inferred by D-SPIN are shown as positive/negative interactions between the drug and gene programs for drug examples. D-SPIN identifies 7 phenotypical classes of drugs based on their response vectors. (B) UMAP embeddings of the cell population after treatment with antibody and example drugs from different classes. The top row shows 5 immune inhibitors with decreasing strength from left to right. The bottom row shows 4 example drugs that each induce novel cell states distinct from both the resting and activated control cell populations. Activated control population (bottom-right) is shown for comparison. (C) Bubble plots show cell-subtype distributions induced by selected drugs and the control population. The bubble sizes scale with population proportion, and are colored by major cell types and resting/activated classification. With decreasing inhibitor strength, the proportion of the activated immune-cell population (deep colors) gradually increases. Some drugs induce cell states that are different from both resting and activated control populations. (D) Bar plots quantify the early accuracy rate and AUROC of the correspondence between the inferred regulatory networks from the drug profiling dataset and protein-protein interactions in the String-db databases [68]. Methods with the (*) symbol utilize TF-motif binding information as prior knowledge in the network inference. (E) Rendering of the subnetwork of D-SPIN-identified regulators in kinases and phosphatases. The node size is proportional to the number of identified interactions. The subnetwork is partitioned into two groups of genes that are primarily expressed in myeloid cells or lymphoid cells. (F) Heatmap of D-SPIN-inferred gene-level responses for drugs with immune inhibitory effects. The dendrogram of hierarchical clustering shows that the gene-level signatures classify drugs into groups with similar molecular mechanisms of action.

Drugs within any one of these seven classes induced similar shifts in cell population structures. For example, three of the drug classes include drugs that inhibit T cell, B cell, and myeloid cell activation. Both UMAP visualization and cell state distributions indicate that these inhibitors act with a spectrum of strengths (Figures 6B and 6C, Figure S6C, STAR Methods 2.15). Very strong inhibitors, including the cancer drug dasatinib and immunosuppressive drug tacrolimus, completely block the immune cell activation and result in a cell population similar to unstimulated PBMCs. Weak inhibitors, such as temsirolimus, only slightly increase the proportions of cells in the resting states.

Beyond inhibitors, D-SPIN identified a drug class of glucocorticoids (GCs), which are steroid-derived small molecules that activate the glucocorticoid receptor (Figures 6A and 6B). The GC class includes well-known immunosuppressive drugs, including halcinonide, budesonide, triamcinolone, and dexamethasone. GCs suppress immune activation but also generate cell populations different from the strong inhibitors, in particular, distinct myeloid cell states as visualized on the UMAP. On the program level, D-SPIN showed GCs more weakly suppress the chemokine secretion program (P25) than strong inhibitors but more strongly induce a program P22 associated with M2 macrophages, including expression of CD163 [86, 87].

D-SPIN also identified a drug group that induces the activation of inflammatory, pathogen-responsive M1 macrophages (Figures 6A and 6B). The class includes activators of Toll-like receptors (TLRs) in macrophages, which sense the presence of pathogens and induce innate immune responses. The M1 macrophage inducer class contains TLR7 agonists (vesatolimod, resiquimod) and TLR8 agonists (motolimod, resiquimod), and produces macrophage states related to host defense that highly express P27 Pathogen response and P29 Metallothionein programs.

The other two classes of drugs, epigenetic modifiers and toxicants, are associated with stress response programs (Figures 6A and 6B). Toxicants include the proteasome inhibitor bortezomib, a potent non-selective histone deacetylase (HDAC) inhibitor, panobinostat, and DNA topoisomerase inhibitor 10-hydroxycamptothecin. Toxicants strongly activate the P30 Stress response and have mostly inhibitory interactions with other gene programs, especially generic cell-type programs such as P6 T cell and P20 Myeloid cell. The epigenetic-modifiers class consists of HDAC inhibitors, which generate an epigenetically disrupted T cell state (CD8 T epi.) that has elevated expression of histone component genes and DNA topoisomerase TOP2A. All together, these data identify classes of drugs by the unique downstream programs that they impact.

### Gene-level network reveals signaling hubs and response signatures of different drug targets

Phenotypical classification reveals a spectrum of inhibitors with different strengths, and these inhibitors employ distinct mechanisms. The majority of these inhibitors have known primary targets, such as tyrosine kinases (TKs) for dasatinib and Janus kinases (JAKs) for cerdulatinib, as well as extensive biochemical assays on the alternative targets and binding strengths of these inhibitors [88, 89]. However, it remains an important question of how the differences in biochemical properties, including target specificity and breadth, translate into different transcriptional responses in the context of a heterogeneous interacting cell community, and ultimately lead to distinct clinical outcomes. Understanding these varying transcriptional responses would be valuable for developing more effective therapeutic strategies.

To identify gene-level signatures of drug actions and gain a global view of the regulatory network controlling T cell-mediated immune activation, we constructed gene-level network models with D-SPIN (STAR Methods 2.4). During immune activation, regulatory interactions are executed by both DNA-binding interactions of TFs as well as signaling transduction pathways of (de)phosphorylation mediated by kinases (phosphatases). We selected 657 highly expressed regulatory genes for the network construction, including 388 TFs, 187 kinases, and 71 phosphatases [90, 91]. We assessed the network inference quality with the physical protein-protein interaction networks in the String-db database, which integrated multiple data sources, including databases of interaction experiments, curated complexes/pathways, and literature text-mining [68]. We also evaluated CellOracle for comparison, as other inference methods, such as PIDC, GENIE3, and GRN-Boost2, do not scale to datasets with millions of cells [71]. D-SPIN achieved significantly higher accuracy rates compared to CellOracle, both without and with prior knowledge of TF-motif binding information (Figure 6D, STAR Methods 2.11). Notably, in the accuracy benchmarking of the K562 Perturb-seq dataset, incorporating motif binding information significantly boosted the accuracy of both D-SPIN and CellOracle. However, in the context of immune activation, where signal transduction pathways play a key role, the binding information only improved the accuracy by a small margin. The result demonstrates the unique advantages of D-SPIN as a data-driven, perturbation-based inference method, compared to motif-based methods like CellOracle or SCENIC+ [71, 74].

Among the hundreds of kinases and phosphatases, D-SPIN identified 41 genes with regulatory roles, highlighting key signaling hubs during immune activation (Figure 6E). The top hub genes with the most interactions are primarily Src-family protein tyrosine kinases, including LYN, FYN, HCK, SYK, LCK, BTK, and FGR. Src-family kinases are major components in the immune signaling pathways [92]. For example, FYN and LCK are among the first activated molecules downstream of the T cell receptor. Accordingly, inhibitors of Src-family kinases, such as dasatinib and bosutinib, were among the strongest inhibitors observed in our experiments. The identified regulatory genes also include multiple phosphatases from the dual-specificity phosphatase (DUSP) gene family, which contribute to regulating the intensity of immune response by controlling MAPK signaling [93]. Globally, the inferred core network is partitioned into two groups: genes primarily expressed in myeloid cells and genes expressed in lymphoid cells. Inhibitory interactions between the two groups indicate that these genes do not coexist in the same cell state.

The gene-level responses inferred by D-SPIN distinguish inhibitors with different biochemical mechanisms of action. Hierarchical clustering of the responses organizes the inhibitors into three major categories: strong inhibitors, weak inhibitors, and glucocorticoids (GCs) (Figure 6F). The strong and weak inhibitors are further split into subclasses that match the primary targets of these small-molecule drugs. The targets of strong inhibitors include Src family kinases (bosutinib, cerdulatinib, dasatinib), JAKs (tofacitinib, rux-olitinib), and calcineurin (cyclosporine, tacrolimus). The targets of weak inhibitors include growth factors (crizotinib, sunitinib, nilotinib), mTOR (everolimus, temsirolimus, sirolimus), cytokine (ralimetinib), cAMP (alprostadil), and topoisomerase (topotecan HCl, doxorubicin HCl). Broadly, the analysis suggests that inhibition can be achieved via a range of distinct biochemical pathways and mechanisms. The strong inhibitors in our data target molecules immediately downstream of T cell receptor activation, such as the Src family kinase LCK.

Furthermore, the fine-grained D-SPIN model highlights gene signatures that separate drugs with different mechanisms (Figure 6F). Compared with strong inhibitors, the anti-inflammatory GC drugs selectively activated KLF9, TSC22D3, MAFB, NEAT1, and DUSP1/2. Each of these genes participates in M2 macrophage polarization or signal transduction of glucocorticoid receptors [94, 95, 96, 97, 98, 99]. In comparison, strong inhibitors had increased repression of STAT1/3, JAK3, and IRF1/4/7/9. JAK-STAT signaling and the IRF family are both central pathways during inflammation [81, 100], whose inhibition corresponds to the shutdown of immune activation by these strong inhibitors. The distinctions between strong inhibitor types are more subtle. A group of responding genes exhibits increased response strength from calcineurin inhibitors to JAK inhibitors to Src inhibitors, including CSF1R, ACP5, JAK3, and DAPK1. The strongest inhibitors, Src inhibitors, have additional suppression of inflammation-associated genes, such as NFKB1A, IL1B, and EGR2.

Weak inhibitors with different mechanisms each had unique response signatures on a few genes (Figure 6F). mTOR inhibitors such as rapamycin induced the activation of EEF1A1, an upstream regulator of the PI3K/AKT/mTOR pathway [101]. The activation also suggests potential compensatory mechanisms on EEF1A1 under mTOR inhibition. Topoisomerase inhibitors specifically activated a few genes associated with the p53 pathway, including ATF3, MDM2, and PHPT1 [102, 103, 104]. p53 plays a vital role in maintaining genome stability, and p53 deficiency is known to sensitize cells to topoisomerase inhibitors [105]. Both ATF3 and MDM2 have been shown to participate in the DNA damage stress response induced by topoisomerase inhibitors [106, 103].

### Drug combinations generate novel cell states with hyper-suppression

Immunomodulatory drugs are often used in combinations. Understanding how specific drug combinations can tune the transcriptional cell-state distribution of the immune cell population could enable more precise drug regimens to meet therapeutic goals. However, the design of drug combinations at the transcriptome scale is challenging due to the large number of potential drug-gene interactions. D-SPIN models provide a framework to compare the action of individual drugs on both the gene level and program level in the context of the regulatory network to identify drug combinations with potentially useful therapeutic applications.

Therefore, we applied D-SPIN to interpret the mechanisms of combinatorial drug action. We selected 10 drugs from different drug classes identified by D-SPIN and profiled all pairwise combinations experimentally (Figure S7A). We found that 83% of the drug interactions in our profiling were additive or sub-additive on the gene program level, meaning that the effect of the drug combination is equal to, or weaker than, the sum of single-drug effects. Other types of drug interactions include dominant, synergistic, and antagonistic (STAR Methods 2.16). The additive interactions between drugs recruited a combination of transcriptional programs from single drugs, creating novel cell states or population states, especially between drugs that have distinct impacts.

Among the combinations, glucocorticoids (GCs) and strong inhibitors induced coherent anti-inflammatory effects on gene programs. However, their combination produced a novel macrophage state that was distinct from the state produced by either drug alone (Figure 7A). To investigate how the drug combination generated novel biological outcomes, we performed single-cell mRNA-seq on cell populations treated with the GC halcinonide in combination with the Src inhibitor dasatinib across a range of doses and analyzed the com-binatorial response with D-SPIN (Figure S7B). Both halcinonide and dasatinib are anti-inflammatory drugs that suppress the activation of lymphoid and myeloid cell populations, and in the D-SPIN model, halcinonide and dasatinib activate or inhibit the same set of gene programs (Figure 7B). However, the two drugs acted on specific gene programs with different intensities (Figure 7C). For example, compared with dasatinib, halcinonide induced weaker suppression of genes involved in macrophage activation, such as IDO1, CD40, and SLAMF7, in the P24 inflammatory macrophage program. Halcinonide also promoted the M2 polarization of macrophages, strongly activating M2-associated genes, including CD163, MS4A6A, and VSIG4 in the P22 M2 macrophage program.

**Figure 7:**
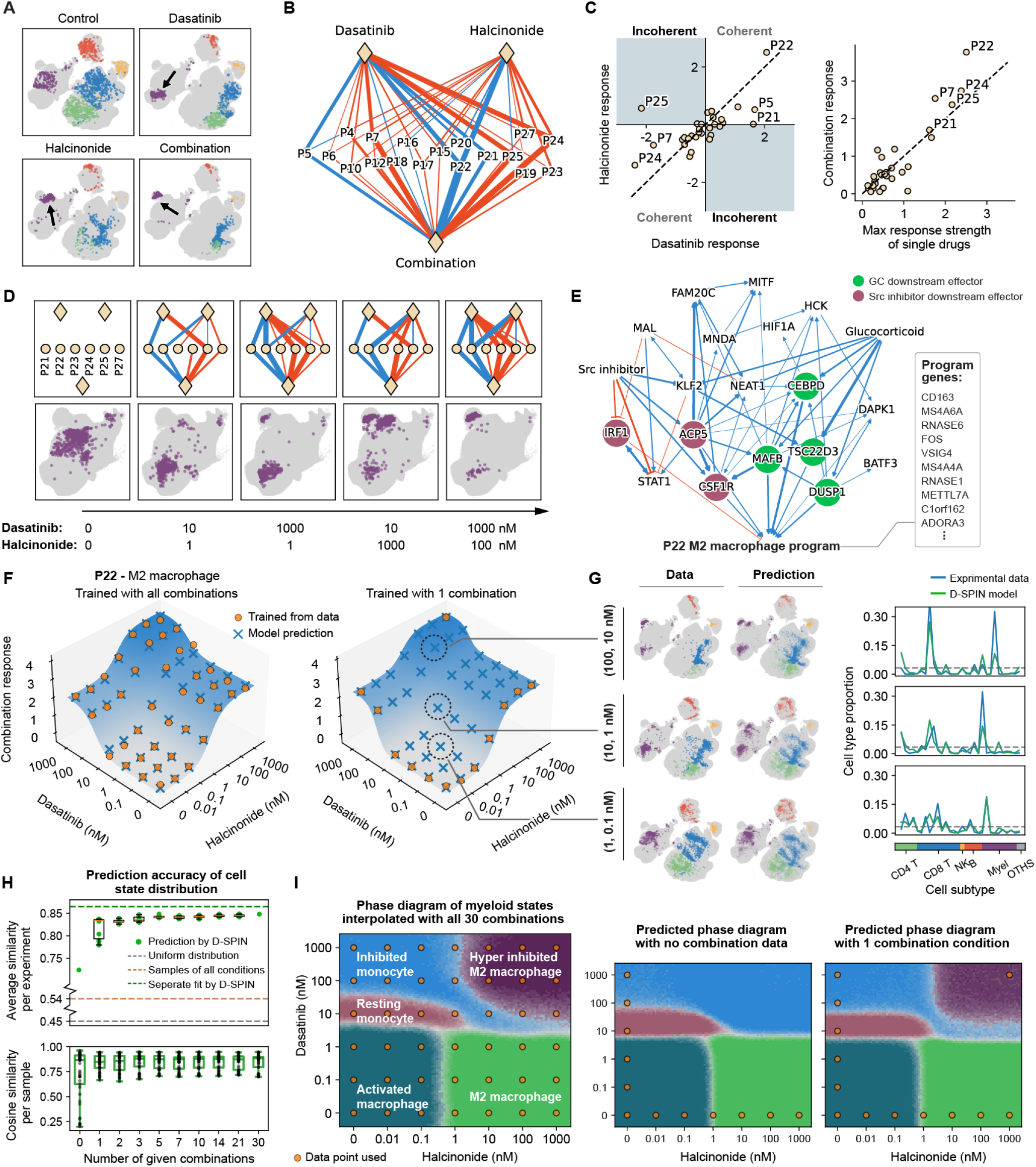
D-SPIN reveals network-level mechanisms of combinatorial drug action and predicts dosage combination response. (A) UMAP embedding of activated control, dasatinib, halcinonide, and drug combination-treated cell population, with arrows indicating different myeloid cell state changes. The drug combination induces a novel macrophage state. (B) Diagram of response vectors for dasatinib and halcinonide alone and their combination. The single drugs activate/inhibit the same set of gene programs, and the combination drug response is the superposition of single drug effects. (C)(left) Scatter plots compare the response vectors on each gene program for dasatinib and halcinonide. The two drugs activate/inhibit the same set of gene programs, but with different strengths. (right) The drug combination responses are plotted against the stronger single-drug response for each gene program. The combination responses are generally similar to or higher than the maximum of the single-drug responses. (D) (top) Diagram of response vectors for two single drugs and their combination on impacted myeloid programs under different example dosages. (bottom) UMAP embeddings of myeloid states for each drug combination dosage. The novel macrophage state is induced by the combinatorial gene program recruitment of the two drugs. (E) Rendering of the subnetwork of Src inhibitor and glucocorticoid responses, as well as inferred regulators of the P22 M2 macrophage program. D-SPIN nominates downstream effectors that mediate the effect of P22 activations, which are highlighted by colored circles. (F) Surface plot shows response vectors on program P22 under different dosage combinations. The D-SPIN inferred combinatorial response follows the sum of two sigmoid-shaped single-drug responses with a multiplicative interaction term, allowing prediction of dosage combination responses with only single-drug data and one drug combination condition. (G) D-SPIN predicts cell state distributions for unseen dosage combinations, as shown in (left) UMAP embeddings and (right) line plots of cell state distribution comparisons between experimental data and D-SPIN-predicted cell state distributions. (H) Cell state distribution prediction error of D-SPIN with different numbers of drug combination conditions used for training, shown as (top) average cosine similarity for all samples across 10 different sets of drug combination training data and (bottom) cosine similarity for each sample in an example set of drug combinations. (I) D-SPIN predicts the phase diagram of myeloid states under different drug dosage combinations of dasatinib and halcinonide with only one drug combination sample. Observing one combination sample is necessary, as the D-SPIN model with no combination data misclassifies the region of hyper-inhibited M2 macrophage state.

We found that the novel macrophage state was induced by the additive recruitment of gene programs by the two drugs, including augmented activation of the M2 macrophage program and hyper-repression of the macrophage activation. The strengths of responses were generally larger than either drug alone but did not exceed their sum (Figure 7C). The two drugs had coherent effects on gene programs across the entire dosage range, and the strength of action increased with drug dosages (Figure 7D). Given that both drugs were profiled at maximally effective doses, the enhanced response of the drug combination indicates pathway-level cooperation between the two drugs rather than additive dosing. The drug combination recruited a combination of gene programs in a dose-dependent fashion, which was also observed in a previous study performed at the single-protein level [107]. Conceptually, by combining the two drugs, we effectively fine-tuned the transcriptional states of the macrophage population by exploiting the additivity of their effects (Figure 7D).

Furthermore, the gene-level regulatory network model revealed that Src inhibitors and GC activate the M2 program through distinct gene regulators, therefore explaining why their effects are additive. We used D-SPIN to specifically analyze the regulators of the M2 macrophage program and extracted the core subnetwork together with the responding genes of Src inhibitors and GCs (Figure 7E). The core subnetwork nominated candidate gene regulators that directly mediate the response of the M2 program by each drug type. We found that the effects of Src inhibitors and GCs are mediated by distinct groups of regulatory genes. For GCs, the response is mediated by the activation of TSC22D3, DUSP1, CEBPD, and MAFB, all of which are known to be associated with glucocorticoid receptor signaling and M2 polarization [95, 98, 108, 96]. For Src inhibitors, the response is mediated by the inhibition of IRF1 and activation of CSF1R and ACP5. IRF1 is a key controller of M1 polarization, and its inhibition would bias macrophages towards the M2 state [109]. CSF1R and ACP5 are also found participating in M2 polarization, especially in the context of tumor-associated macrophages (TAMs) in the cancer microenvironment [110, 111]. Thus, both Src inhibitors and GCs activate the M2 macrophage program, but their responses are mediated by two distinct sets of regulators and pathways. Their distinct mechanisms provide the opportunity to manipulate a spectrum of macrophage states by fine-tuning the dosage of the two-drug combinations.

### D-SPIN predicts drug dosage combination responses with a single combination experiment

The generative D-SPIN model enabled us to predict how transcriptional programs respond to drug dosage combinations by only observing a subset of treatment conditions. The inferred single-drug response vectors exhibit a sigmoid-shaped response curve with the logarithmic dosage (Figure 7F). While most combination responses are qualitatively additive or sub-additive, some programs show substantial deviation from additivity (Figure S7C). Thus, measurements of single-drug responses alone are insufficient predict combinatorial response, and at least some combinatorial treatment conditions are required to quantify the interactions between drugs.

However, while single-drug dosage response data alone are insufficient, we found that D-SPIN only requires one extra drug combination condition to generate quantitatively accurate dosage combination response predictions. For each program, we predict the response with an additive sigmoid (sgm) model with a multiplicative interaction: sgm_1_(*c*_1_) + sgm_2_(*c*_2_) + *γ* sgm_1_(*c*_2_) sgm_2_(*c*_2_), where *c*_1_, *c*_2_ are the log-dosage of the two drugs and *γ* is the interaction strength of the program. Using measurements of single-drug treatment and one combination at saturating dosages, D-SPIN learns the interaction strength and predicts the program responses at unseen dosage combinations (Figure 7F, Figures S7C and S7D, STAR Methods 2.17). Moreover, these predictions on program-level drug responses can be combined with the inferred regulatory network to predict cell state distributions. The regulatory network trained with single drugs plus one combination accurately predicts the cell state distribution of unseen dosage combinations, as shown by the agreement between predicted distribution on UMAP embedding and cell type distributions with experimental data (Figure 7G).

The quality of D-SPIN prediction improves with increasing number of dosage combinations used for training, but the dominant information gain occurs from zero to one combination condition (Figure 7H). Introducing just one combination increases the average cosine similarity between experimental and predicted cell state distributions from 0.72 to 0.84. Without any combination data, many dosage combinations are poorly predicted with cosine similarities below 0.5; with one drug combination condition, predictions for most combinations exceed 0.7 cosine similarity. Nonetheless, adding more combinations yields only modest additional improvements, indicating that the additive sigmoid model with interaction well characterizes the program response landscape. Additional prediction accuracy metrics, such as program mean, correlation, and optimal transport, shows similar trend, reinforcing that the critical information is the interaction strengths between the two drugs, which can be identified from a single combination experiment (Figure S7E).

Furthermore, we used the generative D-SPIN model to construct a phase diagram of myeloid cell states under any dosage combination between dasatinib and halcinonide by using the model trained with only one/three drug combination samples. We computed the predicted program expression distribution of myeloid states under unseen dosage combinations and compared the distribution with the target myeloid population to determine the myeloid state labels (Figure 7I, STAR Methods 2.18). In the phase diagram learned with all combination data, at a low dosage of dasatinib, the myeloid state transitions from the activated macrophage to the M2 macrophage with increased halcinonide dosages. At higher dosages of dasatinib, increasing halcinonide dosage produces resting monocyte, inhibited monocyte, and hyper-inhibited M2 states. Overall, manipulating the drug doses allowed a smooth conversion of the macrophage population between different states. In comparison, the model trained with single-drug data also captures the majority of cell state transition boundaries, but fails to identify the hyper-inhibited M2 macrophage state. In contrast, including a single combination condition is sufficient for D-SPIN to qualitatively recover the full phase diagram (Figure 7I). With three combination conditions, the predicted phase diagram boundaries closely match the model trained with all available dosage combinations (Figure S7F). Together, these results show that D-SPIN can extrapolate from single-drug data with a minimal number of combination experiments to accurately reconstruct the landscape of dosage combination response at both gene program and cell state levels.

## Discussion

Here, we introduce D-SPIN, a generalizable and interpretable framework that can be applied to study the perturbation response of cells, including genetic perturbations, small molecules, and other signaling conditions. D-SPIN constructs quantitative, generative models of gene regulatory networks by integrating information from perturbation conditions. The mathematical structure of D-SPIN allowed us to develop a computationally efficient, parallel inference procedure that can be run on hundreds of CPU cores to perform network inference on datasets with thousands of perturbations and millions of cells. Single-cell mRNA-seq methods enable large-scale perturbation response studies across cell types and organisms. D-SPIN provides the framework to integrate such data into regulatory network models that can be analyzed and compared to reveal the architecture, logic, and evolution of these networks across species and time.

A major goal in biology is to control the distribution of cell states in a cell population. Being able to generate a specific set of progeny from stem cells or modulate the state of the immune system has tremendous implications for treating cancer and autoimmune diseases. We showed that D-SPIN identified network-level mechanisms of cell-state modulation by TF perturbations in synthetic HSC networks (Figure 2G) and nominated key regulators of erythroid-myeloid fate choices in K562 perturb-seq datasets (Figures 3F and 3G). Further, D-SPIN revealed the phase diagram of transitioning between macrophage states by modulating the dosage combination of small molecule drugs (Figure 7G), and the effector genes that mediate the additive drug response on the M2 program (Figure 7H). Together, these results suggest that D-SPIN could be used to design interventions that precisely tune networks and cell states.

For drug combinations, we demonstrated that Src inhibitors and glucocorticoids created a novel hyper-inhibited M2 state by additively recruiting a set of gene programs. The novel cell state encompasses complicated expression-level changes of numerous single genes, but at the regulatory network level, the cell state neatly originates from the superposition of single-drug effects. By focusing on the regulatory networks, D-SPIN enables us to dissect these combinatorial mechanisms and interpret them through the inferred D-SPIN regulatory network and responses, and nominate specific regulators that mediate the observed program-level responses. The results provide a conceptual framework for interpreting and predicting the effect of drug combinations. D-SPIN further predicts cell state distribution under dosage combination treatment with only single-drug data and one combination condition. Profiling the dosage response of single drugs and only a few combinations could serve as a strategy to systematically depict the combination response landscape for therapeutic objectives. Additivity could arise from the modularity of gene regulatory circuits, such that different pathways impact gene expression levels independently. While this principle has been studied on a small scale for a small number of drugs [107], our results suggest that such principles of superposition might hold at the transcriptome scale. Further work is needed to reveal the specific conditions where additivity holds or breaks down.

Cells are distributed control systems that modulate many internal processes to maintain homeostasis and execute cell-fate transitions. However, the principles of distributed control at the transcriptome scale are poorly understood. With D-SPIN, we showed that in K562 cells, perturbations triggered system-wide response strategies where distinct cellular processes are coordinated with long-range information flow. For example, knocking down translation initiation factors or mTOR pathway genes induces coherent down/up-regulation of proteasome/ribosome subunits, pointing to the presence of sensing and controlling mechanisms in the cell. The long-range coordination between pathways might be general, as we also observed in the drug profiling experiments that mTOR inhibitors like rapamycin activated EEF1A1, a key factor in translation. Identifying these feedback control points with combinatorial knockdown screening may lead to the discovery of new regulators or regulating mechanisms.

Our work on D-SPIN represents a significant advancement in constructing biologically interpretable predictive models of regulatory networks underlying cellular decision-making. The transparent graphical model architecture naturally captures interactions between genes and can be interpreted as pathways, circuits, or networks that delineate the flow of information. In the meantime, the model can generate the full distribution of transcriptional states in cell populations across perturbation conditions. While low-dimensional embedding visualizations such as UMAP have become a common practice of single-cell analysis to overview the cell type proportion and shifts, D-SPIN constructs an interpretable circuit/pathway type of model that uncovers internal regulatory connections controlling these shifts of cell population in large-scale perturbation profiling. The simplicity of D-SPIN’s formulation originates from the maximum entropy principle [85].

Toward the ultimate goal of building a cell-scale “digital twin” model of the cell, deep neural network models, especially transformer models, have been gaining popularity [8, 9, 10]. Similar to D-SPIN, these deep models can integrate information from large datasets and make predictions with accuracies that improve with more training data. However, in contrast to D-SPIN, their complex architectures and immense number of parameters provide limited mechanistic insight. D-SPIN directly identifies biological pathways and regulators from perturbation-based single-cell gene expression data. This interpretability has enabled us to uncover candidate regulators of K562 cell fate selection, as well as identify drug response pathways in myeloid cells that generate novel cell states under combinatorial drug conditions. Importantly, D-SPIN achieves this without requiring additional data sources such as ATAC-seq, making it particularly valuable as perturbation-based data becomes increasingly abundant through scaled condition-based barcoding strategies [29, 33].

D-SPIN’s connection with spin network models provides insights into the fundamental nature of cellular regulation. The spin network model is an equilibrium model used to study physical systems at or near thermal equilibrium. Theoretically, D-SPIN depicts the cell population as a collection of points residing in an energy landscape that can be tilted by the perturbation vector to shift the distribution of cell states in a cell population. Energy-landscape models represent a highly simplified class of dynamical systems, as their behavior can be captured within a single energy (potential) function. Equilibrium spin network models have been used to study a much broader range of systems that are far from thermal equilibrium, including neural networks and bird flocks [39, 40, 38]. However, it remains unclear why equilibrium models can yield such significant predictive power for strongly non-equilibrium systems. The ability of equilibrium models to produce low-error reconstructions of cell population gene-expression states suggests that cells, in certain situations, may be effectively modeled as equilibrium systems driven through various configurations by a biasing drive. Such models have been demonstrated for cell-fate regulation and might represent a simplifying principle [5]. The ability to model a cell as an equilibrium system driven through different states presents a powerful simplification, as also explored in other chemical systems, offering a potential path toward more global theories of gene regulation [112, 113].

### Limitations of the Study

The D-SPIN framework has several current limitations that represent targets for extension in future work. First, for simplicity and interpretability, D-SPIN only considers pairwise interactions between genes or gene programs in the network. These interactions correspond to the second-order terms in the energy function. The inclusion of higher-order multi-body interactions or concentration-dependent regulatory responses would enhance accuracy and predictive capabilities [114, 115, 116, 117]. Second, D-SPIN is an equilibrium model and does not account for the dynamics of the system. Spin network models have natural extensions that incorporate dynamics [118, 119, 120]. Incorporating temporal dynamics would allow the simulation of directed networks, as directed network models in general cannot define consistent stationary distributions when the network contains feedback loops [59, 58, 60]. Third, D-SPIN assumes that the interactions within the core regulatory network ***J*** are not altered by the perturbations, which is a reasonable approximation in perturbation response scenarios such as gene knockdown or small molecule action. However, in scenarios such as cellular differentiation or disease progression, the regulatory network may undergo changes under different conditions due to shifts in epigenetic regulation. Epigenetic reorganization can be included in future versions of D-SPIN by allowing interactions encoded in ***J*** to also be condition-dependent.

## Author Contributions

Conceptualization, J.J., S.C., D.A.S., Z.J.G., M.T.; Methodology, J.J., D.A.S., M.T.; Data Analysis, J.J., S.C., C.S.M., J.H., M.T.; Experimental Investigation, S.C., T.T., C.S.M., T.K., J.H.P., J.V., E.D.C.; Data Curation, C.S.M., T.K., Q.Z., J.H., E.D.C.; Resources, S.C., T.T., C.S.M., T.K., Q.Z., J.V., J.H., E.D.C., Z.J.G., M.T.; Writing, J.J., S.C., D.A.S., M.T.; Visualization, J.J., S.C., I.S., M.T.; Software, J.J., Y.G.; Supervision, J.J., S.C., Z.J.G., M.T.; Funding Acquisition, Z.J.G., M.T.

## Supplementary Tables

Supplementary tables are available at Caltech Research Data Repository https://doi.org/10.22002/1pxgr-eqa61

## Data and code availability

Gene counts and metadata of drug profiling experiments are available at Caltech Research Data Repository https://doi.org/10.22002/2cjss-wgh69

Supplementary data of the analyses are available at Caltech Research Data Repository https://doi.org/10.22002/tbmca-wbj97

MATLAB, Python implementations and Jupyter Notebook demonstrations of D-SPIN are available on GitHub https://github.com/JialongJiang/DSPIN

## Acknowledgements

We would like to acknowledge the NIH (TR01 GM150125, R01HD100039), the Heritage Medical Research Institute, Charles Trimble, the Shurl and Kay Curci Foundation, the Merkin Institute for Translational Research, 10x Genomics, Amgen, and Chan Zuckerberg Initiative. The single-cell profiling experiments were performed at the Beckman Institute Single-cell Profiling and Engineering Center (SPEC). We acknowledge the Protein Expression Center at the Beckman Institute of Caltech for the use of the liquid-handling robot. Sequencing was performed at the UCSF CAT, supported by UCSF PBBR, RRP IMIA, and NIH 1S10OD028511-01 grants. We acknowledge Dr. Guy Riddihough, Dr. Ariane Helou, and Dr. Jenna Sternberg for editorial assistance with the manuscript. DAS acknowledges the support of an NSERC Discovery Grant and a Tier-II Canada Research Chair. We acknowledge Rob Phillips, Eric Siggia, and Venkat Chandrasekaran for insightful scientific discussions. JH acknowledges the support of NIH 5R01GM135337 grants.

## STAR Methods

## Supplementary Figures

**Figure S1:**
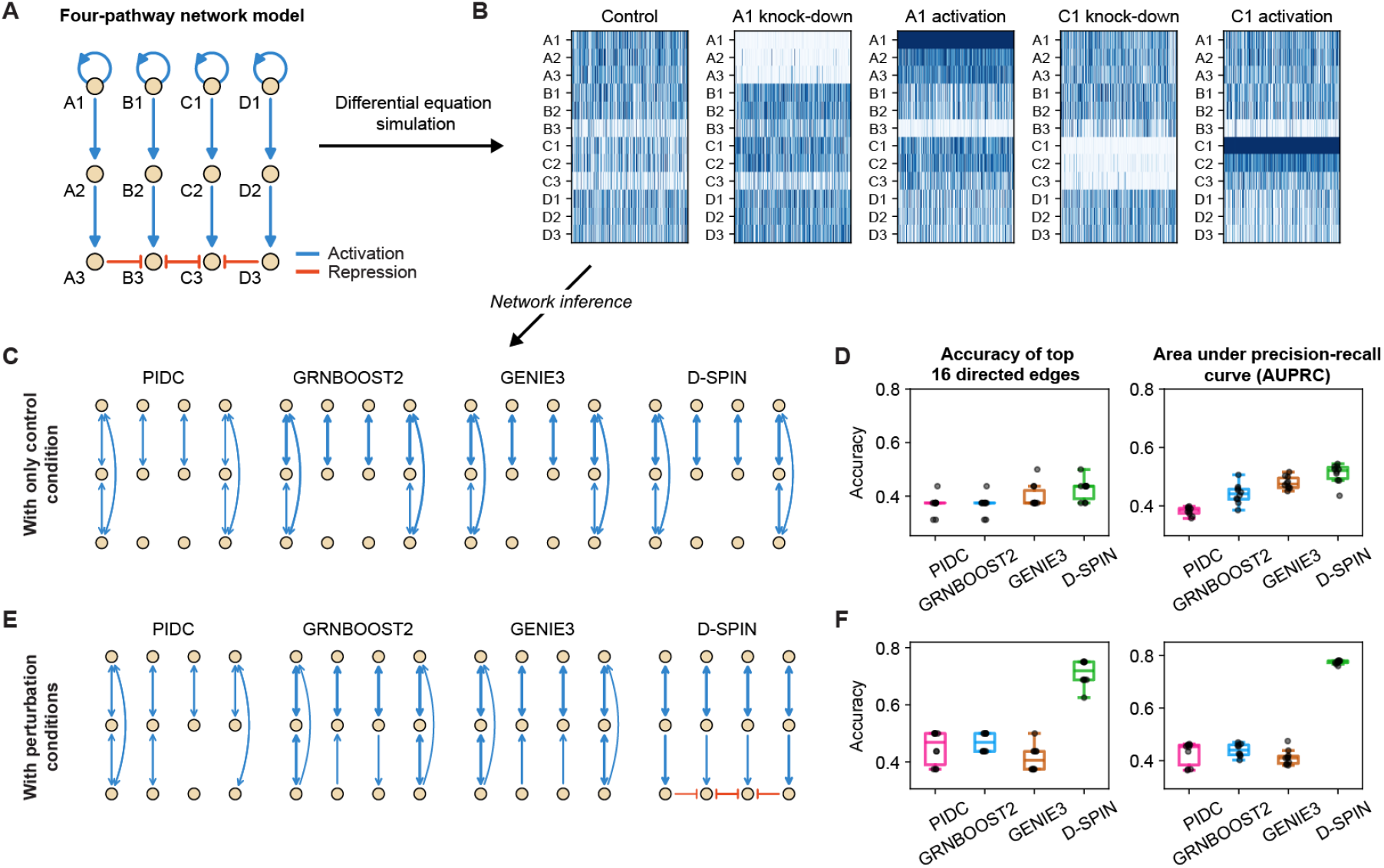
D-SPIN recovers hidden interactions by integrating information from perturbations in an example four-pathway model; related to Figure 1. (A) Structures of a four-pathway circuit model where the inhibitory interactions between A3-B3, B3-C3, and D3-C3 are redundant to inhibit B3 and C3, thus difficult to discover. (B) Examples of simulated gene expression profile heatmaps from the network across a series of single-gene knockdown and activation conditions. (C) The network diagrams show the top 16 directed edges of inferred regulatory networks by D-SPIN and other state-of-the-art methods with only control conditions. Without perturbations, all methods fail to discover the interactions associated with B3 and C3, as they are significantly repressed by other genes. (D) Box plots quantify the accuracy of the top 16 inferred interactions and AUPRC for each method. With only control data, D-SPIN has slightly better accuracy than other methods. (E) Network diagrams of the top 16 directed edges of inferred regulatory networks with perturbation data. D-SPIN successfully recovers all hidden interactions, while other methods do not identify the inhibitory interactions associated with B3 and C3. (F) Box plots quantify the accuracy of the top 16 inferred interactions and AUPRC for each method. With perturbation data, D-SPIN achieves significantly higher accuracy than other methods.

**Figure S2:**
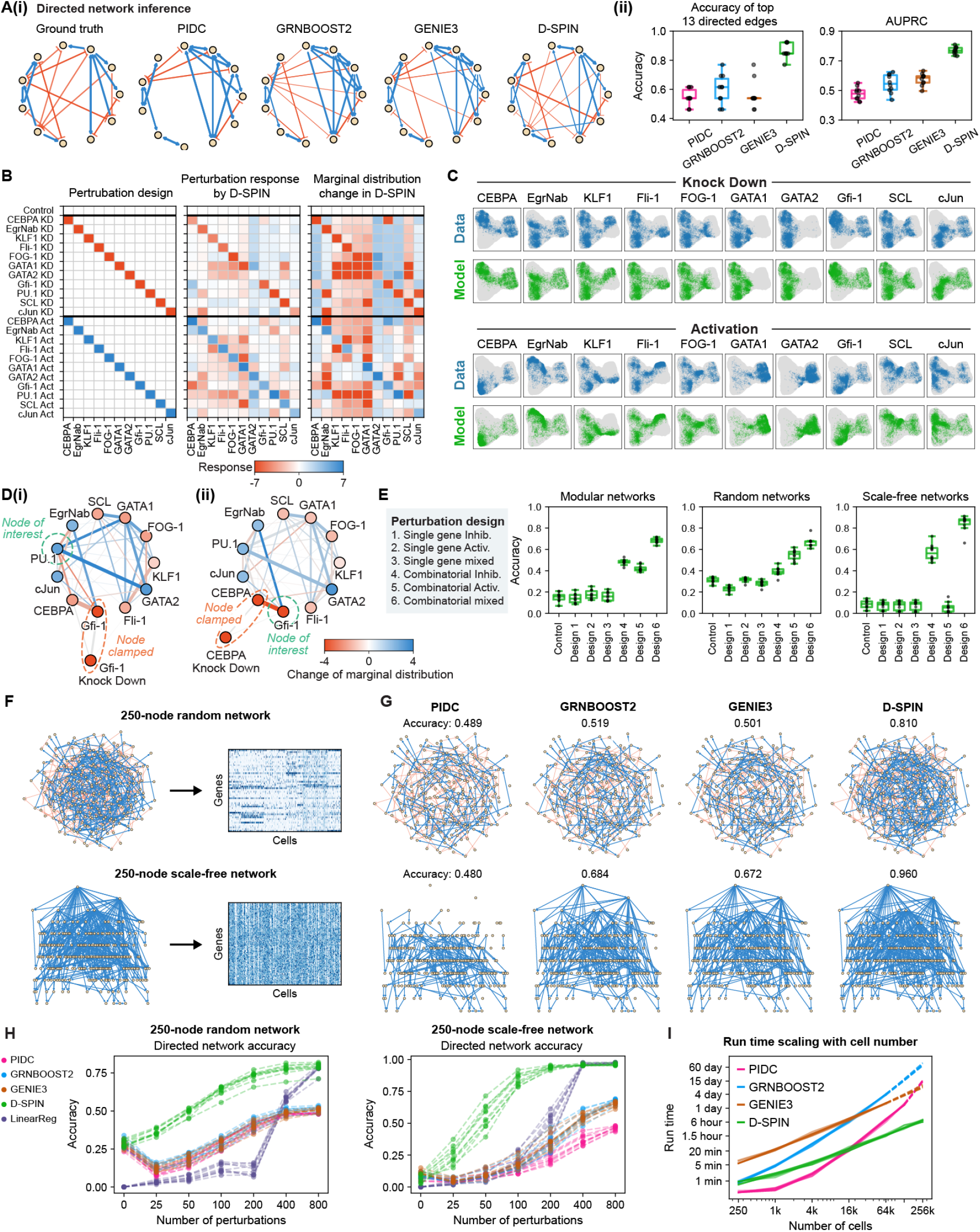
D-SPIN model recapitulates cell state distributions and allows reasoning through the network; D-SPIN achieves high accuracy in large-scale directed network inference benchmarking; related to Figure 2. (A)(i) Diagrams of the ground truth directed network and inferred directed networks by D-SPIN and other state-of-the-art methods. (ii) The box plots quantify the accuracy of the top 13 inferred directed edges (the true network has 26 directed edges) and AUPRC of the inferred networks. (B) Heatmaps indicating (left) applied perturbations to the network; (middle) perturbation response inferred by D-SPIN; (right) marginal distribution changes of single genes as detailed in Sec. 2.7. D-SPIN response vectors identify the applied perturbations and quantify their impact on single genes. The major discrepancy of inferred perturbation occurs on GATA2 knockdown, potentially because GATA2 is a transient regulator in the network. (C) The D-SPIN model accurately generates cell-state distributions with the regulatory network and perturbation response vectors, as visualized by UMAP embeddings of simulated single-cell data by BEELINE (blue) and state distribution generated by the D-SPIN model (green) across all single-gene knockdown/activation perturbations. (D) Network diagrams visualize edge sensitivity analysis in D-SPIN, showing how each network edge positively (blue) or negatively (red) contributes to the marginal distribution change of the gene of interest under a clamped perturbation, as detailed in Sec. 2.7. Network nodes are colored by the upregulation or down-regulation of their activities defined by marginal distributions, and edges are colored by their contribution to the activity of the node of interest. For example, panel (i) shows that under Gfi-1 knockdown, the upregulation of PU.1 is significantly positively contributed by the edge EgrNab-Gfi-1. (E) Box plots quantify the directed network inference accuracy by D-SPIN on 125-node networks with different topologies under 100 perturbations from different perturbation design strategies. Combinatorial random perturbation with both activating and inhibiting perturbations is most effective among the tested strategies. (F) (top) Network diagram of an example 250-node Erdős–Rényi (ER) network with equal probability of activating (blue) and inhibiting (red) edges. Heatmap showing simulated single-cell data for evaluating the accuracy and scalability of D-SPIN as detailed in Sec. 2.8. (bottom) Network diagram and simulated single-cell data for an example 250-node scale-free network. (G) Diagrams show the inferred directed network of different methods as the subnetwork of correctly inferred edges for the example ER network (top) and scale-free network (bottom). D-SPIN achieves higher accuracy compared with other methods. (H) The directed network inference accuracies are plotted with the number of perturbations in 10 randomly generated ER networks (left) and scale-free networks (right). The accuracy of D-SPIN continuously increases with the perturbation number. The linear regression model has low accuracy with a small number of perturbations, but achieves high accuracy only with a large number of perturbations. (I) The running time of different network inference methods as a function of cell number in a 250-node ER network. D-SPIN only takes a few hours for 256,000 cells on 2 CPU cores, while other methods require weeks. Dashed lines are interpolations, as detailed in Sec. 2.9.

**Figure S3:**
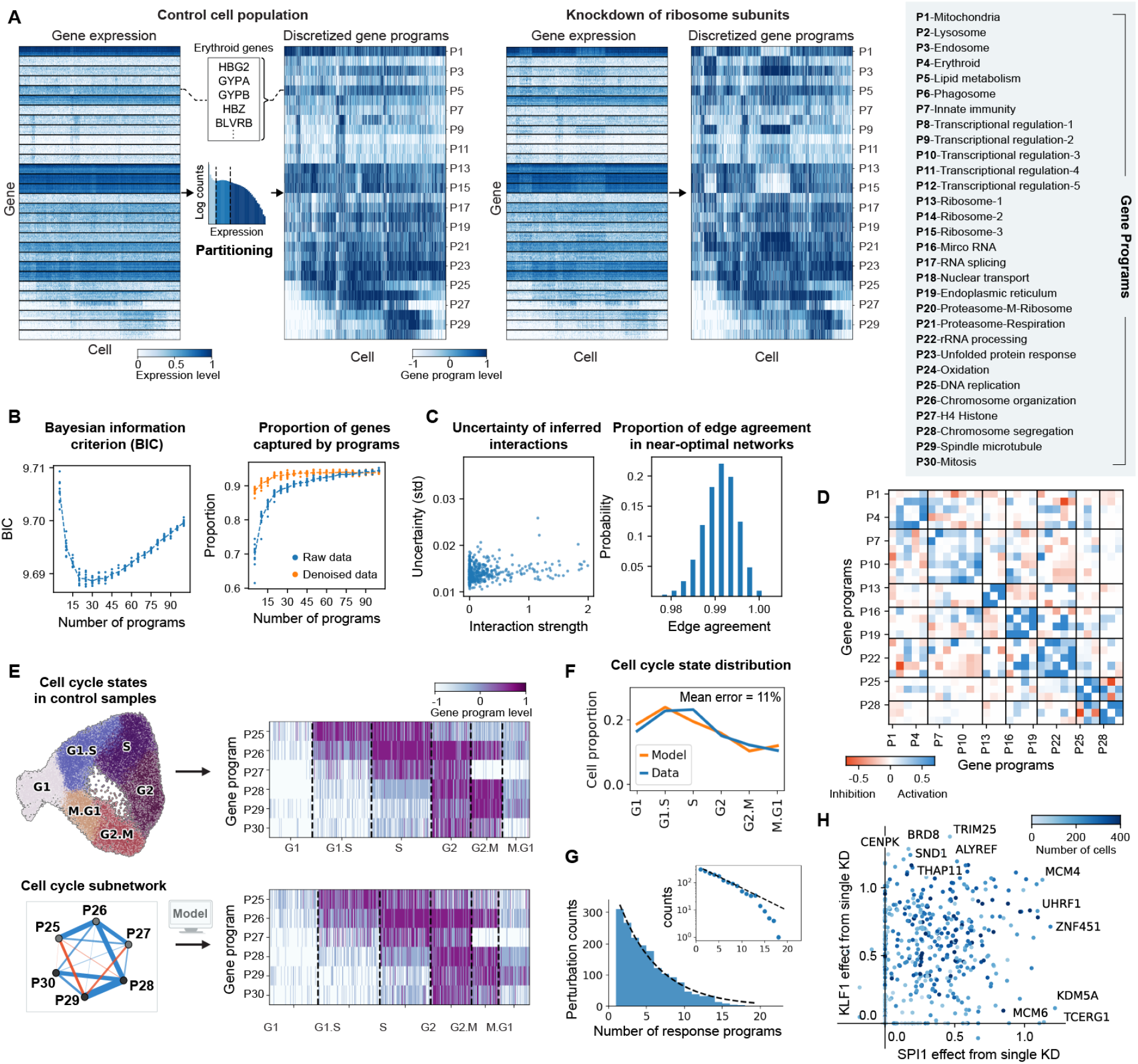
D-SPIN decomposes gene expression into programs and reconstructs cell cycle states; related to Figure 3. (A) Heatmaps of gene expression and discretized gene program level for (left) control samples of non-targeting guide RNAs and (right) knockdown of the large or small ribosome subunits. The discretized gene programs characterize and denoise major expression patterns in the gene matrix. (B)(i) Bayesian information criterion (BIC) of the gene program decomposition as a function of gene program number. 30 is the optimal program number that minimizes BIC when the effective sample size is 20k cells. (ii) The proportion of genes that have a high correlation with their corresponding gene programs as a function of gene program number. For raw data (without denoising), a high correlation is defined as exceeding 4 standard deviations of the gene-gene correlation distribution. In the denoised data by MAGIC [1], a high correlation is defined as larger than 0.5. The plot shows that 30 programs is an elbow point of information gain, as detailed in Sec. 1.11. (C) (left) Uncertainty of inferred network interactions as a function of interaction strength quantified by the standard deviation among alternative network solutions, as detailed in Sec. 1.7. The uncertainty is generally below 0.02 and much lower compared with interaction strengths. (right) Histogram showing the proportion of edge agreement between alternative solutions of the D-SPIN network inference problem. Alternative solutions are obtained by sampling from the Bayesian posterior distribution given the experimental data, as detailed in Sec. 1.7. Alternative solutions of D-SPIN are highly consistent with the inferred network, with typically more than 98% of the edges being in the same category of activation, inhibition, and non-interacting. (D) Heatmap visualizations of the inferred program regulatory networks. Gene programs are grouped into modules, whose boundaries are marked by black lines. (E) (left-top) UMAP embedding of cell-cycle-related gene programs in control samples exhibits a circular structure. The clusters of cell cycle states are annotated and ordered based on the gene program expression and sequential positions on the UMAP. (left-bottom) Network diagram of the D-SPIN cell-cycle program subnetwork taken from the full 30-program network. (right-top) Heatmaps of cell-cycle programs for cell states from the Perturb-seq control samples and (right-bottom) the D-SPIN model under the control condition. D-SPIN reconstructs the distribution of cell states associated with cell cycle progression from the 6-node subnetwork. (F) The distribution of cell cycle states in generated data from the D-SPIN model and experimental Perturb-seq control samples. The distribution generated by D-SPIN quantitatively agrees with the data distribution with an 11% mean error. (G) Histogram showing the number of responding gene programs for each gene knockdown perturbation on a linear scale and (inset) log scale. The responding program number is exponentially distributed, suggesting that perturbations influencing a large number of programs are relatively rare. (H) Each scatter dot represents the inferred interaction strength of KLF1 to the erythroid program and SPI1 to the myeloid program for a specific gene knockdown condition. Dot colors indicate the number of cells in the knockdown condition. Most gene knockdown conditions assign positive weights to the two interactions. Highlighted conditions that generate high inferred weights are primarily knockdowns of epigenetic regulators.

**Figure S4:**
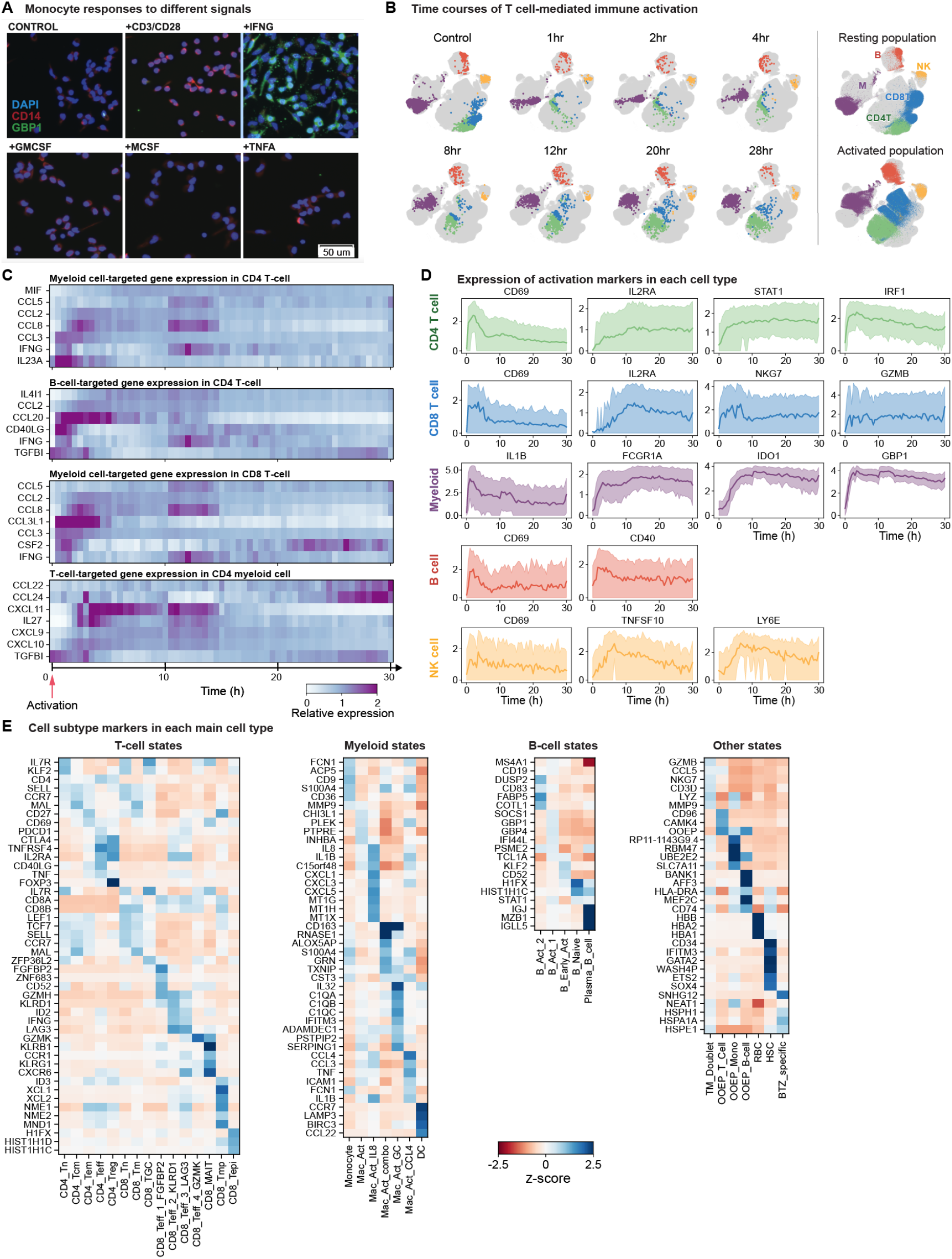
Timecourse of T-cell-mediated immune activation; cell typing of the drug profiling experiments; related to Figure 5. (A) Immunostaining of primary human monocytes cultured with CD3/CD28 antibody (25uL/mL cells), IFNG (5ng/mL), GM-CSF (5ng/mL), M-CSF (5ng/mL), or TNFA (5ng/mL). The monocyte activation marker GBP1 is only expressed under IFNG activation. The results show that the anti-CD3/CD28 antibody does not activate monocytes alone, and the observed monocyte activation in the PBMC is due to signaling between cell types. (B) UMAP embeddings of example time points in the 30-hour time-course experiment of T-cell-mediated immune activation, where samples were taken every 30 minutes for single-cell mRNA-seq. The immune population gradually moves from the resting state to the activated state. (C) Gene expression over time for selected signaling genes in four cell types. Expression is normalized by the mean expression across the time course. Many signaling genes are immediately upregulated in T cells after antibody activation, while in myeloid cells, signaling genes such as IL27 and CXCL11 are upregulated after a few hours, showing rich dynamics of communications between different cell types. (D) The dynamics of gene markers for immune activation in each major cell type. The color-shaded range is the 10th and 90th percentiles of gene expression. After T cell activation by the anti-CD3/CD28 antibody, major cell types in PBMC are activated with different time dynamics. (E) The heatmap shows the z-score of gene expression for cell type markers. Marker genes are identified by differential expression analysis of each Leiden cluster. The z-score is computed from the mean marker gene expression in each Leiden cluster compared to the mean and standard deviation in all the cells of the same major cell type (T cell, myeloid, B cell, and others).

**Figure S5:**
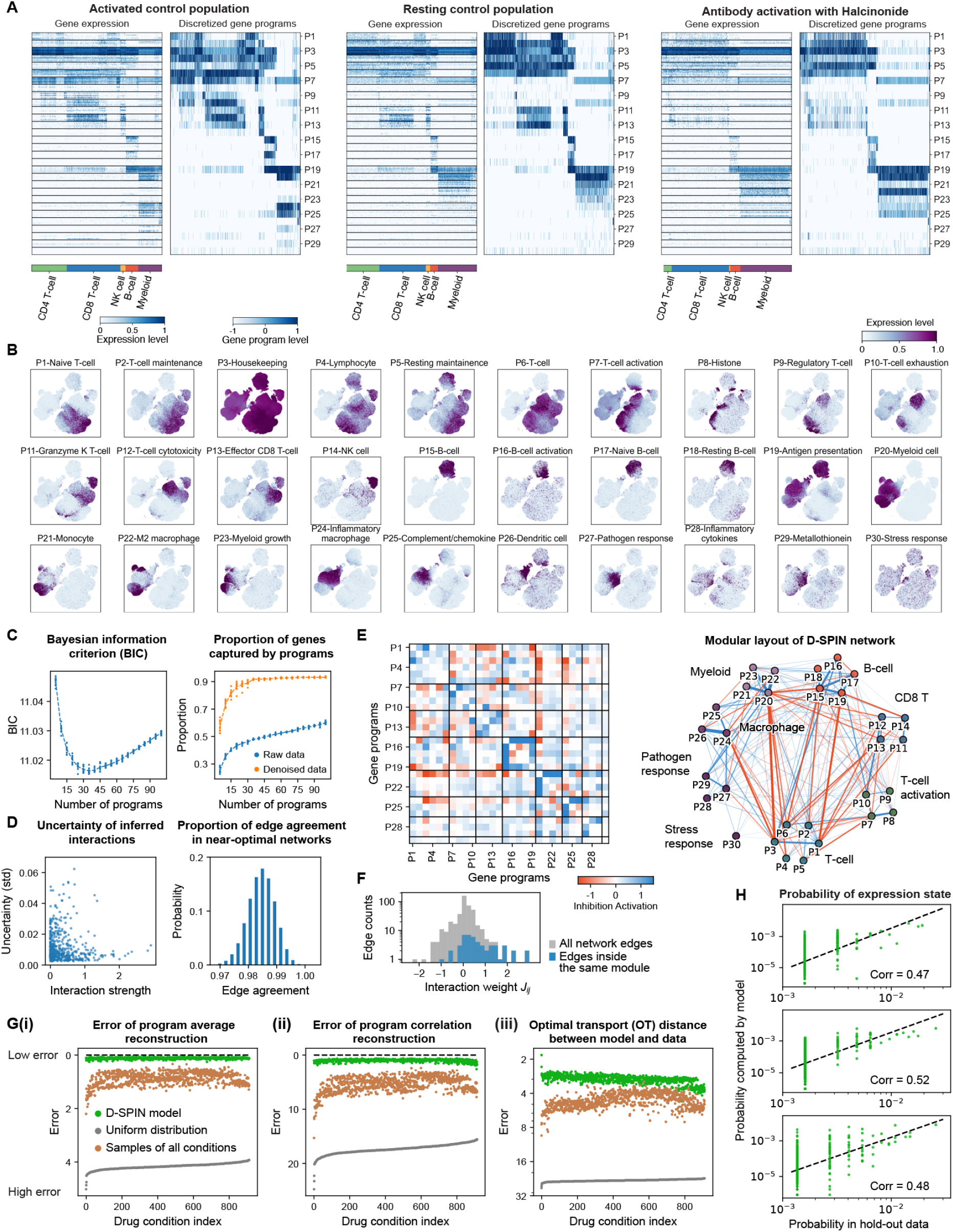
D-SPIN decomposes gene expression into programs and accurately reconstructs cell state distributions from a subset of training data; related to Figure 5. (A) Heatmaps of gene expression and discretized gene program level on (left) activated control population, (middle) resting control population, and (right) T cell receptor activation with halcinonide treatment. Discretized gene programs characterize and denoise major expression patterns in the gene matrix. (B) UMAP rendering of the expression of each gene program. Each cell is colored by its expression level on the gene program. The expression level is normalized by the 95th percentile for visualization only. Gene programs are specifically expressed by certain cell states, which are localized in separate regions on the UMAP. (C) (left) Bayesian information criterion (BIC) of the gene program decomposition as a function of gene program number. 30 is the optimal program number that minimizes BIC when the effective sample size is 15k cells. (right) The proportion of genes that have a high correlation with their corresponding gene programs as a function of gene program number. In raw data, a high correlation is defined as exceeding 3 standard deviations of the gene-gene correlation distribution. In the denoised data by MAGIC [1], a high correlation is defined as larger than 0.5. The plot shows that 30 programs is an elbow point of information gain, as detailed in Sec. 1.11. (D) (left) Uncertainty of inferred network interactions as a function of interaction strength quantified by the standard deviation among alternative network solutions, as detailed in Sec. 1.7. The uncertainty is generally below 0.06 and much lower compared with interaction strengths. (right) Histogram showing the proportion of edge agreement between alternative solutions of the D-SPIN network inference problem. Alternative solutions are obtained by sampling from the Bayesian posterior distribution given the experimental data, as detailed in Sec. 1.7. Alternative solutions of D-SPIN are highly consistent with the inferred network, with typically more than 97% of the edges being in the same category of activation, inhibition, and non-interacting. (E) (left) Heatmap visualization and (right) network diagram of the regulatory network model inferred by D-SPIN. The network exhibits a modular structure with tightly connected subnetworks. The 8 network modules correspond to major cell types or cell type functions in the PBMC population, such as T cells, B cells, and myeloid cells. Black lines in the heatmap mark boundaries of modules. (F) The histogram quantifies the distribution of all network edges (*J* matrix entries) and edges inside the same module. Edges inside modules are mostly positive interactions and contain the majority of strong positive interactions in the network. (G) Scatter plots of evaluating the distance between the 50% held-out experimental test data and the distribution generated by the D-SPIN model trained with the other 50% training data. Distribution distances are quantified by (i) Euclidean distance between the average of gene program expression, (ii) Frobenius norm of the difference between the cross-correlation of gene program expression, and (iii) optimal transport (OT) distance of probability distributions. The two null distributions are (gray) uniform distribution over all states and (brown) the distribution of pooling samples from all experimental conditions to reflect relative cell type abundance, as detailed in Sec. 2.14. Across all experimental conditions, the distributions defined by D-SPIN quantitatively agree better with the test data distribution compared with the two reference distributions. (G) Scatter plot comparisons between state distributions of held-out experimental data and the D-SPIN model fitted on training data. Even though the experimental observations are sparse samples of a few thousand cells in the total 3^30^ ≈ 2 × 10^14^ possible states, the probabilities of the empirical distribution and D-SPIN model probabilities are moderately correlated with correlation 0.47, 0.52, and 0.48. The plot shows that high-probability states in the D-SPIN model are also high-probability states in experimental data.

**Figure S6:**
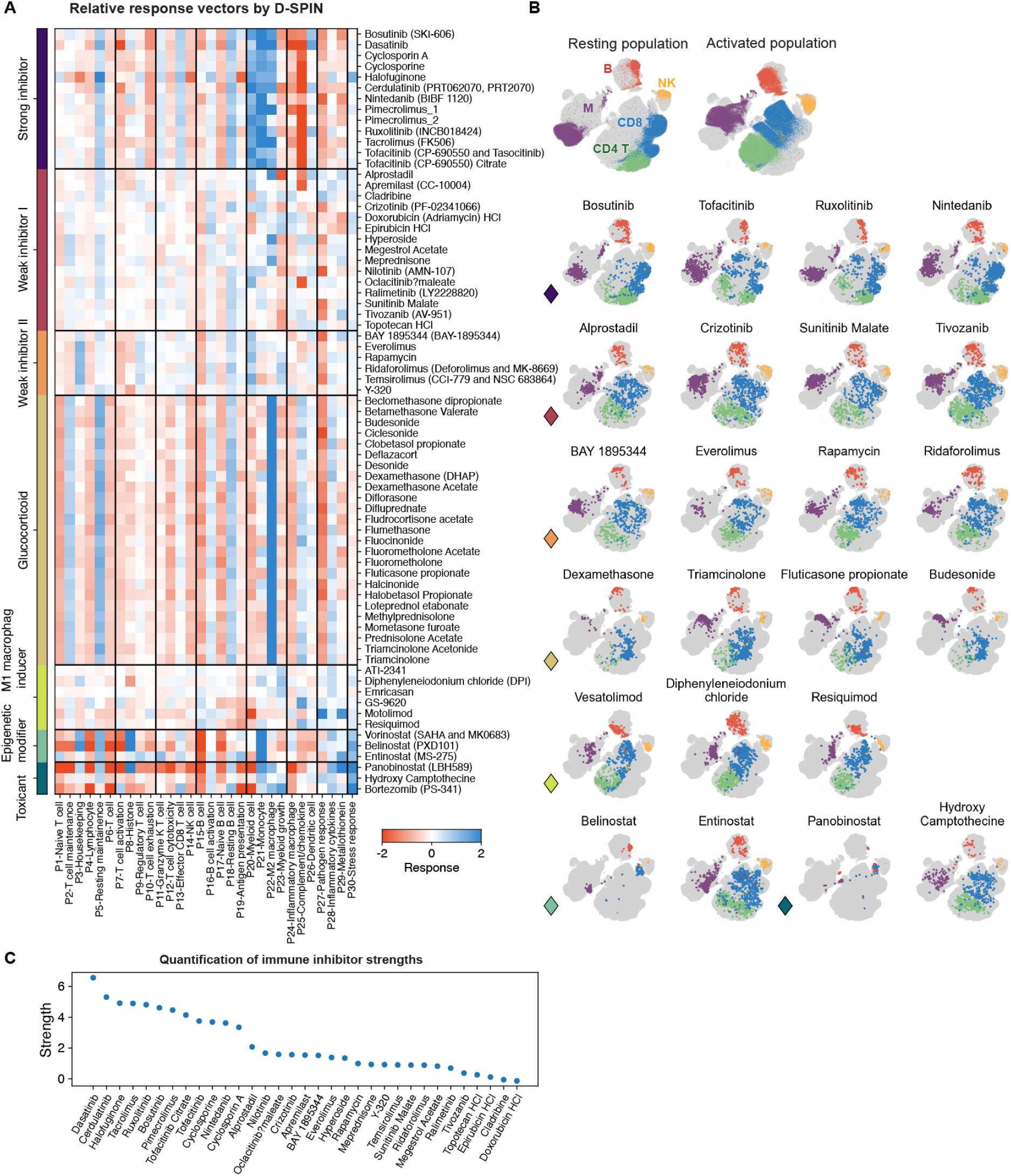
D-SPIN identifies phenotypical classes of drug categories; related to Figure 6. (A) Heatmap of drug relative response vectors inferred by D-SPIN. The relative response vectors are defined as the change of the drug response vector over control samples. D-SPIN identifies 70 effective drugs on the immune population that have statistically significantly different relative response vectors from control samples, as detailed in Sec. 2.12. We group these drugs into 7 phenotypical classes: strong inhibitor, weak inhibitor I, weak inhibitor II, glucocorticoid, M1 macrophage inducer, epigenetic modifier, and toxicant. (B) UMAP embeddings of example drugs from each class. Drugs in the same phenotypical class identified by D-SPIN exhibit similar cell population shifts on the UMAP embedding, while the cell population states vary greatly between different classes. (C) Strengths of immune inhibition of all D-SPIN-identified immune inhibitors. Inhibitor strengths are computed by projecting the relative response vector towards the leading singular vector of all inhibitors, as detailed in Sec. 2.15.

**Figure S7:**
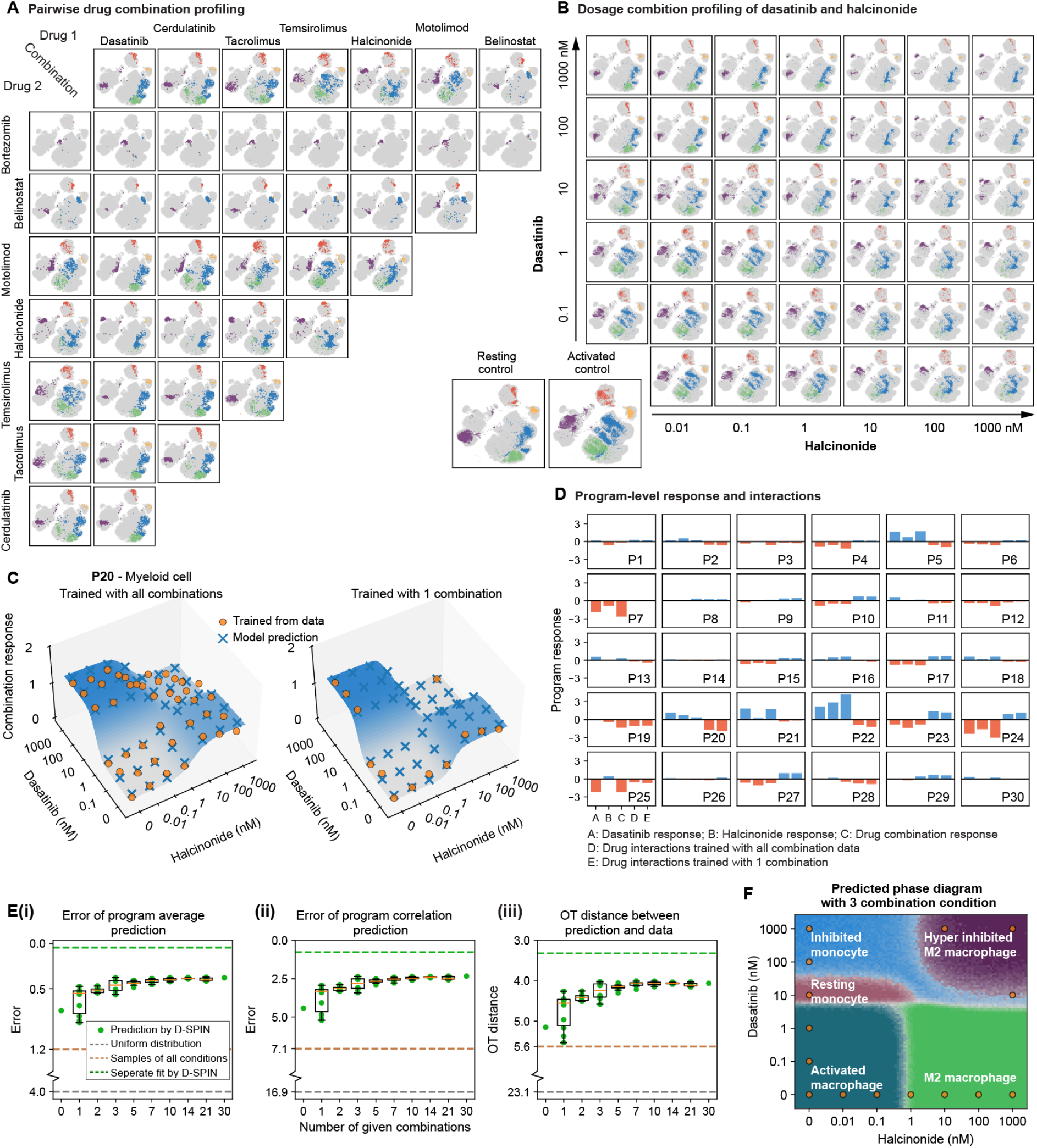
D-SPIN predicts continuous cell state transitions modulated by dosage combinations from a subset of combinatorial conditions; related to Figure 7. (A) UMAP embeddings of selected single drugs and pairwise drug combinations. We performed pair-wise drug combination profiling for 12 selected drugs (8 shown here) from different categories. (a) Strong inhibitor: broad-spectrum kinase inhibitor dasatinib, BCR-ABL, and Src tyrosine kinase inhibitor bosutinib, SYK/JAK kinase inhibitor cerdulatinib, calcineurin inhibitor tacrolimus; (b) Weak inhibitor: mTOR kinase inhibitor temsirolimus; (c) Glucocorticoid: halcinonide; (d) M1-macrophage inducer: motolimod; (e) Epigenetic modifier: vorinostat, belinostat; (f) Toxicant: bortezomib; (g) Others: forsythin, naproxen sodium. Forsythin and naproxen cause the loss of macrophage population in single-drug profiling, which is not observed in repeats in drug combination profiling. (B) UMAP embeddings of dosage combination profiling of dasatinib plus halcinonide. The two drugs create a monocyte-like resting myeloid population at a medium level of combination (10nM dasatinib, 1nM halcinonide) and a novel combinatorial myeloid state expressing both monocyte and M2 macrophage gene programs. On T cells, the drug combination also produces inhibited T cell states. (C) Surface plot shows response vectors on program P20 under different dosage combinations. The D-SPIN-inferred combinatorial response on P20 shows significant non-additivity, but is still well characterized by the sum of sigmoid models with a multiplicative interaction term. The model form allows prediction of dosage combination responses with only single-drug data and one drug combination condition. (D) Bar plots show the D-SPIN-inferred perturbation responses of each program for single drugs, drug combinations, and drug interactions. Drug interactions are defined as the difference between the combination effect and the sum of the single-drug effects. Observing one combination condition is adequate for D-SPIN to quantify interactions between drugs in each program. (E) Cell state distribution prediction error of D-SPIN with different numbers of drug combination conditions used for training. The error is quantified by the average (i) Euclidean distance between the average of gene program expression, (ii) Frobenius norm of the difference between the cross-correlation of gene program expression, and (iii) optimal transport (OT) distance of probability distributions. The two null distributions are (gray) the uniform distribution over all states and (brown) the distribution of pooling samples from all experimental conditions to reflect relative cell type abundance. The green line is the error of the D-SPIN model trained with all dosage combination conditions without the constraints of the responses following the sigmoid model with multiplicative interaction. (F) Predicted myeloid state phase diagram by D-SPIN with single drug responses and 3 drug combination samples. The cell state transition boundary closely matches the phase diagram learned with all dosage combination data.

## Key resources table

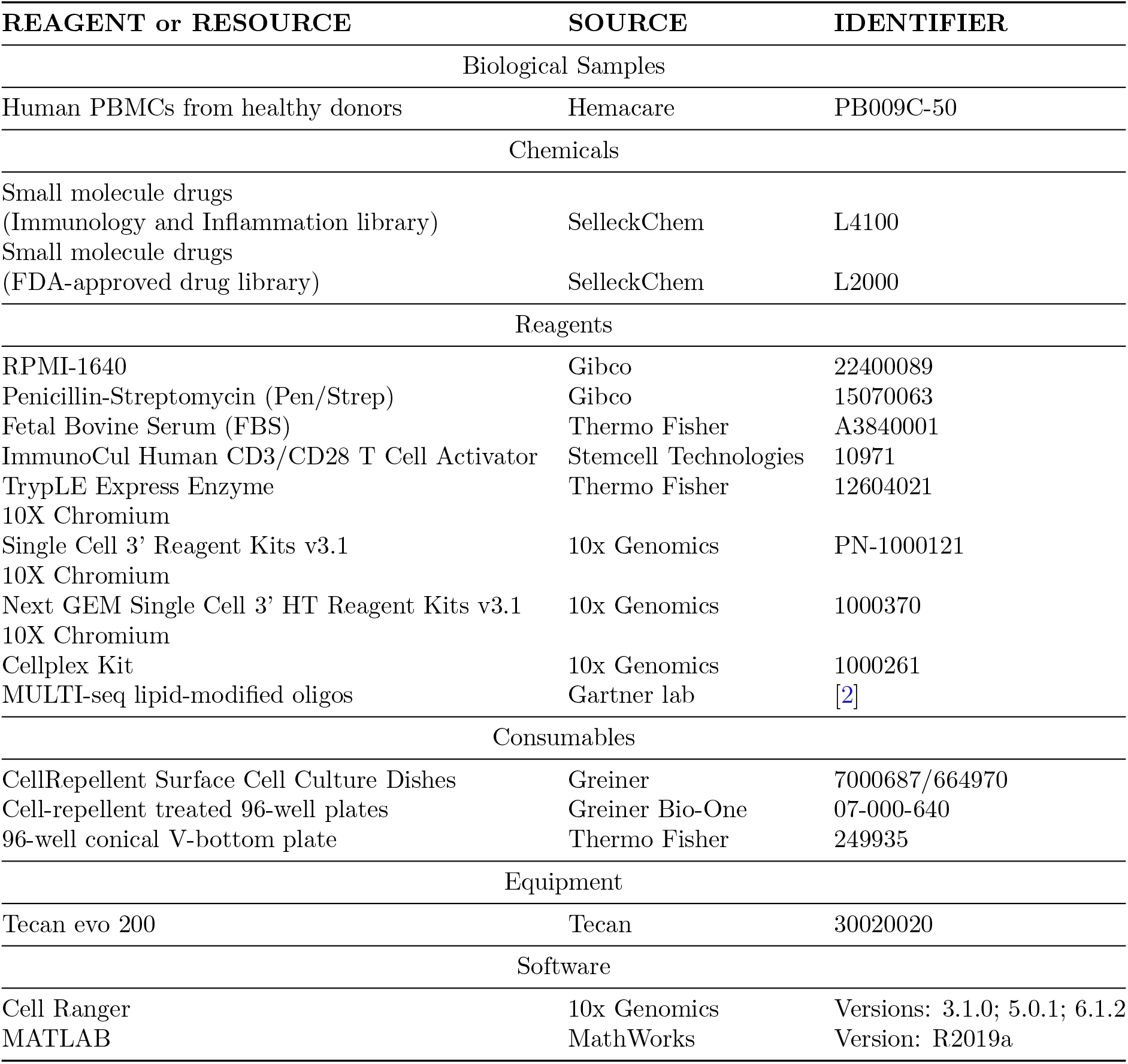

## Resource availability

### Lead contact

Further information and requests for resources and reagents should be directed to and will be fulfilled by the lead contact, Matt Thomson.

### Experimental model and subject details

We thawed cryopreserved human peripheral blood mononuclear cells (PBMCs) from healthy donors (Hemacare, PB009C-50, Lot 20062904) in a 37 °C water bath and transferred these to a conical vial. Pre-warmed RPMI-1640 (Gibco 22400089) was then added at a 10:1 ratio relative to the original cell suspension to dilute the cryoprotectant DMSO. Cells were pelleted for 2 min at 300 rcf and then resuspended in pre-warmed RPMI-1640 + 1% Pen/strep (Gibco 15070063) + 10% Fetal Bovine Serum (Thermo Fisher A3840001) to a final cell density of 10^6^ cells/mL. Cells were plated into CELLSTAR CellRepellent Surface Cell Culture Dishes (Greiner 7000687/664970) and then rested in a tissue culture incubator at 37 °C, 5% CO2 for 16 hours before being exposed to small molecule drugs.

## 1 Method details

### 1.1 Overview of D-SPIN framework

D-SPIN is a probabilistic network inference framework designed to analyze, interpret, and predict single-cell transcriptional responses measured across perturbation conditions. Given a single-cell perturbation dataset, D-SPIN learns (1) a unified regulatory network between modeled variables, which can be either single genes or gene programs, and (2) how each perturbation acts on the network that induces the observed changes of cell state distribution. A typical workflow of D-SPIN contains

1. Automated gene program discovery to identify the co-expressed transcriptional programs in the dataset.
2. Program-level network model construction that reveals the coordination between pathways and physiological functions in response to perturbation.
3. Gene-level network model construction that identifies signaling hubs and finer-grained perturbation response signatures.
4. Associating gene-level and program-level networks by identifying key regulators of program responses.

The program discovery and program-level models are optional when modeling a specific subset of target genes rather than transcriptional-scale cell states. Here, we first present a general overview and will delve into specific theories and implementations in subsequent sections.

The core theoretical development of D-SPIN is an accurate, versatile, and highly scalable statistical framework that infers a unified network model that simultaneously explains all observed perturbation responses. Conceptually, the diverse responses to perturbation input are all orchestrated by the same underlying regulatory circuitry; different perturbations selectively excite and repress distinct parts of the network, causing the cell population to explore the landscape of transcriptional states. By modeling all perturbation conditions together, we have achieved significantly improved network inference accuracy in both synthetic and biological dataset benchmarks.

D-SPIN applies a discretized spin-network modeling framework originating from statistical physics. The discretization can account for multiple different gene expression states while being resistant to counting noise in single-cell data. Further, discretization naturally generates multiple stable states and accommodates multiple cell types within a population and with a minimal interaction model, which can be interpreted as the steady state of a reaction system with nonlinear activations (Sec. 1.8).

We derived two variants of the inference algorithm under the same statistical framework that are suited for different scenarios (Sec. 1.3). One algorithm using pseudolikelihood is able to learn directed interactions between variables and is highly scalable for thousands of genes and millions of cells. The directed network inference is designed for building a gene-level understanding of the information flow between signaling pathways and identifying core regulators of biological processes. The other algorithm, using maximum likelihood, only learns undirected networks, but is generative and defines a complete probabilistic distribution of transcriptional states across multiple cell types. The generative model allows us to perform reasoning on the network model and generate predictions beyond the training data. The undirected network inference is also scalable to millions of cells, but is limited to a moderate network size of under 50 nodes due to the curse of dimensionality in defining full distributions in high dimensions. We build undirected network models on gene programs to capture the coordination between physiological functions to maintain homeostasis under perturbations.

For building program-level network models with more interpretability, D-SPIN contains an unsupervised pipeline of gene program discovery. Gene programs are defined as non-overlapping subsets of genes, as shared genes between programs will cause confounding interactions in the network. D-SPIN identified gene programs from the gene expression matrix by orthogonal non-negative matrix factorization (oNMF) to partition genes into gene programs (Sec. 1.9), but also can accommodate gene programs obtained by separate analysis or prior biological knowledge. As each gene is constrained to be assigned to one program, regulatory genes that control multiple biological processes are more likely to be misclassified and not grouped with genes that they control. To address the limit, we typically build single-gene networks on single-cell datasets to complement the program-level networks, and associate the two network models by identifying key regulators for each program from the single-gene network independent of the program components (Sec. 1.4). Therefore, D-SPIN can model shared regulatory control across pathways while preserving a non-overlapping program-level decomposition.

In addition to the regulatory networks, the inferred perturbation response vectors provide compact representations of the perturbation action on the network, enabling the clustering and visualization of perturbation responses across experimental conditions. Furthermore, the response vectors are minimal representations that can generate the entire cell state distribution together with the undirected network model. Therefore, we can generate predictions of cell state distributions by performing simple algebraic operations on the response vectors, such as predicting additive drug combinatorial responses, quantifying interactions between drugs, and extrapolating to unobserved drug dosage combination conditions (Sec. 2.17).

### 1.2 Formulation of D-SPIN framework

The probabilistic graphical model we used in the formulation of D-SPIN originates from the Ising model in statistical physics, also called a spin glass, spin network, or Markov random field in different disciplines. The model defines a probability distribution on a discretized state vector ***s*** = [*s*_1_, *s*_2_, …, *s*_*M*_], *s*_*i*_ ∈ {−1, 0, 1}, where *M* is the number of genes or gene programs. The probability distribution defined by the model is

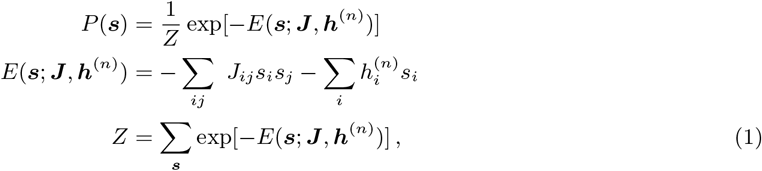

where *E* is the energy function and *Z* is a normalizing constant called the partition function. States with lower energies have higher probabilities; therefore, positive interactions *J*_*ij*_ *>* 0 promote *s*_*i*_ and *s*_*j*_ being simultaneously on or off, and negative interactions promote the two variables being at different states. The bias vectors *h*_*i*_ increase probability of the variable *s*_*i*_ being on (*h*_*i*_ *>* 0) or off (*h*_*i*_ *<* 0).

D-SPIN assumes that across perturbation conditions, the interaction network ***J*** stays the same and the bias vector ***h***^(*n*)^ is modulated by each perturbation condition *n* to activate different modes in the network and to produce different cellular transcription state distributions. We train the model from data by gradient ascent of the log-likelihood function.

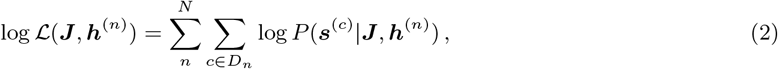

where *c* is the cell index and *D*_*n*_ is the set of all cells in the *n*-th experimental condition. We then compute the gradient of the objective function as

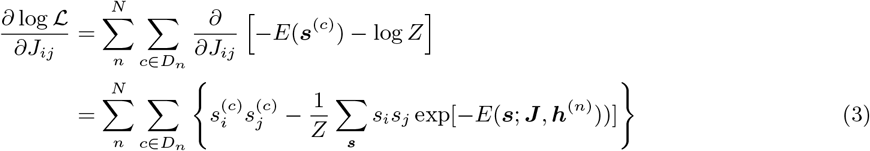

The first term is the sum *s*_*i*_*s*_*j*_ over all experimental samples, and the second term is a constant, which is the expectation of *s*_*i*_*s*_*j*_ of the current model, defined by the parameters ***J*** , ***h***. We normalize the gradient by sample number to improve numerical stability under a given step size. With a similar derivation for *∂* log ℒ*/∂h*_*i*_, the gradients of the objective function have the following form.

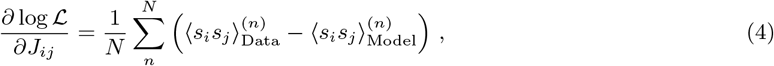

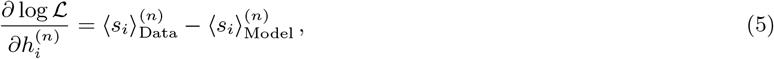

where 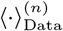 is the average over the data of *n*-th conditions, and 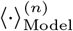 is the expectation on the distribution defined by current model parameters ***J*** , ***h***^(*n*)^. Therefore, the modeling learning is essentially matching the cross-correlation ⟨*s*_*i*_*s*_*j*_ ⟩ and mean ⟨*s*_*i*_⟩ between the model and data.

The model has a few unique advantages:

1. From a statistical perspective, the spin network is the maximum entropy model given the mean and pairwise cross-correlation of data [3]. Entropy describes how a distribution is “spread out” over all possible states, and the principle of maximum entropy states that the best model describing a system is the model that generates data distributions with the maximum entropy while agreeing with relevant statistics of the data. Such a model has no extra assumptions about the structure of the system apart from the measured statistics.
2. From a computational perspective, the inference of the spin network, i.e., maximization of the log-likelihood function, is a concave problem where the only local maximum is the global maximum (Sec. 1.6). Therefore, optimization techniques to avoid the traps of local optima are not necessary. However, it is worth noting that the optimization is still a difficult NP-hard problem in the field of computational complexity [4], primarily due to the computational complexity of accurate gradient estimation and large condition numbers from model identifiability issues [5]. Inferring the model across perturbation conditions can mitigate these issues, and we computationally evaluated the curvature of the inference problem in our dataset (Sec. 1.7) and explored it further theoretically in another work [6].
3. From a physics perspective, the spin network is a Boltzmann distribution defined by the energy of each state. The model can be connected to the distribution of microscopic states in thermodynamics and dynamics of a microscopic system under thermal noise (Sec. 1.8)

There are a few potential choices of discretizing the expression level ***l***: the number of discretized states *m* and the choice between {−⌊*m/*2⌋, −⌊*m/*2⌋+ 1, … , ⌊*m/*2⌋− 1, ⌊*m/*2⌋} or {0, 1, … , *m*}. We choose to discretize ***l*** into 3 states {−1, 0, 1} due to the following reasons:

1. Larger *m* increases computational complexity, and makes the model closer to a Gaussian distribution that has a single probability density maximum.
2. We find *m* = 2 is insufficient to characterize phenomena in our drug profiling experiments. We observed different levels of program activation instead of on-and-off switching. For example, both glucocorticoid drugs and immune inhibitors activate the program P22 M2 macrophage, but with different expression levels.
3. The choice {−1, 0, 1} is preferable because the self-interactions term *J*_*ii*_ has more clear biological interpretations. Considering a single program only, self-activation *J*_*ii*_ *>* 0 is similar to a bi-stable switch produced by nonlinear activation, where −1 and 1 states are preferable. Similarly, self-inhibition *J*_*ii*_ *<* 0 is similar to negative feedback, where the 0 state is preferable.
4. In the setting of gene perturbation experiments, the three states -1, 0, and 1 can also be interpreted as inhibited, unperturbed, and activated to match with experimental settings.
5. On the network inference benchmarking on synthetic networks, there is no significant difference between the choice of {−1, 0, 1} and {0, 1, 2}.

### 1.3 Computational methods for inference in D-SPIN

Various computational methods have been proposed to solve the spin network inference problem [3], and we adapt three of the most accurate methods to the context of D-SPIN. Each method has its advantages and specific niches depending on the problem setting, such as network size, number of cells, and number of perturbation conditions.

#### Exact maximum likelihood inference

For small networks, the probability distribution *P* (***s***|***J*** , ***h***) can be explicitly computed, thus the exact gradient of ***J*** , ***h*** can be computed using Eq. 4, Eq. 5. With the exact gradient, the optimization problem can be solved using gradient ascent or other optimizers such as Momentum or Adam.

As the gradient estimation requires enumerating all possible states, the computational complexity scales exponentially with the number of nodes. Though exact, the method is only applicable to small networks of around 10 nodes (3^10^ ≈ 6 × 10^4^ states).

#### Markov Chain Monte Carlo (MCMC) maximum likelihood inference

As the gradient only requires computing the mean and cross-correlation of the samples, we can approximate the complete distribution *P* (***s***|***J*** , ***h***) by sampling an empirical distribution. Without evaluating the exact distribution, we can construct a Markov Chain between states whose stationary distribution is *P* (***s***|***J*** , ***h***). Specifically, we utilize the Gibbs sampling scheme. Starting from a random initial state ***s*** = [*s*_1_, *s*_2_, … , *s*_*M*_], at each step we randomly take an index *k* in the *M* nodes, and update the value of *s*_*k*_ by its conditional distribution given all other nodes *P* (*s*_*k*_|*s*_1_, … , *s*_*k*−1_, *s*_*k*+1_, … , *s*_*M*_ , ***J*** , ***h***). After a burn-in period of steps to allow the Markov Chain to equilibrate, the sequence of samples is an empirical distribution of *P* (***s***|***J*** , ***h***) and can be used to estimate the gradient Eq. 4, Eq. 5.

In each sampling step, computing the marginal distribution is of complexity 𝒪(*M*). For accurate cross-correlation estimation, the required number of samples scales as 𝒪(*M* ^2^), without considering the burn-in period of the MCMC process. Therefore, the overall computational complexity is at least 𝒪(*M* ^3^). The MCMC method applies to medium-sized networks up to 30 ∼ 50 nodes.

#### Pseudolikelihood method

The major challenge involved in scaling inference to large networks is the partition function *Z* in the distribution *P* (***s***|***J*** , ***h***), which involves exponentially many terms with the network size *M* . An alternative approximation method called pseudolikelihood was developed originally for spatial statistics and adapted to spin network problems [3, 7, 8]. Pseudolikelihood methods approximate the distribution *P* (***s***|***J*** , ***h***) as the product of the conditional distribution of each individual variable, and provide exact inference in the limit of an infinite number of samples.

Specifically, we denote ***s***_\*k*_ = [*s*_1_, … , *s*_*k*−1_, *s*_*k*+1_, … , *s*_*M*_]^⊺^ as the state ***s*** except *s*_*k*_. The pseudolikelihood function for the inference problem of a single condition is

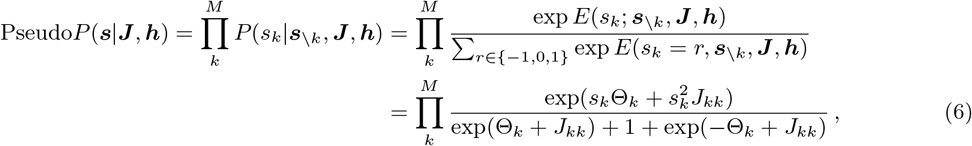

where Θ_*k*_ = *h*_*k*_ + ∑_*j*≠*k*_ *J*_*jk*_*s*_*j*_ is the effective field conditioned on all other nodes ***s***_*\k*_. The pseudolikelihood function decouples the mutual dependence between nodes, thus removing the exponentially complex partition function *Z*. As a cost, the pseudolikelihood function in general does not sum to 1, and thus is not a distribution; this is where the “pseudo” name comes from. The gradient for the log pseudolikelihood objective function is

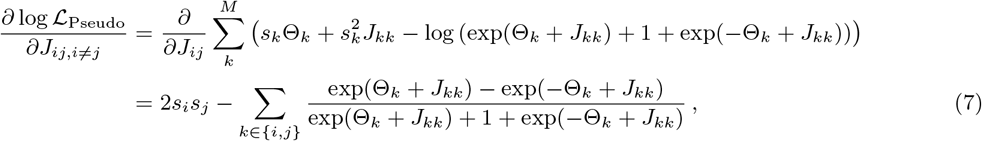

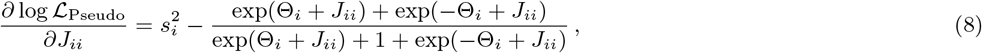

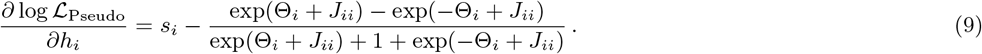

The computational complexity of the gradient computation scales with 𝒪(*M* ^2^). Also, note that the effective field Θ_*k*_ depends on each cell state ***s***, so the gradient computational scales with the total number of cells. Practically, the pseudolikelihood method is highly scalable and applies to networks of thousands of nodes. The approximation of pseudolikelihood is more accurate when the number of samples is high. However, typically the number of observed samples is far lower than the total number of possible states (3^*M*^). Therefore, exact maximum likelihood and MCMC maximum likelihood are preferred when they are computationally feasible.

#### Inferring direction of interactions

Furthermore, the form of pseudolikelihood is closely related to regression models, enabling the assignment of directionality to the inferred network. The distribution of a single node conditioned on all other nodes

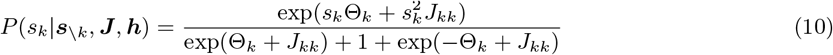

can be interpreted as a regression problem, where we predict the state of the dependent variable *s*_*k*_ with other variables are predictors. The interactions *J*_*ij*_ are the coefficients of the regression problem. If gene A predicts gene B better than B predicts A, this suggests a regulation direction of A to B. To compute the directional network, the gradient estimation Eq. 7 can be simply replaced with an asymmetric version of

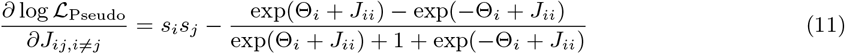

Even though D-SPIN is capable of inferring a directed network, we focus our analysis on undirected networks because directed networks cannot define a stationary distribution on the data when time information is not supplied. In the context of probabilistic graphical models, directed models are always constrained to be acyclic, i.e., with no cycle. Such a constraint is reasonable in the field of causal inference, where the circularity of causal relations is rare. However, in cellular regulatory networks, feedback loops are prevalent to maintain homeostasis or signal amplification. For example, the regulatory network models of hematopoietic stem cell (HSC) differentiation contain several loops, such as PU.1-GATA1-GATA2-PU.1 [9].

The acyclic constraint of directed probabilistic graphical models is fundamental, as cycles in the conditional dependence between variables will produce inconsistent distributions. As mentioned in the seminal work of Judea Pearl on causal inference [10, 11], even three cyclic dependent variables “will normally lead to inconsistencies”, and the acyclic constraint can ensure consistency of the distributions. Constructing distributions on even two mutually dependent variables requires nontrivial constraints [12]. Another approach to include directionality in regulatory network models is to include explicit time dependence, which also has limited application due to the lack of experimental approaches to measure the dynamical evolution of transcriptome profiles. Therefore, in the context of probabilistic models of regulatory networks, undirected networks are the more appropriate choice.

### 1.4 Identifying regulators of gene programs from gene-level networks

The directed network inference method can also link gene-level networks with gene program expression to uncover how these programs are regulated. A joint model that contains both single genes as candidate regulators, as well as gene programs, has the form of

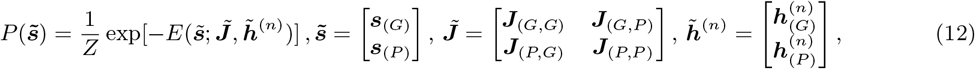

where ***s***_(*G*)_, ***s***_(*P*)_ are discretized single-gene states and program states to be concatenated together. 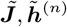 are concatenated matrix characterizing the regulatory interactions and perturbation responses of the concatenated state 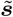. ***J***_(*G,P*)_ therefore characterizes the regulatory interactions from genes to programs. However, the straightforward joint model construction does not properly characterize the regulations between genes and programs. Gene programs are not separate physical entities and represent the collective activity of genes. The regulatory interactions between programs in the program-level model are a phenomenological, collective description of regulations carried out by “hidden” gene variables. Therefore, in the joint model, these collective descriptions of regulations should be removed in the presence of explicit modeling of gene-level interactions, in other words, ***J***_(*P,P*)_ = **0** and ***J***_(*P,G*)_ = **0**.

Another constraint on the equation is on the offset term 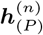 of the gene programs. Because the single-gene network model 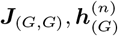 already contains the perturbation-dependent bias, and we seek to explain the expression change in the programs solely from the gene regulators, the programs do not require an additional perturbation-dependent offset. Conceptually, the condition-dependent program/gene offset is to carve out variations that do not come from the regulation of other programs/genes, which is inapplicable to programs in the joint model with both single genes and programs. In practice, including the perturbation-dependent offset to gene programs will deprioritize the weights of regulatory genes and deteriorate the results. With the discussed constraints, the joint model now has the form of

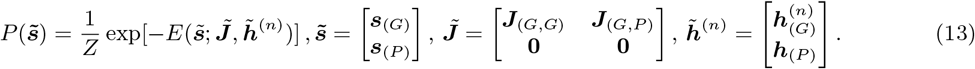

These additional constraints further simplify the mathematical optimization of the joint model. As the gene-level states ***s***_(*G*)_ have no dependence on the program-level states, inference of ***J***_(*G,G*)_ and 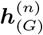 is now split from the joint model and identical to inferring these parameters alone in the gene-level network model. As the program states ***s***_(*P*)_ for each program only depends on gene states ***s***_(*G*)_ but not other programs, the inference of ***J***_(*G,P*)_ and ***h***_(*P*)_ can be performed separately for each program and the pseudolikelihood gradient becomes accurate likelihood gradient of the model. For a gene program *k*, denoting *w* = ***s***_(*P*) *k*_, ***j*** = ***J***_(*G,P*) ·,*k*_, *h* = ***h***_(*P*) *k*_, together with gene expression state {***s***_(*P*)_}, the likelihood function of the program state with gene regulation ***j*** and global bias *h* is

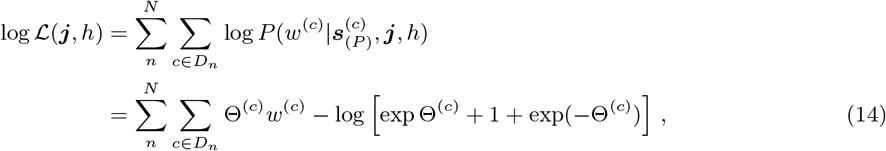

where *n* is the index for perturbation conditions, and 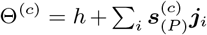 is the effective field on the program created by the state of single genes and the global bias *h* in each cell. The form is similar to Eq. 11, thus the same optimization procedures can be applied.

### 1.5 Regularization of the inference

In statistical inference, it is common to leverage prior knowledge about the potential form of the solution, a process known as regularization. Regularization nudges the solution towards a preferred direction, which can also be interpreted as assigning a prior distribution of the model parameters in the framework of Bayesian inference. For example, *ℓ*_1_ (Lasso) regularization promotes sparsity of the solution, while *ℓ*_2_ (Ridge) regularization promotes solutions with smaller magnitudes.

Specifically, in the D-SPIN framework, we sometimes have prior knowledge of the perturbation action from the experimental design. For example, in single-cell profiling of gene knockdown or activation, the response vector relative to the control should have a strong inhibition/activation at the target gene. Suppose we have an estimation of the relative action of all the perturbations as 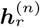. Then we can infer shared unperturbed single gene (program) activity ***h***_0_ and penalize the difference between ***h***^(*n*)^ − ***h***_0_ and 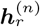 by *ℓ*_2_ norm. In this case, the objective function becomes

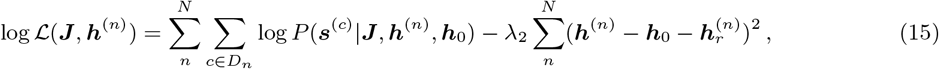

where *λ* is the strength of the regularization, representing the uncertainty of the prior knowledge of the relative response estimation. For this objective function, the gradient of ***J*** stays the same and

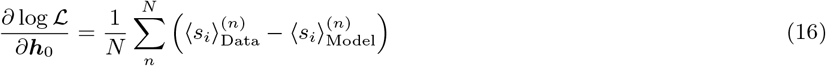

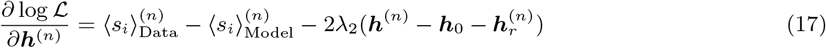

Similarly, the inference of the network can also be improved with prior knowledge of the network architecture. Methods like CellOracle or SCENIC+ can utilize information of TF-motif binding or chromatin accessibility from ATAC-seq measurements [13, 14]. Typically, these prior network contains a list of candidate interactions with unknown strengths, so regularization can be applied to edges that do not belong to the prior network. Suppose we denote the Boolean adjacency matrix of the prior network as ***J***_0_, an example *ℓ*_1_ regularization would be

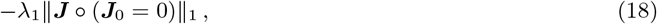

where the ∘ symbol is element-wise product ant ∥ · ∥_1_ is the *ℓ*_1_ norm.

### 1.6 Proof of convexity of the inference problem

We prove that the multi-condition/multi-***h*** inference problem by D-SPIN is a convex optimization problem. Specifically, the log-likelihood objective function is concave. Therefore, the only local optimum is the global optimum, and the solution to the optimization problem is unique.

For a single condition, the D-SPIN formulation is the same as the inverse Ising problem, whose log-likelihood function is concave, and strictly concave if all coupling and fields are finite. We follow the structure of the proof from a review [3]. We denote ***λ*** = {***J*** , ***h***}, and *Q*_*k*_(***s***) = {*s*_*i*_*s*_*j*_, *s*_*i*_} for the model parameters, and the log-likelihood can be written as

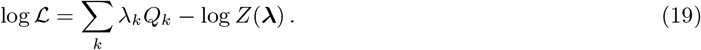

The second derivative of the log-likelihood function follows

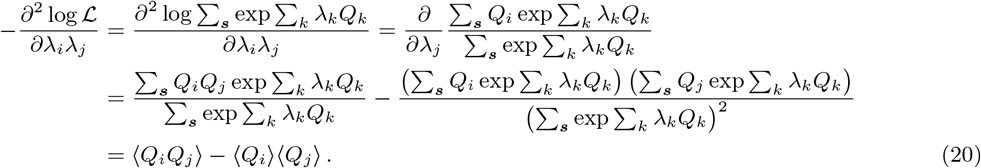

The negative matrix of second derivatives 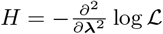 is positive semidefinite because for any vector ***x***,

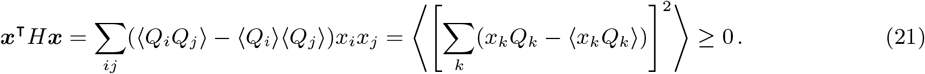

Further, the matrix is positive definite if no observable *Q*_*k*_ has vanishing fluctuations, which holds when all the couplings and fields are finite. Therefore, the log-likelihood function is strictly concave with finite couplings and fields, and the maximum likelihood estimation is a convex optimization problem.

For the formulation of D-SPIN of multiple conditions, we prove the case with two conditions, and the proof of more conditions naturally follows. We start with proving a lemma:

Lemma: Let *A*1, *A*2, *B*1, *B*2, *C*1, *C*2 be matrices such that the block matrices 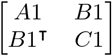 and 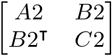 are both positive (semi)definite. Then the block matrix

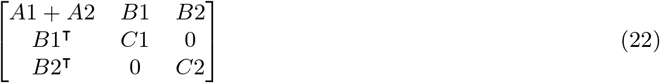

is also positive (semi)definite.

Proof: For any test vector ***x***, we denote 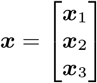, where each component is compatible with the shape of *A*_1_ + *A*_2_, *C*_1_ and *C*_2_, then we have

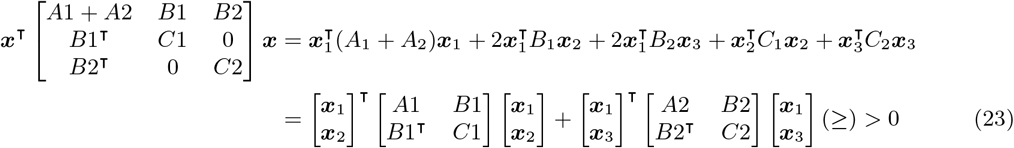

as both matrices are positive (semi)definite.

Similarly, we denote the parameter set ***λ*** = {***J*** , ***h***^(1)^, ***h***^(2)^}, and the log-likelihood function of the two conditions are log ℒ^(1)^, log ℒ^(2)^, and the second order derivative of the objective function follows

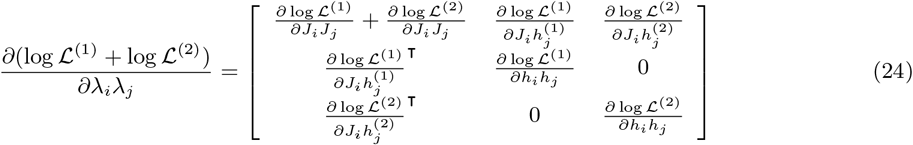

which is the rearrangement of the block matrices of the second-order derivative of the inverse Ising problem.

According to the lemma, the negative of the second-order derivative matrix is positive semidefinite, and positive definite if all couplings and fields are finite. This concludes our proof that the formulation of D-SPIN is a convex optimization problem.

### 1.7 Quantification of the inference uncertainty

Even though the optimization problem of D-SPIN inference has a unique solution, there is a possibility that there are other sub-optimal solutions that are distinct from the true solution but have similar values of the objective function. These alternative solutions are phenomenologically referred to as “flat valleys” of the optimization landscape, or “sloppy directions” of models. This flatness is quantified by Fisher information, which is a metric describing the curvature in the space of a parametric family of probability distributions. Large Fisher information indicates that small changes in the model parameter cause great changes in the distribution, while small Fisher information indicates that the distribution is insensitive to the parameter changes.

The Fisher information and uncertainty of the inference are connected by the Cramér-Rao bound, that for any unbiased estimator, the variance is lower bounded by the inverse of the Fisher information. Specifically, the bound is achieved by efficient estimators, and the maximum likelihood estimator is asymptotically efficient. Therefore, estimating the Fisher information matrix of the inference problem provides an estimation of the variance of the inferred network.

Specifically, using the same notation ***λ*** = {***J*** , ***h***}, and *Q*_*k*_(***s***) = {*s*_*i*_*s*_*j*_, *s*_*i*_} as in the previous section, the Fisher information matrix of a single condition can be computed by definition

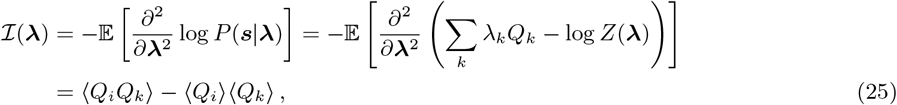

which has exactly the same form as Eq. 20 as the first linear term ∑ _*k*_ *λ*_*k*_*Q*_*k*_ vanishes under second order derivative. The bracket ⟨·⟩ is average over *P* (***s***|***J*** , ***h***), which is not directly available but can be estimated by the empirical distribution defined by samples. In practice, the Fisher information of a specific condition is estimated by

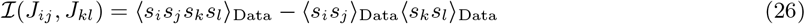

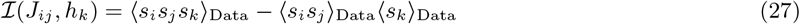

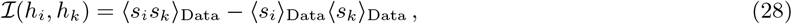

where the bracket ⟨·⟩_Data_ is average over all states in the data.

In the case of multiple samples, the overall Fisher information of the Fisher information has a similar form as Eq. 24. We also only write for two conditions for simplicity of notation.

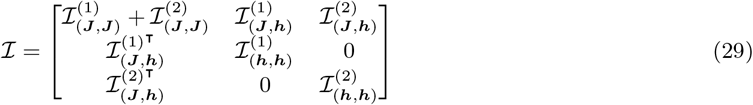

As we have proved, ℐ is positive definite when all couplings and fields are finite, ℐ_(***J***,***J***)_ and ℐ_(***h***,***h***)_ are all invertible. According to the Schur complement of the block matrix, we have

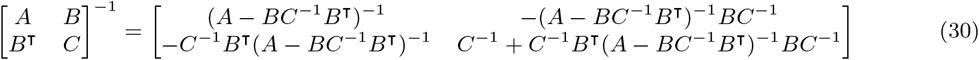

Therefore, the Cramé-Rao bound of the network estimation for *N* conditions is

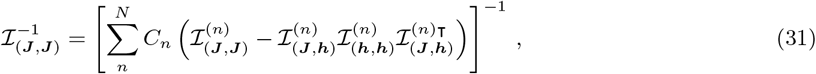

where *C*_*n*_ is the number of cells in the *n*-th condition. Specifically for each network edge *J*_*ij*_ , the diagonal term of the Cramé-Rao bound 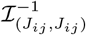 quantifies the variance of the inference uncertainty of the specific edge as presented in Figures S3C and S5D.

The inverse of Fisher information 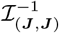 is the covariance matrix of the inferred network. Further, according to the Bernstein-von Mises theorem, the posterior distribution of the inferred network will converge to a normal distribution

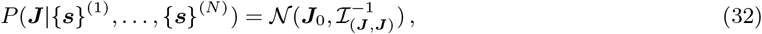

where ***J***_0_ is the result of inference. Therefore, we can sample from the posterior for alternative networks to validate the robustness of the inferred regulatory network as presented in Figures S3C and S5D.

### 1.8 Correspondence between D-SPIN and biochemical reaction systems

D-SPIN is closely connected to dynamical models of biochemical reactions as an approximation of the steady state of a chemical reaction system with saturating non-linear activation functions. For a biochemical reaction system with state *x* ∈ ℝ^*M*^ , the general dynamics can be written as

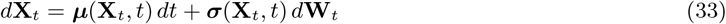

where ***µ***(**X**_*t*_, *t*) are the reactions and ***σ***(**X**_*t*_, *t*) *d***W**_*t*_ is the stochastic noise. If the stochastic noise can be assumed a constant *σ*, then the state distribution *P* (***x***) follows the Fokker-Planck equation

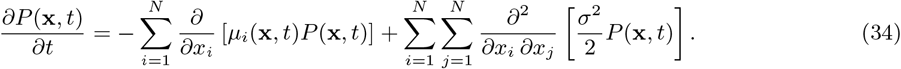

Specifically, if the reaction term can be written as the derivative of a potential function *µ*_*i*_(***x***) = −*∂V* (***x***)*/∂x*_*i*_, the equation has a Boltzmann distribution as the steady state solution for *∂P* (***x***, *t*)*/∂t* = 0

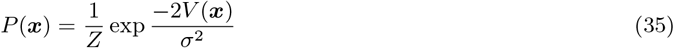

where *Z* is the normalization constant. A typical example is a linear expansion near the stable point of a dynamical system, where

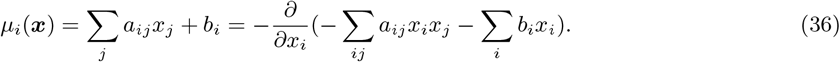

This solution is a Gaussian distribution near the stable point [15, 16]. Such models only have a single minimum in the convex potential function, and are therefore not suitable for characterizing cell populations with various stable cell types and cell states.

The potential function, i.e., energy function Eq. 1 in D-SPIN, is in a similar form to the Gaussian distribution as a second-order polynomial of variables. However, the energy in D-SPIN is a function of discretized state variables, which can be viewed as the discrete limit of composing the continuous state *x*_*i*_ with a saturating activation function *s*_*i*_ = *ϕ*(*x*_*i*_), for example, a sigmoid function for a two-state model. Given the potential function, the reaction term in the corresponding dynamical model is

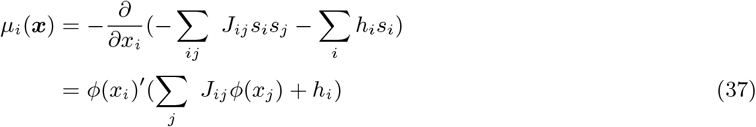

The first term *ϕ*(*x*_*i*_)^*′*^ is the derivative of the activation function, which can be approximated by a constant in the transition regime of the activation function, and close to 0 at the saturating regime of the activation function. The corresponding interpretation is that during the transition between states, the regulation received by component *i* is a linear function of the activation *ϕ*(*x*_*j*_) of all other components *j*. The nonlinearity is conceptually similar to the Hill function used in models of genetic circuits and allows the existence of multiple stable states controlled by the regulatory networks in the cell.

The structure of the dynamical model also helps to clarify the physical interpretations of the biasing vector ***h***. In classical causal discovery literature of directed acyclic graphs (DAGs), perturbations, specifically loss-of-function (LOF) experiments, are modeled by a “surgery” operation. Specifically, “surgery” means fixing the value of the perturbed variable and eliminating all incoming edges to ensure the variable is not influenced by other variables [17]. “Surgery” sets the target gene to a specific level (zero expression, for example) and blocks all other incoming regulations, while ***h*** is equivalent to modulating the baseline production rate or decay rate in the corresponding biochemical system.

In single-cell perturbation experiments, the rate-modulation formulation better represents the molecular implementation of perturbation compared to clamping the value of target gene expression. For instance, in Perturb-seq knockdown experiments by CRISPRi, the knockdown is primarily achieved by guide RNA recruiting dCas9 to block transcription, i.e., decreasing production rate; therefore, the knockdown efficiency is not 100% and the targeted gene still has low expression, which potentially can still be regulated by other genes. In the genome-wide K562 Perturb-seq experiment, many gene knockdowns have less than 70% knockdown efficiency [18]. Therefore, the choice of characterizing the perturbations by a bias term ***h*** is motivated by biological considerations, which turns out to be effective in modeling practice.

Furthermore, the correspondence between D-SPIN and the biochemical reaction system allows us to understand how the stationary distributions of directed regulatory interactions are mapped to undirected networks in D-SPIN. The generative distribution modeling of D-SPIN constructs an undirected regulatory network, while the biochemical reaction network naturally permits directed interaction between genes. The key information encoding directionality, the temporal information, is mostly lost in single-cell experiments with only snapshot observations of the steady state. For simplicity, we use the linear model to derive the relationship between the interaction matrix in the dynamical system and the quadratic terms in the stationary distribution, which are comparable to the interaction ***J*** in D-SPIN.

Specifically, for ***µ***(***x***) = *A****x*** + ***b*** in the matrix form, assuming *A* is invertible and structurally stable, the dynamical system

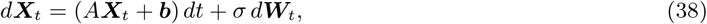

has a stationary distribution

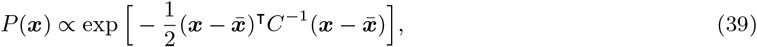

where the stationary mean 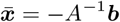 and covariance matrix *C* given by the Lyapunov equation

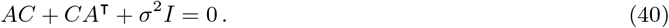

When the interaction matrix *A* is symmetric and negative definite, the equation has a unique solution *C* = −*σ*^2^*A*^−1^*/*2. The quadratic term in the distribution *P* (***x***) is the interaction matrix *A* up to a constant factor.

The covariance matrix *C* does not have an analytical solution when *A* is asymmetric, which corresponds to a directed chemical reaction network. For a minimal example, consider a two-gene network with directional regulation from *x*_1_ to *x*_2_

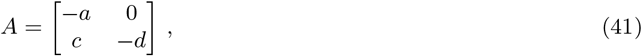

the approximate solution in the weak interaction limit (*c*^2^ ≃ 0) is

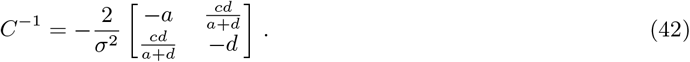

The form of the interaction term reveals how the directed interaction is reflected in the stationary distribution. When the regulator gene *x*_1_ has a fast degradation rate *a* relative to *x*_2_, the distribution of *x*_1_ and *x*_2_ becomes uncorrelated and exhibits weak interactions. Conversely, when the regulator has a slow degradation dynamics, *x*_1_ and *x*_2_ simultaneously turn on in the gene expression dynamics, and the inferred undirected network accurately reflects the interaction between *x*_1_ and *x*_2_. Therefore, the undirected network provide better approximation to the directed biochemical network when the regulator is more stable compared to the effector.

### 1.9 Gene program discovery by oNMF

We use orthogonal nonnegative matrix factorization (oNMF) for gene program discovery because oNMF generates a set of programs that are mathematically constrained to provide a high-accuracy representation of transcriptional states in the data, but with no overlap between gene programs. Compared to typical matrix factorization methods like principal components analysis (PCA), oNMF applies two constraints to the gene programs: non-negative weights and orthogonality. We used oNMF based on the following considerations:

1. Linearity: To ensure the interpretability of the model, each gene program is a linear combination of single genes
2. Non-negativity: The non-negative constraint avoids the ambiguity of interpreting negative weights, ensuring the programs are a set of co-expressed genes. Specifically, in the context of regulatory network models, negative components in methods like principal components analysis (PCA) would complicate the interpretation of activation and inhibition interactions between gene programs.
3. Orthogonality: The orthogonality constraint makes it easier to interpret the data and aids model construction by forcing representations to be non-overlapping. Each gene program can be interpreted as a set of biological functions, and each cell is represented by a collection of biological function activities. Without orthogonality, each cell is likely to be represented by a single program containing all genes expressed by the same cell type. Further, in the context of D-SPIN, shared genes between gene programs would cause confounding interactions between the programs. Therefore, each gene program should be completely independent: in other words, orthogonal.

Given a non-negative gene matrix *X* ∈ ℝ^*C*×*M*^ of *C* cells and *M* genes and a program number *K*, oNMF solves the following optimization problem.

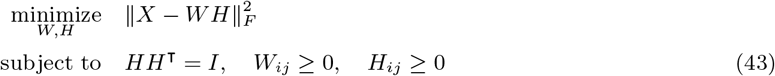

where *H* ∈ ℝ^*K*×*M*^ is the gene program representation, *W* ∈ ℝ^*C*×*K*^ is the cell state represented on the gene programs and 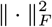 is the matrix Frobenius norm. We implemented the iterative algorithm proposed in this study [19], with *W* and *H* randomly initialized by uniform distribution on [0, 1]. The random initialization of *H* is orthogonalized using singular value decomposition (SVD) and takes absolute value before the start of iteration; we use 500 iterations in the computation by default.

According to the mathematical formulation of oNMF, the influence of each gene and cell state on the objective function is determined by the norm of the reconstruction error. To improve the discovery of gene programs, we utilized the following strategies to balance the contributions from each gene and cell state:

1. Negative binomial distribution gene CV filtering (Sec. 2.1). We noted that genes with high expression levels tended to have greater variance than typically expected under Poisson distribution CV filtering. Consequently, these genes often encode proteins involved in multiple housekeeping programs in the oNMF. Using a negative binomial distribution for CV filtering, we accounted for the higher variance of high-expression genes, enabling better gene selection. We also removed genes that are expressed in a tiny fraction of cells in gene filtering, as genes with close to 0 expression are not informative and risk introducing extra noise.
2. Gene normalization by standard deviation. Genes with high variance contribute more to the reconstruction error and can thus dominate the oNMF gene program discovery. To mitigate this, we divided the expression of each gene by its standard deviation across cells, ensuring that each gene contributed equally. The expression of genes with only a few nonzero entries may be disproportionately large after the scaling. Ideally, such genes should be filtered out during gene filtering, but they can also be removed by setting a cap on the scaling factor, for example, no more than a 5 times increase.
3. Cell state balancing by cell clusters. The gene program discovery will be dominated by reconstructing a specific cell state if that state constitutes a large proportion of the cell population. To address this, we used class-balance strategies frequently applied in machine learning practices. Specifically, we balanced different cell states by sub-sampling cells based on cell clusters obtained by clustering or gene markers. We employed two different schemes: (1) Equal-sample balancing: to identify gene programs that are only expressed in a small fraction of the cell population, we sampled an equal number of cells from each cell cluster; (2) Square-root balancing: in machine learning, taking samples from each data category proportional to the square root of the category size was found to be effective. Similarly, we took samples from each cell cluster proportional to the square root of the cell number in that cluster.

The exact solution of non-negative matrix factorization is an NP-hard problem [20]. But there are heuristic approximations of oNMF by iterative matrix update, and the solution varies with random initialization. Therefore, we run oNMF with different random seeds and compute a consensus gene program decomposition. For example, we take each row of *H* across different random seed repeats, and perform K-means clustering to compute the consensus composition of each gene program. The gene programs can be annotated with a combination of bioinformatics databases, including DAVID, Enrichr, and String-db, and also manual lookup [21, 22, 23].

### 1.10 Evaluating gene program expression and discretization

A gene program is defined as a set of genes that co-express across conditions in single-cell transcriptional profiling. Gene programs can either be identified through unsupervised learning techniques, such as oNMF (Sec. 1.9), or prior biological knowledge. The continuous program expression level corresponds to the cell-by-program matrix *W* . Although each oNMF run produces a *W* , after computing the consensus program and adding user-defined new programs, *H* is altered, and we need to re-evaluate *W* for the new gene sets of each gene program.

In the D-SPIN framework, the expression level of gene program *k* of cell *i* is described by the variable *S*_*ik*_ ∈ {−1, 0, 1}. To transform the expression matrix of genes *X* ∈ ℝ^*C*×*M*^ into the discretized program matrix *S* ∈ ℝ^*C*×*K*^, two steps are performed consecutively: computing a continuous weighted average of expression level *W*_*ik*_, and discretizing *W*_*ik*_ into three levels. For gene-level network construction, each gene is a unique program; thus, the first step is replaced by *W* = *X*, and the same discretization procedures in the second step can be applied.

#### Weighted average

To reduce noise and to synthesize the expression of all genes in the program, the continuous expression level across all cells ***w***_*k*_ = *W*_·,*k*_ should be a weighted average of genes in the program. By the consensus program in Sec. 1.9 and user-defined programs, we now have a mask matrix 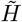 that records which program each gene belongs to

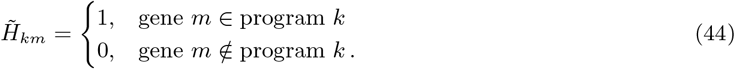

Note that 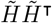 should be a diagonal matrix as each gene only belongs to a single program. With known gene assignments 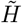, the objective function Eq. 43 can be split by each program and optimized individually as

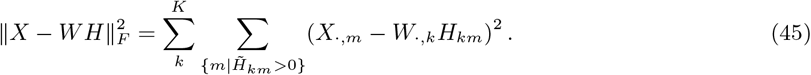

For every program *k*, we only need to consider genes that are known to belong to the program 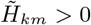. For notation simplicity, we denote *X*^(*k*)^ ∈ ℝ^*C*×*P*^ as the matrix of *P* genes that are known to belong to program *k, W* ^(*k*)^ ∈ ℝ^*C*×1^ as the cell expression level on program *k*, and *H*^(*k*)^ ∈ ℝ^1×*P*^ as the weights on genes in the program *k*. The optimization problem on program *k* is

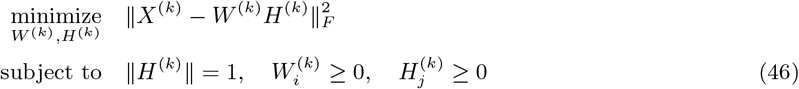

With only 1 component, the orthogonal constraints between programs become trivial. The problem is equivalent to a 1-component non-negative matrix factorization (NMF) problem and has mature implementations in Python scikit-learn packages.

Additionally, although the optimization problem in oNMF is nonlinear, the optimal solution corresponds to a linear transformation that applies to any new gene expression matrix. At the optimal solution, the partial derivatives regarding 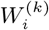 should be 0. Specifically,

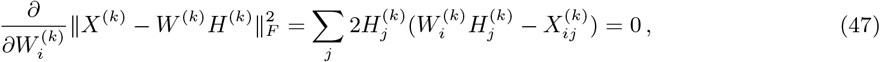

which implies

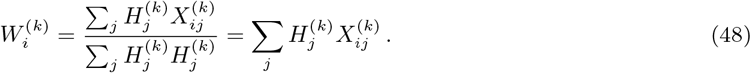

Generally, after computing the optimal program representation matrix *H*^*^ ∈ ℝ^*K*×*M*^ , the oNMF represents a linear transformation and for any new gene expression matrix *X*, the corresponding program representations of cell states are

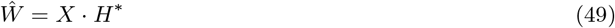

#### Discretization

Various methods have been empirically used to partition continuous expression into discrete levels, such as using percentiles or standard deviations. Discretization with K-means was found to perform well [24] and have a clear interpretation of minimizing the variance inside each expression level category. The 3-state K-means minimizes the following objective function for each gene program *k* across all cells indexed by *i*

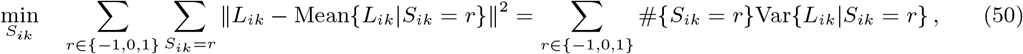

which is assigning −1, 0, 1 to each of *S*_*ik*_ so to minimize the variance of *L*_*ik*_ inside each group of cells classified by *S*_*ik*_. The K-means clustering also has mature implementations in the Python scikit-learn packages.

### 1.11 Selection of gene program number in oNMF

The selection of the number of gene programs is a trade-off between model expressive power, computational complexity, and model interpretability. The optimal choice of program number typically hinges on the specific requirements of the application scenario. In the context of single-cell transcriptional profiling, the number of gene programs is typically set to 10 ∼ 40, depending on the desired resolution of the gene matrix decomposition. To aid in this decision, various model selection criteria have been proposed to facilitate this choice, among which the Bayesian information criterion (BIC) and the elbow method are widely used examples.

BIC is derived from maximizing the model evidence *P* (Data|Model), where a penalty factor of the model dimension arises when integrating the parameter prior distribution over the whole parameter space [25]. Specifically, BIC is defined by maximized model log-likelihood 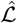 with a linear penalty term of model dimension.

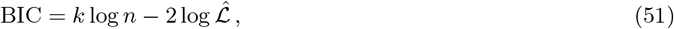

where *k* is the model dimension and *n* is the number of samples. Models with lower BIC are preferred.

As oNMF is not a probabilistic model by definition, BIC is not directly applicable. However, we can model the residue of the gene expression matrix after the oNMF fitting. In general, if the model error can be described as independent and identically distributed normal distribution variables,

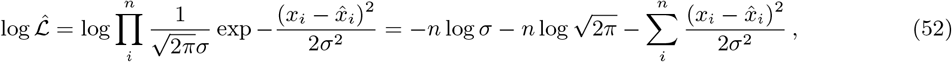

where *x*_*i*_ and 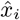 are data and model fitting, *σ* is the standard deviation of the error. As *σ* for a model is generally unknown and estimated from data, we have 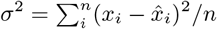. Therefore, the BIC is

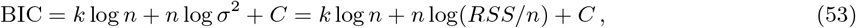

where *C* is a constant independent of model choice and 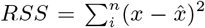 is the residual sum of squares. Motivated by this form, in oNMF we can use the matrix Frobenius norm of the fitting residual as *RSS*, number of cells as *n*, and number of programs as *k* to compute BIC to help to decide the number of gene programs.

Aside from statistical criteria like the BIC, the elbow method provides a heuristic approach to estimate the number of gene programs required in the oNMF. This method involves plotting a relevant cost function or objective function against the number of gene programs and looking for a point in the plot where the rate of change drastically alters, resembling an “elbow”.

Gene matrices of single-cell profiling are known to be especially noisy; therefore, traditional metrics such as model-explained variance may not be effective in revealing the elbow point. Instead, we use the number of genes that significantly correlate with corresponding gene programs as a more robust objective function. To consider the effect of noise, we can evaluate the pairwise gene-gene correlation as a reference distribution, which has a standard deviation in the order of 0.05 in our datasets. The correlation threshold between genes and programs can be set as 3*σ* or 4*σ* of the gene-gene correlations. Additionally, various methods (including MAGIC) have been proposed to denoise the gene expression matrix [1]. After denoising, both gene-gene correlation and gene-program correlation drastically increase, allowing us to use a high correlation threshold, such as 0.5. These refined approaches would facilitate the decision of gene program numbers in D-SPIN.

### 1.12 High-throughput drug perturbation experiments on PBMCs

All high-throughput drug perturbation experiments were performed in cell repellent-treated 96-well plates (Greiner Bio-One 07-000-640). We selected the drug library from the SelleckChem Immunology and Inflammation-related library (L4100), or the FDA-approved drug library (L2000), which were supplied at a stock concentration of 10 mM. The layout for drugs and/or drug combinations for each 96-well plate was predefined before starting the experiment. The final working concentrations for drugs varied between 0.01 nM to 10 uM. To stimulate PBMCs into an activated state, some wells included the use of ImmunoCult Human CD3/CD28 T Cell Activator (Stemcell Technologies 10971), which was used at the commercially recommended final concentration of 5 uL/200uL. After completing drug dilutions, we dispensed drugs into a 96-well plate layout, loaded 200,000 PBMCs into each well, and cultured for 24 hours in a tissue culture incubator at 37 °C and 5% CO2. All working drug dilutions and cell suspensions were prepared in RPMI-1640 + 1% Pen/strep + 10% FBS. The final concentration of DMSO within these experiments was always below 0.1%, which has been shown to produce minimal effects on immune cell activation [26].

### 1.13 High-throughput single-cell profiling using sample multiplexing techniques

After 24 hours of drug exposure, we harvested cells for single-cell profiling. Cells were enzymatically dissociated into a single-cell suspension and multiplexed using MULTI-seq lipid-modified oligos [2]. One experiment (MULT-19) was performed using the 10X Chromium Cellplex Kit (1000261), and manufacturer protocols were followed. Although plates were treated to be cell repellent, activation of certain cell types within the PBMC mixture can produce strong adhesive tendencies to the plate and each other. Thus, we enzymatically digested both the cell suspension and the plate surface with 100 uL of TrypLE (Thermo Fisher 12604021) to ensure the complete recovery of all cells. Briefly, the digestion was performed as follows: Using a multichannel pipette, all cells from each plate were transferred to a conical V-bottom 96-well plate (Thermo Fisher 249935), and then 100 uL of TrypLE was added to each well of the original plate and kept at room temperature. The conical V-bottom well plate, which contains the cells originally in suspension, was centrifuged at 400 rcf for 4 minutes. The supernatant was removed, and 100 uL of TrypLE was added to each well of pelleted cells and triturated gently. Both the original plate and the V-bottom plate were then incubated at 37 °C for 2 minutes simultaneously. After checking that any adherent cells were removed from the original plate, the cells in each plate were triturated and pooled back together. Cells were then labeled with MULTI-seq lipid-modified oligos according to the original protocol [2] and profiled using the 10x Chromium 3’ chemistry (v3.1, PN-1000121) or using the HT version (HT Kit v3.1, 1000370). Cells were super-loaded onto each lane of a Chromium chip at 25,000 - 40,000 cells per lane, or onto a Chromium HT chip at 60,000 cells/lane. Sample tags were purified and amplified according to the MULTI-seq procedure or according to manufacturer’s instructions for Cellplex. Libraries were sequenced using Illumina Novaseq S4 chips at the UCSF CAT Sequencing facility to a target depth of 40,000 transcriptome reads/cell barcode and 5000 sample tag reads/cell barcode. Cells that were multiply labeled by sample tags were computationally removed from the analysis.

### 1.14 Time-course profiling of T cell-mediated immune activation

A reverse time course of CD3/CD28 T Cell Activator was performed using a Tecan Evo robot (Caltech Protein Expression Center). Briefly, cryopreserved PBMCs from healthy donors were thawed and rested for 16 hours as described above. Cells were then pelleted and dissociated with gentle trituration and seeded into 96-well plates at 200,000 cells/well in 150 uL of RPMI-1640 + 10% FBS + 1% P/S. The cells were then brought to a Tecan Freedom Evo2 liquid handling robot, which was programmed to dispense CD3/CD28 activator into each well at 30-minute intervals for 30 hours. The CD3/CD28 activator was diluted 1:10 in RPMI-1640 + 1% Pen/strep + 10% FBS, and 50 uL of the dilution was added at each time point. One well that did not receive any activator served as timepoint 0. Between dispenses, the plate was held in an onboard incubator at 37 °C and 5% CO2. After the conclusion of the time course, cells were immediately harvested for single-cell profiling using multiplexing tags (MULTI-seq) as described above.

### 1.15 Alignment and de-multiplexing

We used the Cell Ranger software (10x Genomics) to align single-cell RNA sequencing reads to the transcriptome. For experiment batches MULT-6, 7, 8, 9, 10, 12, 13, 14, 15, 16, TC, we used Cell Ranger 3.1.0. For experiment batches MULT-17, and 18, we used Cell Ranger 5.0.1. For experiment batches MULT-19 and MULT-20, we used Cell Ranger 6.1.2. The 10x Genomics hg19-1.2.0 genome build was used as a reference transcriptome.

Except for MULT-19, where we used the 10X Cellplex pipeline to demultiplex the conditions of each cell, we used a custom Python pipeline to assign cells to each condition based on the sequenced tags following the pipeline in MULTI-seq [2].

## 2 Quantification and statistical analysis

### 2.1 Filtering, normalization, and clustering of the Perturb-seq dataset

The Perturb-seq dataset is a genome-wide screening study where every single gene is knocked down individually [18]. In Perturb-seq, a library of guide RNAs is introduced by lentiviral transduction into engineered cells that express the dCAS9-KRAB CRISPRi effector, after which single RNA sequencing (scRNA-seq) is conducted to simultaneously read out the transcription profile and guide RNA for each cell. Because guide RNA transfection is a stochastic process, the number of cells receiving each perturbation varies vastly. Here, the dataset contains perturbations with a sample size ranging from 2 to 2,000 cells. To reduce noise introduced by the small sample size, we excluded perturbations with less than 20 cells (metadata entry “number of cells (filtered)”), which contained 92 perturbations, 0.86% of all perturbations.

More than half of the perturbations do not yield significant single gene-level expression changes (apart from the perturbed gene), possibly due to genome redundancies that ensure the robustness of cellular activities, and that some genes may not function in the profiled experimental condition. To avoid the model overfitting the measurement noise on these perturbations with no significant effect, we excluded perturbations with less than 10 differentially expressed genes (metadata entry “Number of DEGs (anderson-darling)”), which contained 7,523 perturbations, 70.5% of all perturbations. Similarly, we only selected a subset of the 585 control samples with non-targeting guide RNAs to avoid overfitting on measurement noise. We selected control samples that were labeled as core control (metadata entry “core control”) and had no less than 200 collected cells (metadata entry “number of cells (filtered)”). In total, we selected 3,136 perturbation conditions and 105 control conditions.

Normalization of single-cell gene expression data is required because (1) the total number of transcripts captured for each cell can vary from 1,000 to 100,000 due to technical variability in reagents and library preparation steps, and (2) transcript expression levels across genes can span 5 orders of magnitude. We normalized the raw counts for each cell by the total counts detected in each cell, times a scaling factor that is roughly the median of the total counts of transcripts in a cell. We used the scaling factor 10^4^ as the target total count, that is, computing 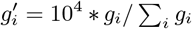 for every cell, where *g*_*i*_ is the count of gene *i*. We also used a log transformation to mitigate the great variety of expression levels across genes 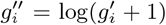. Except for highly variable gene filtering, all the following analysis was performed on the log-transformed data.

Due to measurement noise, genes that have constant expression across cells do not contribute to biological impact, but only increase noise in the data. Therefore, single-cell data analyses are performed on a subset of genes that have different expression levels across cells, i.e., highly variable genes. Here, we developed a highly variable gene filtering model adapted from the coefficient-of-variation (CV) filtering using the negative binomial distribution as the statistics of gene counts. The model was selected due to our observation that genes with high expression generally have higher CV than the prediction of the Poisson model. For a negative binomial distribution NB(*r, p*), the mean and CV of the distribution are

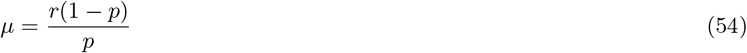

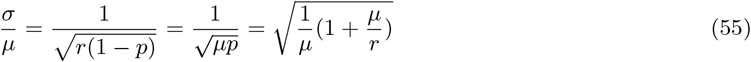

The negative binomial distribution suggests another type of log CV (*y*) - log mean (*x*) relationship where the expected log CV is higher than the prediction of a Poisson model for high-expression genes.

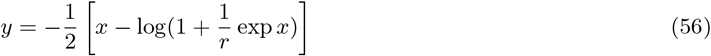

Therefore, we followed the form of the equation and obtained the empirical log CV - log mean relationship by curve fitting (scipy.optimize.curve fit) of the following function

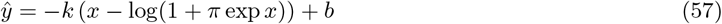

where we denoted 1*/r* as a parameter *π* for numerical stability. The genes are ranked by log CV − *ŷ*, the difference between log CV and curve fitting. A higher value indicates that the gene is more variable.

We selected highly-variable genes as the union of the following three sets: (1) we performed the Wilcoxon test of log-transformed data for each guide RNA and control (sc.tl.rank genes groups) and selected the top 10 differentially expressed genes for each guide that passed *p <* 0.05 under Bonferroni multi-test correction (we selected less than 10 if no enough genes pass the test), (2) the top 1,000 highly-variable genes selected by our negative binomial CV filtering, and (3) for comparison with the original Perturb-seq study, we included the genes that appeared in the gene programs in the original Perturb-seq study. In total, we selected 3,273 genes for the analysis using the D-SPIN framework.

To visualize and cluster single cells in the Perturb-seq dataset, we corrected the batch effect by constructing latent representations with scVI on raw gene counts (dispersion=‘gene-batch’, n latent=10, n layers=2, max epoch=20, batch size=2048) [27]. We constructed a 15-nearest-neighbor graph (scanpy.pp.neighbors) on the latent representation and computed Leiden clustering for cell clusters (scanpy.tl.leiden). As K562 is a purified cell line, it contains primarily a single cell type. Thus, the UMAP embedding of the cell population is a circular structure, representing the progression of the cell cycle, which is less relevant to the study of perturbation responses. Therefore, we only show the UMAP embedding on control samples along with D-SPIN-generated samples using cell-cycle-associated gene programs (Sec. 2.14).

### 2.2 Filtering, normalization, and clustering of the drug profiling dataset

The single-cell gene expression normalization in the drug profiling dataset is generally identical to the Perturb-seq dataset. We normalized the raw counts by each cell with the same scaling factor 10^4^, and gene expressions were log-transformed for the following analyses, except for the highly variable gene selection.

The highly variable gene selection was based on the negative binomial expression model we developed (Sec. 2.1). The drug profiling dataset contains (1) a large dataset of single drug and drug combination profiling (Drug Profiling) and (2) a dataset of follow-up drug dosage combinations (Dosage Combination). The gene filtering was performed on the Drug Profiling dataset, and Dosage Combination follows the same filtering. To avoid noise in low-expression genes, we excluded genes that only expressed in less than 0.1% of the cells in the Drug Profiling dataset. (In the Perturb-seq dataset, the low-expression genes were already filtered out by the authors in the published data). We selected 3,287 highly variable genes in the remaining genes, which is log 10(1.1) above the curve fitting Eq. 57 for the negative binomial distribution CV model.

Collected single-cell profiling data have batch effects across experiments, likely due to variations in cell culture conditions and technical differences across sequencing machine runs. To visualize, cluster, and cell-typing single-cells in the Drug Profiling dataset, we corrected batch effects by constructing latent representations on raw gene counts of 5,000 selected genes (scanpy.pp.highly variable genes) with scVI (dispersion=‘gene-batch’, n latent=10, n layers=2, max epoch=200, batch size=5120) [27]. We constructed a 15-nearest-neighbor graph (scanpy.pp.neighbors) on the latent representation, computed Leiden clustering for cell clusters (scanpy.tl.leiden), and conducted UMAP embedding for visualization (scanpy.tl.umap). The cell subtype identities of each Leiden cluster were annotated manually by comparing the most significantly up-regulated genes of the cluster with the remaining cells. We removed 3 clusters that mostly came from the experimental batch “MULT-6” and were putatively cell debris, as they expressed the maternal effect gene OOEP and disrupted transcriptional states were present. We also excluded red blood cells (RBC) and plasma B cells for D-SPIN analysis, as they are small and isolated cell populations with distinct transcriptional profiles.

The Dosage Combination dataset followed the same normalization and log-transformation pre-processing and used the same set of highly variable genes as identified from the Drug Profiling dataset. For direct comparison with the Drug Profiling dataset, we computed the leiden clusters and UMAP embedding of the Dosage Combination dataset by homogenizing with the Drug Profiling dataset. Specifically, batch MULT-18 in Drug Profiling contains drug combination experiments and has the most similar setup to the new Dosage Combination dataset. We trained a scVI batch correction model with MULT-18 and the Dosage Combination dataset to obtain a joint embedding of both datasets (dispersion=‘gene-batch’, n latent=10, n layers=2, max epochs=100, batch size=128). For each cell in the Dosage Combination dataset, we found the 10 nearest neighbors in the MULT-18 dataset. The cell is then assigned to the most frequent Leiden cluster among these 10 neighbors, and has UMAP coordinates as the average of the neighbors with the same Leiden cluster label.

### 2.3 Constructing D-SPIN models for the Perturb-seq dataset

The analysis of a single-cell perturbation dataset with D-SPIN contains four major components.

1. Identifying gene program using oNMF or prior biological knowledge.
2. Inferring program-level regulatory network model and perturbation responses.
3. Inferring gene-level regulatory network model and perturbation responses.
4. Associating the program-level and gene-level network models to find gene regulators The detailed procedures for the Perturb-seq dataset are as follows.

#### Gene program discovery

In the Perturb-seq study, we used a subset of 10^5^ single cells, with the Square-root balancing scheme in Sec. 1.9 for the Leiden cluster of cell states. We ran oNMF 100 times with *K* = 30 with different random seeds to solve Eq. 43 and computed a consensus gene program set by K-means as detailed in Sec. 1.9. The identified gene programs are functionally annotated through gene ontology annotation tools and manual lookup [21, 22].

#### Program-level network model inference

In the Perturb-seq study, a large proportion of perturbations induce insignificant effects on the gene program level. To avoid overfitting on measurement noise, we selected a subset of 128 samples that were most different from each other (Sec. 1.1). The subset of samples was selected by clustering all samples into 128 groups by their cross-correlation and taking one sample from each group. We then learned the response vectors of all other samples independently conditioned on the inferred network.

We used MCMC maximum likelihood inference for the 30-node program-level networks. In MCMC estimation of the gradient in Eq. 4 and Eq. 5, we used 5 × 10^7^ samples in each iteration for each condition, with 10^5^ samples for warm-up for network inference and 10^7^ samples for perturbation response vector inference. We used *ℓ*_1_ regularization on the network with *λ*_1_ = 0.01, promoting the sparsity and interpretability of the network. We used *ℓ*_2_ regularization on the response vectors with *λ*_2_ = 0.005 to avoid an excessively large magnitude of response with strong activations and inhibitions. In optimization, we used the Adam optimizer with a base learning rate of 0.02.

#### Gene-level network model inference

We selected candidate regulatory genes as transcription factors (TFs) that were expressed in over 4% of cells and also had effective knockdown samples as the criteria in Sec. 2.11. In total, we selected 469 TFs and constructed the gene-level regulatory network on the 469 knockdown conditions. Additionally, we incorporated 46 control conditions with non-targeting guide RNA, with a minimum of 250 cells each.

We used the pseudolikelihood algorithm to compute the gradient for a directed network model as Eq. 11 for inference with the Adam optimizer and a base learning rate of 0.05. We used *ℓ*_1_ regularization on the network with *λ*_1_ = 0.01. As we knew the target of each knockdown perturbation, we used the prior 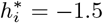 for the knockdown target genes, and perturbation prior regularization with *λ*_2_ = 3.

#### Identifying gene regulators for programs

For a comparable evaluation with the results in the original Perturb-seq study, we used the gene program of erythroid and myeloid differentiation of the Perturb-seq study [18]. To avoid confounding interactions, TFs were removed from the gene programs to ensure the identified interactions are not caused by shared genes. The expression level evaluation and discretization of the erythroid/myeloid differentiation program follows the standard procedure in Sec. 1.10. The regulators from the gene-level network were identified by solving Eq. 14 under *ℓ*_1_ regularization with *λ*_1_ = 0.005.

For comparison, the z-score of differentiation programs is evaluated on the continuous program expression value before discretization. The z-score of all TF perturbation conditions is normalized relative to the mean and standard deviation of these scores in non-targeting control conditions.

### 2.4 Constructing D-SPIN models for the drug profiling dataset

The detailed procedures for the drug profiling dataset are as follows.

#### Gene program discovery

In the drug profiling study, we used a subset of 3.3 × 10^5^ single cells using the equal-balancing scheme, sampling 1,000 single cells from each Leiden cluster of cell states. The sampling scheme was to better capture gene programs in drug-specific cell states that composed a small proportion of all the cells. As specific drug-induced cell states were a small fraction of the whole dataset, we increased the resolution of oNMF to *K* = 50 and solve for Eq. 43. During preliminary runs of oNMF, we identified drug-specific response programs such as pathogen response and stress response. We then manually curated random seeding to select the oNMF decomposition that included these target gene programs.

We combined the 50 programs into 30 by merging programs with similar biological functions and distributions of cell subtype expression. The program function was annotated by a combination of informatics databases and manual lookup [21, 22, 23] and subsequently validated on a single-cell atlas of human immune cells [28].

#### Program-level network model inference

We infer the regulatory network with all 1,132 experimental conditions in the Drug Profiling dataset. The computational setup was generally identical to the program-level network inference in the Perturb-seq dataset. We use MCMC estimation of the gradient in Eq. 4 and Eq. 5 with the Adam optimizer and a base learning rate of 0.02. We used *ℓ*_1_ regularization on the network with *λ*_1_ = 0.01, and *ℓ*_2_ regularization on the response vectors with *λ*_2_ = 0.005

#### Gene-level network model inference

During immune activations, the signal transductions by protein phosphorylation and dephosphorylation also play important roles in addition to TF-controlled transcriptional regulation. Therefore, we included TFs, kinases, and phosphatases in the network construction. We selected TFs that expressed in over 5% of cells, and kinases and phosphatases that expressed in over 2% of cells. In total, we selected 657 regulatory genes for the network construction, including 388 TFs, 187 kinases, and 71 phosphatases. We use all 1,132 experimental conditions for network construction.

The network construction followed the same procedure as the gene-level network inference in the Perturb-seq dataset. We used the pseudolikelihood algorithm to compute the gradient for a directed network model as Eq. 11 for inference with the Adam optimizer and a base learning rate of 0.05. We used *ℓ*_1_ regularization on the network with *λ*_1_ = 0.01, and *ℓ*_2_ regularization on the response vectors with *λ*_2_ = 0.005

#### Identifying gene regulators for programs

The regulators from the gene-level network were identified by solving Eq. 14 for all of 1,132 conditions in the dataset under *ℓ*_1_ regularization with *λ*_1_ = 0.005.

### 2.5 Numerical optimization and parallelization of D-SPIN

#### Numerical optimization

In optimization, we used the Adam optimizer with a learning rate of 0.2 for exact maximum likelihood, 0.02 for MCMC maximum likelihood, and 0.05 for pseudolikelihood. The decay rate was the default of Adam optimizer as *β*_1_ = 0.9, *β*_2_ = 0.999, *ϵ* = 10^−6^. Adam optimizer, as a first-order method, can significantly speed up the convergence rate of optimization, but it suffers from instability during the final stage of optimization. We therefore deploy a backtracking scheme where we roll back 20 iterations and half the step size when the current gradient is larger than twice the gradient 20 iterations before, and stop the training after 5 backtracking operations. We used 800 learning epochs if the algorithm did not early stop with backtracking, and selected the network with the smallest gradient in the learning trajectories. The combination of the Adam optimizer and backtracking schemes made the training process robust with the base learning rate and the maximum number of epochs, as the algorithm will adaptively reduce the step size under unstable training and stop training when the loss curve has stabilized.

#### Parallelization of D-SPIN inference

An advantage of D-SPIN is that we can parcel out computing jobs to different CPUs. For instance, the gradient estimation of each summation term in Eq. 4 and each condition in Eq. 5 is independent of the others. Leveraging the property, D-SPIN performs parallelized model inference by parceling computation for each condition to different CPUs. We use the MATLAB “parfor” function on the Caltech Resnick cluster, with each task taking 2 cores and 32 GB of memory. The inference of response vector ***h*** given on a fixed network ***J*** is also independent for each condition and is parallelized with the Slurm Workload Manager system on the Caltech Resnick cluster with 2 cores and 32 GB of memory.

### 2.6 Simulations of the four-pathway model and the HSC network

The BEELINE framework provides a systematic tool to simulate single-gene expression data from Boolean regulatory networks and to benchmark the accuracy of network inference methods [29]. Our simulation and evaluation mostly followed the formulation and code of BEELINE, but we complemented the framework with the simulation of gene knockdown and activation perturbations. BEELINE simulates the following stochastic differential equation (SDE) models of gene expression.

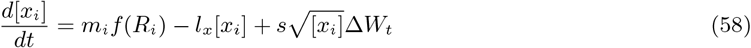

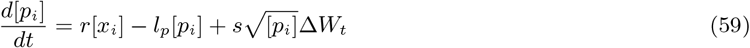

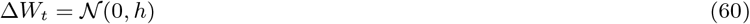

where [*x*_*i*_] and [*p*_*i*_] are mRNA and protein concentration of the gene *i, m*_*i*_ and *r* is the mRNA and protein production rate, *f* (*R*_*i*_) is the regulatory term computed from the boolean network by BEELINE, *l*_*x*_ and *l*_*p*_ are mRNA and protein degradation rate, *s* is thermal noise strength, *W*_*t*_ is the Wiener process representing thermal noises, and *h* is simulation time step. The default parameter values are *m*_*i*_ = 20, *l*_*x*_ = 10, *r* = 10, *l*_*p*_ = 1, *s* = 20. At *t* = 0 all mRNA concentrations [*x*_*i*_] are initialized as 1, and all protein concentrations [*p*_*i*_] are initialized as 0. Specifically, the mRNA default initial concentration 1 in the current BEELINE framework (v1.0) is different from 0.01 used in the version (v0.1) presented by the BEELINE paper [29], possibly to improve numerical stability and to avoid the system being trapped by the all-0s attractor state.

To model the impact of gene knockdown and activation, we included two extra terms into the dynamics of mRNA: *e*_*i*_, representing a constant external mRNA production not regulated by the network, and *β*_*i*_, representing the proportion that the mRNA production rate is suppressed in addition to the network regulation. After the modification, the dynamics of the mRNA are

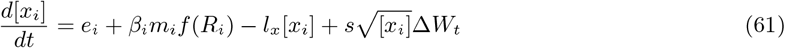

Without perturbation, we have *e*_*i*_ = 0, *β*_*i*_ = 1.

In the four-pathway model, as an illustrative example, we used strong perturbations with *β*_*i*_ = 0 for simulating gene knockdown and *e*_*i*_ = 40 for simulating gene activation. In the HSC network, we used *β*_*i*_ = 0.2 knockdown and *e*_*i*_ = 15 for activation, which corresponds to 0.75 of the expression of fully activated genes. We used weaker perturbations to HSC networks to avoid interrupting the global cell differentiation trajectories. We simulated 2,000 cells for control conditions and 500 cells for each perturbation condition.

We simulated both network models under 10 different sets of randomly sampled parameters. Each time the parameters *m*_*i*_, *l*_*x*_, *l*_*p*_, *r* were assigned by the formula *αD*, where *D* represents the default value of the parameter and *α* is a random factor sampled from a truncated Gaussian distribution (mean = 1, std = 0.1, truncation to [0.9, 1.1]). By default, the BEELINE framework sets the same type of parameters for all genes to the same value for numerical stability. In the simulated HSC network data, we clustered the cell states and computed the UMAP embedding of the discretized simulated data with D-SPIN-generated samples for visualization, and annotated each cluster based on the research that proposed the HSC network model [9].

### 2.7 Clamping analysis of node marginal distribution and edge sensitivity

As a generative model of transcriptional state distribution, D-SPIN allows us to quantify how a perturbation alters the marginal distributions of any node in the network and to decompose this effect into contributions from individual edges. This procedure allows us to identify critical interactions that determine a given perturbation response and to make quantitative statements about how the regulatory network controls cell state distributions under perturbation.

To quantify the marginal distribution change of each node *i*, we define an effective marginal activity *ĥ*_*i*_(***J*** , ***h***) by the ratio between the node being on (*s*_*i*_ = 1) and the node being off (*s*_*i*_ = −1) in the marginal distribution *P* (*s*_*i*_|***J*** , ***h***)

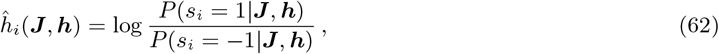

where positive *ĥ*_*i*_ indicates the node being activated and negative *ĥ*_*i*_ indicates the node being repressed. The definition is rationalized by parametrizing the complete three-state marginal distribution of node *i* by an effective 1-node D-SPIN model

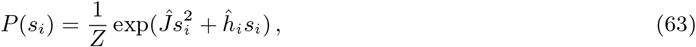

whose analytical solution is 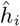, and we drop the constant coefficient for simplicity. As *ĥ*_*i*_ is defined as a linear term inside the energy function, the difference of *ĥ*_*i*_ between conditions represents the relative activation or inhibition of a node.

To quantify the contribution of each edge to the marginal distribution change, we compare the marginal distribution with and without the edge of interest. The edge sensitivity of *J*_*kl*_ to node *i* is defined as the marginal activity *ĥ*_*i*_ change after removing the edge *J*_*kl*_ from the network. Specifically, we define a clamped network 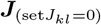 where *J*_*kl*_ is set to 0 while all other edges remain the same, and compute the marginal activity 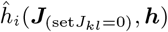 of node *i*. The edge sensitivity *ϵ*(*kl, i*) is thus defined by

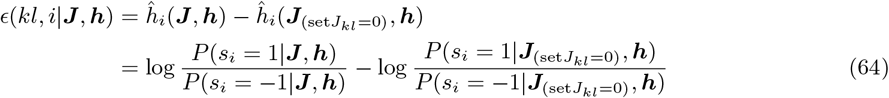

With the HSC network example, we demonstrate how this procedure identifies the impact of PU.1 knock-down on the node Gfi-1, and which network edge contributes to the change of Gfi-1. D-SPIN infers the network ***J*** , response vector ***h***_0_ without perturbation and ***h*** under PU.1 knock-down. Without perturbation, by marginalizing the full distribution *P* (***s***|***J*** , ***h***_0_), we have

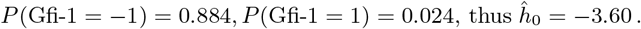

Similarly, under PU.1 knock-down, marginalizing the complete distribution *P* (***s***|***J*** , ***h***_0_) gives

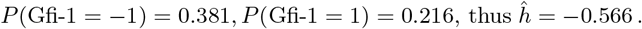

Therefore, PU.1 knockdown has an overall impact of 3.04 activation on Gfi-1. For edge sensitivity, take the edge Gfi-1-EgrNab, for example. After clamping the edge to 0, the complete distribution *P* (***s***|***J***_(set Gfi-1-EgrNab = 0)_, ***h***) has an marginal distribution of

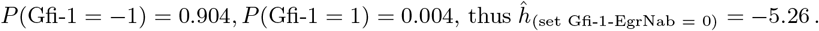

Therefore the edge sensitivity *ϵ*(Gfi-1-EgrNab, Gfi-1) = *ĥ* − *ĥ*_(*set*Gfi-1-EgrNab=0)_ = 4.69. The edge Gfi-1-EgrNab has a high contribution in activating the node Gfi-1. The contribution of other edges can be computed similarly, and the top contributing edges to Gfi-1 under PU.1 knock-down are Gfi-1-EgrNab 4.69, CEBPA-Gfi-1 -2.85, Gfi-1-PU.1 -2.60, and Gfi-1-cJun 2.31.

### 2.8 Simulation of large synthetic networks

To assess the accuracy and scalability of D-SPIN, we constructed three distinct sets of large regulatory network models, each comprising networks with 125, 250, 500, and 1000 nodes. These models represent three major categories of networks:

1. Random networks: One of the most representative random networks is Erdős–Rényi models (ER models). These networks are completely random in nature, where each node has the same probability of interacting with any other node.
2. Scale-free networks: In these networks, the number of interactions for each node follows a power-law distribution, which means there are some hub nodes that have a significantly higher number of interactions. Some biological networks, such as metabolic networks, exhibit such statistics.
3. Modular networks: These networks are organized into modules, characterized by activating interactions inside each module and inhibitory interactions between modules. Consequently, gene expression of these networks also exhibits a modular pattern, where genes within the same module are more likely to be activated at the same time.

The construction of a random network is straightforward. To ensure the same average degree of network nodes across different network sizes, each node has a 1.25*/M* probability to activate any other nodes and a 1.25*/M* probability to inhibit any other nodes, where *M* is the number of nodes in the network. We ensure the network is fully connected by regenerating the network with a different random seed when the network has a disconnected structure.

The scale-free networks are constructed by the canonical rules of preferential attachment. Starting from two connected nodes, a new node is added to the network and selects two existing nodes as its regulators. The selection probability is proportional to *k* + *k*_0_, where *k* is the current out-degree of the existing nodes and *k*_0_ = 2 is a constant; therefore, nodes with higher degrees are more likely to be selected to connect with the new nodes. The iteration continues until the target number of nodes is reached. Hub nodes with at least 10 outgoing connections have extra self-activations to ensure the dynamics of the entire network do not die out and stay at all 0s.

In our construction of modular networks, the number of modules in the networks is proportional to the number of nodes. The 125, 250, 500, and 1000-node networks contain, respectively, 2, 4, 8, and 16 modules. The exact number of nodes in each module is determined by sampling Poisson random numbers, with the distribution expectation being the expected module size. Within each module, we generate a subnetwork using the same algorithm as generating scale-free networks. Nodes with no less than 2 outgoing connections are labeled as hub nodes and assigned self-activation to ensure the dynamics do not die out. Between different modules, each hub node randomly selects 2 other hub nodes from other modules as targets of inhibitory interaction.

Using these synthetic networks, we simulated the dynamics of each model with our modified BEELINE framework. Given the complexity and scope limitations of designing combinatorial perturbations on large networks, we evaluated 6 different heuristic strategies for perturbation design and evaluated D-SPIN’s accuracy of network inference on 125-node networks with all three network topologies (Figure. S2(E)). Among the tested strategies, the most informative perturbations are random perturbations, where each node was subject to a random perturbation following a truncated Gaussian distribution. Specifically, for each node *i*, the perturbation 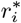 was taken from a Gaussian distribution with a mean of 0, a standard deviation of 0.8, and truncated to [−1, 1]. These perturbations dictated the biochemical parameters in the simulation by

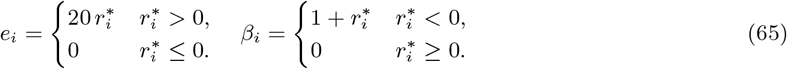

For a perturbation 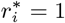, the strength of external activation is the same as the strongest internal expression in the network. For a perturbation 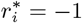, the gene *i* is completely deactivated. Therefore, this perturbation assignment covered a spectrum of perturbation actions for assessing the capability of D-SPIN in constructing regulatory networks from perturbations.

### 2.9 Benchmarking time consumption with cell number

To assess the scalability of each method relative to cell number, we performed time benchmarks for D-SPIN together with PIDC, GENIE3, and GRNBoost2, using the dataset of 10 different random repeats of 250-node Erdős–Rényi networks from Sec. 2.8. Specifically for D-SPIN, the gradient evaluations were performed in batches for efficient computation, with each batch containing 2,000 cells. For PIDC, GENIE3, and GRNBoost2, we used the Docker images of the three methods provided by the BEELINE framework. The analyses were performed on the Caltech Resnick cluster, with each task taking 2 cores and 48 GB of memory. The longest running time of each task was 7 days. The time of unfinished tasks was interpolated using the running time of the same methods with smaller cell numbers using the function *y* = *Ax*^*B*^ + *C* in the log-log space.

### 2.10 Benchmarking network inference accuracy with synthetic networks

We performed network inference benchmarking in 4 different setups.

1. Directed network inference for the four-pathway model
2. Undirected network inference for HSC network
3. Directed network inference for HSC network
4. Directed network inference for large-scale networks with three different topologies.

For D-SPIN, we used exact maximum likelihood inference for the undirected HSC network and the pseudo-likelihood method for all directed network inference. For both undirected and directed network inference of the HSC network, we use no regularization to examine the performance of D-SPIN without any prior knowledge of the dataset. For the directed network inference of the four-pathway model and large-scale networks, we used *ℓ*_1_ regularization on the network with *λ*_1_ = 0.01, following our default setting of network inference. As the perturbation is known, we applied a prior on the perturbation response as described in Sec. 1.5. Note that the perturbation actions were changing the coefficient of stochastic differential equation (SDE) models of biochemical reactions (Sec. 2.8), which is a totally different type of model from the probabilistic graphical models in D-SPIN. Nonetheless, the information about the perturbations could still assist the network inference.

Specifically, to determine the strength of the regularization factor *λ*_2_ and a scaling factor *α* that converts the perturbation *r*^(*n*)^ to the SDE to a relative response *h*^(*n*)^ = *αr*^(*n*)^, we examined the inference accuracy of D-SPIN with these parameters for a small subset of data, 125-node networks with 100 perturbations. We found that in the examined range, the accuracy monotonically increases with these parameters and saturates at around *λ*_2_ = 2 and *α* = 3. The results suggest that a stronger prior on the perturbation is generally beneficial for the accuracy of D-SPIN. We therefore use *λ*_2_ = 2 and *α* = 3 as the parameters for regularization.

The linear regression model is derived from modeling the gene expression by a multidimensional Gaussian distribution *X* ∼ 𝒩 (*µ* + *Wr*, Σ). The coefficient matrix *W* between the perturbation input and *r* and the average response ⟨*X*⟩ can be viewed as a regulatory network model, which can be solved by linear regression (sklearn.linear model.LinearRegression).

To compare the network reconstruction accuracy with other methods, including PIDC, GRNBoost2, and GENIE3, we ran the Docker images of the three methods provided by the BEELINE framework [29]. As the three methods do not explicitly model the impact of perturbations, we intended to run these methods on the full single-cell dataset with the perturbation label removed. However, as evaluated by Sec. 2.9, these methods have significantly increased computation time with datasets of large cell numbers, and cannot finish within days or even weeks.

Therefore, we ran these methods on the full single-cell datasets for the smaller datasets, including the four-pathway model and the HSC network. For inference in the large-scale networks, we reduced the total cell number by grouping *κ* similar cells from the same condition into a pseudobulk cell or metacell. We chose *κ* = 10 for evaluating these methods on large networks with 800 total perturbations, which is the minimal *κ* that allows these methods to finish in 1 day with 16 CPU cores. The only exceptions are GRNBoost2 on 1,000-node networks, which cannot finish in 1 day even with 64 cores under *κ* = 10, therefore we used *κ* = 20 for these cases. Specifically, for each condition (2,000 cells in control or 500 cells in perturbed conditions), we performed K-Means clustering with cluster number as *N/κ, N* being the cell number in the condition. We averaged the gene expression for each cluster and combined them into a pseudobulk dataset.

All of the three methods, PIDC, GRNBoost2, and GENIE3, are classified as not requiring hyperparameters in the BEELINE benchmark framework [29]. PIDC constructs the network by estimating mutual information between genes, which requires partitioning gene expression into bins for counting joint distribution. PIDC develops an adaptive Bayesian discretizer that automatically determines the number of blocks to partition, therefore requiring no input parameters. Both GENIE3 and GRNBoost2 (both provided by the Python package arboreto) build regression models for each gene to identify predictors as gene regulators. GRNBoost2 uses Stochastic Gradient Boosting Machine (SGBM), and GENIE3 uses Random Forests (RF). Regression models are generally robust to hyperparameters, and the package has determined a set of fixed hyperparameters and does not expose these parameters to the user interface. Given that, we followed the code examples and the Docker images of the three methods provided by the BEELINE framework [29].

### 2.11 Evaluating gene network reconstruction on biological datasets

To further assess the performance of D-SPIN on regulatory network reconstruction with biological measurement datasets, we applied different network inference methods to the genome-wide Perturb-seq dataset and compared the results with biological validations obtained from the ChIP-seq database. Chip-seq data measure the binding of transcription factors (TFs) to target gene promoters and have been used in the literature to assess the quality of inferred gene regulatory network models. We evaluate the correspondence between the inferred directed network and ChIP-seq data by the number of inferred edges that can be associated with a measured TF-target gene binding event.

Specifically, we generated three datasets from the Perturb-seq data, similar to the BEELINE framework [29], labeled as TFs+500HVGs, TFs+1000HVGs, and TF+1500HVGs. We selected TFs that expressed in over 5% of cells and the top 500, 1000, and 1500 highly variable genes (HVGs) that were identified with the negative binomial distribution model we developed (Sec. 2.1). We only included genes that had corresponding knockdown perturbation conditions that satisfied the following criteria.

1. More than 20 cells collected (metadata entry “number of cells (filtered)”).
2. More than 2 differentially expressed genes (DEGs) identified (metadata entry “Number of DEGs (anderson-darling)”).
3. A knockdown efficiency exceeding 5% (metadata entry “percent knockdown”).

Additionally, we incorporated 46 control conditions with non-targeting guide RNA, with a minimum of 250 cells each. After filtering, the TFs+500HVGs dataset comprised 84 cells, 335 genes, and 381 conditions; the TFs+1000HVGs dataset included 124k cells, 506 genes, and 552 conditions; the TFs+1500HVGs dataset included 157k cells, 663 genes, and 709 conditions.

For existing network inference methods, PIDC, GENIE3, and GRNBoost2 were directly evaluated on the entire dataset with the perturbation label removed. As previously benchmarked in Sec. 2.9, these methods do not scale well with large datasets exceeding 100k cells and took multiple days on 64 CPU cores to complete. We ran CellOracle (Version 0.20.0) on the entire dataset following the document instructions. We use the pre-built human promoter base GRN provided by CellOracle for network inference with prior knowledge, as the genome-wide Perturb-seq data did not have scATAC-seq measurement. We used a fully connected graph as the base GRN to quantify the accuracy of CellOracle without prior knowledge. For the differential expression (DE) / z-score method, regulatory interactions A to B are computed as the gene expression change (z-score) of gene B under the knockdown of gene A [30, 31].

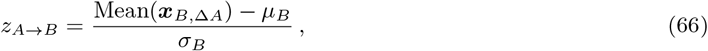

where *µ*_*B*_ , *σ*_*B*_ are the mean and standard deviation of gene B across all cells, and ***x***_*B*,Δ*A*_ are gene B expression under gene A knockdown.

For D-SPIN, we partitioned the data based on each perturbation condition. D-SPIN can handle perturbations with small cell numbers as information is shared through inferring a unified regulatory network. We used pseudolikelihood inference because of the large network size. As we knew the target of each knockdown perturbation, we used the prior 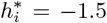 for the knockdown target genes, and perturbation prior regularization with *λ*_2_ = 3. With the pre-built promoter base GRN from CellOracle as prior knowledge, we applied *ℓ*_1_ regularization on edges not belonging to the base network with *λ*_1_ = 0.02. Without the base network, we regularized all edges in the network with *λ*_1_ = 0.01.

The inferred networks were compared with processed ChIP-seq data of TF-target interactions from human ChIP-seq datasets. We used the interactions provided by the BEELINE framework, which were integrated from multiple databases [29]. As the ChIP-seq data only contained TF-target regulations, we removed all inferred interactions with non-TF regulators, which was consistent with the BEELINE framework. The correspondence with ChIP-seq data is quantified by the early precision rate (EPR), which measures the proportion of inferred edges that can be associated with a TF binding event detected by ChIP-seq among the top *K/*4 predictions, relative to the expectation of a random predictor. *K* is set to the total number of TF-target interactions in the ChIP-seq database. We also quantified the inference quality by AUROC (Area Under the Receiver Operating Characteristic Curve), which provides a global metric of the model’s performance on ranking positive examples higher than negative examples.

Additionally, we also evaluated the network reconstruction accuracy of D-SPIN and CellOracle on the entire drug profiling dataset, as other methods cannot scale to datasets with more than a million cells. We used the same gene set of 657 regulatory genes as described in the single-gene network model in Sec. 2.4, including 388 TFs, 187 kinases, and 71 phosphatases. CellOracle was applied to the entire dataset in the same procedure as in the Perturb-seq data. The D-SPIN network construction also followed the same procedure as in the Perturb-seq dataset. With the base GRN from CellOracle, we applied *ℓ*_1_ regularization on edges outside the base GRN with *λ*_1_ = 0.02. Without the base network, we regularized all edges with *λ*_1_ = 0.01. As the complete list of targets of many small molecules in the dataset is not yet clear, we did not provide prior estimation of the response vectors, but regularized the response vector to be sparse relative to the control sample with *λ*_1_ = 0.01 and regularized its magnitude with *λ*_2_ = 0.005.

The inferred regulatory network was compared with the String-db database, as the ChIP-seq database does not contain regulations by the kinases and phosphatases. We used the physical interaction (v12.0) network from the String-db database [23], and selected edges with an evidence score larger than 0.8 as the reference network. The quality of inferred networks was evaluated by the early accuracy rate and AUROC in the same way as the Perturb-seq dataset.

### 2.12 Discovering regulatory network modules and clustering perturbation responses

To gain insight into the structure and modular organization of the regulatory networks inferred by D-SPIN, we applied the Leiden algorithm for graph community detection to both Perturb-seq and drug profiling datasets. The Leiden algorithm identifies densely connected subnetworks, or communities, by optimizing a modularity score that reflects the density of positive interactions within communities compared to a random network. For networks with both positive and negative interactions, the following score function is employed [32].

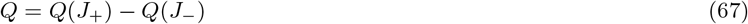

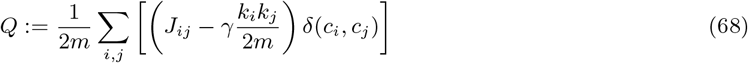

where the regulatory network is split into *J*_+_ for positive and *J*_−_ for negative interactions. In the score function *Q, m* is the sum of all edge weights, *k*_*i*_ denotes the sum of weights for edges connected to node *i, c*_*i*_ is the community to which *i* belongs, and *γ* is the resolution parameter. We set *γ* = 2 to achieve a finer resolution of module discovery, while the presented result is robust with a range of *γ* from 1.05 to 2.5. The Leiden algorithm partitions the network into distinct communities, whose biological functions were inferred from the gene programs they contained.

In addition to analyzing the network architecture, we also clustered the perturbation response vectors to uncover interaction patterns between perturbations and the network. In general, we computed the relative response 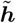 for each ***h*** by subtracting the average ***h*** of control samples. Effective perturbations were then selected by comparing with the distribution of relative responses in control samples.

In the Perturb-seq dataset, we selected 1,755 effective perturbations with *p <* 0.05 under Bonferroni multi-test correction using non-targeting guide RNA as the null distribution. We clustered perturbations into groups because the perturbation impacts are noisy due to the low cell number in each condition and varying knockdown efficiencies. We used the Leiden algorithm for better clustering of the noisy perturbation responses. We built an 8-nearest-neighborhood graph and ran Leiden algorithms with resolution *γ* = 3 to partition the perturbation response vectors into 40 groups. The relative responses of guide groups were defined as the average relative responses in the group.

In the program-level network model drug profiling dataset, due to some batches only having a single control sample, a unified statistical test for the significance of perturbation response vectors is not viable. We scored each drug by the negative log probability density function at 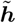 of a multivariate Gaussian distribution with zero mean and diagonal covariance. The covariance was determined by the variance of control samples of the same experimental batch if there were more than 1 control sample, otherwise by the variance of all control samples in the dataset. We selected 158 candidate drugs with a score above 20, and performed K-means clustering with 20 clusters. We found that some drug clusters were putative experimental batch effects, as these contained biochemically different drugs from the same experimental batch. We also removed the drug cluster where a complete loss of macrophage population was observed, as the loss of macrophage population was not reproducible in other experiments, and likely resulted from cell adhesion to the culture plate. The remaining drug clusters were then manually reviewed and consolidated based on their biological impacts, resulting in 70 drugs in 7 major categories.

In the gene-level network model drug profiling dataset, we selected the relative response vectors of all identified immune-inhibitory drugs (strong inhibitors, weak inhibitors I, weak inhibitors II, and glucocorticoids) and performed hierarchical clustering with scipy.cluster.hierarchy. The primary targets of each drug cluster were then identified through databases and manual lookup [33].

### 2.13 Automated identification of regulatory strategies and network coarse-graining

The response vectors can further reveal the information processing of the regulatory network by elucidating how perturbation responses are organized across network modules. The response vectors not only identify perturbation impacts on individual programs, but also provide a global view of how cells respond to the stress induced by the perturbation. To further consolidate perturbation groups, typical measures such as Euclidean distance are not sufficient, as each gene program is not an independent dimension but interacts closely with other gene programs through the regulatory network. Therefore, we used a graph clustering strategy to integrate information of program interactions into the clustering.

Specifically, we built an extended network by also including perturbations as nodes of the network. The extended network encompassed both interactions between gene programs and interactions between perturbations and programs. Formally, the adjacency matrix of the extended network is

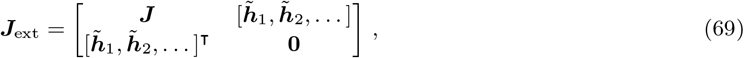

where 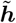 is the average relative response for each perturbation group defined in Sec. 2.12. We then ran the Leiden algorithm on the extended network to detect perturbations that had similar impacts at the network level. In Leiden algorithms, modules of gene programs were fixed, and we only partitioned the perturbations. Perturbations were partitioned into the same community if they influenced gene programs that play analogous roles within the regulatory network. We set *γ* = 1 in the Leiden algorithm, while the algorithm always generates 4 perturbation clusters for *γ* from 0.5 to 2.5, and the presented cluster assignment result is robust with a range of *γ* from 0.9 to 1.45.

Through graph clustering, perturbation groups were assigned to modules in the regulatory network. Based on the score function Eq. 68 of the Leiden algorithm, this suggested that the perturbations predominantly activated gene programs within the module and were more likely to inhibit the gene programs outside the module. Conceptually, this correspondence between perturbations and modules establishes a connection of homeostasis regulation logic, that the cell upregulates the gene programs in the module in response to the loss of function caused by the perturbations. Therefore, we named the global strategy of perturbation responses according to the gene module with which the perturbations were grouped. For example, perturbations that were assigned to the Translation module were named “Translation up-regulation strategy”.

We further visualized the global organization of biological functions and perturbation response strategies with a minimal network derived from the D-SPIN model by coarse-graining. In this simplified representation, both gene program modules and response strategies were condensed into single network nodes, as programs in the same module mostly have positive interactions inside and collectively contribute to a specific biological function. In the minimal network, the interactions between modules were determined by the most significant positive or negative interactions across their constituent gene programs. Similarly, the sign of the interaction between a gene module and a perturbation strategy was based on whether the activating or inhibiting interactions were stronger on average across the constituent gene programs and perturbation groups, and the strength was set to the average strength. As a visualization tool, the minimal network spotlighted the major response strategies employed by the network. For example, the impairment of ribosome subunits induced downregulation of the Translation module and upregulation of the Degradation module.

### 2.14 Comparison between distributions generated by D-SPIN and data

To access the alignment between samples generated from D-SPIN and experimental data (or synthetic data as in Sec. 2.6), we employed several measures, including Uniform Manifold Approximation and Projection (UMAP), cell state distributions, distribution moments, optimal transport, and exact probability distribution.

For small networks, the number of possible states ***s*** was relatively small, and the distribution defined by D-SPIN could be explicitly computed, such as the HSC network and the cell-cycle subnetwork of the Perturb-seq data. We directly took samples from the D-SPIN distributions, computed the joint embedding of D-SPIN samples and experimental data with UMAP, and computed Leiden clustering on the nearest-neighborhood graph obtained during the computation of UMAP. As the states in the model are discretized states [−1, 0, 1], to avoid the neighborhood graph being disconnected due to frequent states forming cliques between themselves, we added a Gaussian noise on sample states with zero mean and 0.1 standard deviation.

With the drug profiling dataset, for a comprehensive evaluation of whether the state distribution computed by D-SPIN is generalizable beyond training data, we trained a program-level D-SPIN model only with 50% of the data in each condition, and compared the cell state distribution with the other 50% held-out testing cell states. To reduce sampling noise, for each condition, we divided each Leiden cluster into two groups of the same size (or differing by 1), and randomly assigned one as training data and the other as testing data.

For 30-node networks, the total 3^*M*^ possible states were impossible to enumerate, and the network encompassed a more complex landscape of cell states, such as the network of the drug profiling dataset. We therefore sampled 10^6^ samples by MCMC (Sec. 1.3) for each experimental condition from the D-SPIN model trained with all the data (full model) or trained with training data only (half-data model), and performed the following analysis.

1. (Full model) UMAP embeddings of samples from the D-SPIN model. In the drug profiling dataset, there were more than 1,000 conditions, and the total number of D-SPIN-generated states was more than 10^9^ (10^6^ for each condition) and intractable for joint UMAP embedding. Therefore, we projected the states generated by D-SPIN to the batch-corrected UMAP of the experimental data with over 1 million cells (Sec. 2.2). Specifically, for each D-SPIN sample, we mapped it to the nearest neighbor of the discretized states in the experimental data. In the case of multiple nearest neighbors, the probability was evenly distributed between these cells. In this way, the 10^6^ samples from D-SPIN for each condition defined a distribution over the entire observed dataset, and could be visualized on the UMAP. We sampled 10^5^ cells from the distribution to render the UMAP for D-SPIN samples in Figure 5E.
2. (Full model) Similarity of distribution on cell subtypes. The projected distribution over the whole dataset also defined a distribution over the cell subtypes as the clusters in the UMAP embedding. We computed the cosine similarity between cell subtype distributions of experimental data and the D-SPIN model for each condition. For reference comparison, we also defined two null distributions of cell state: the first null distribution is the uniform distribution over all possible states in the discrete space of [−1, 0, 1]^*M*^ , the second null distribution is the uniform distribution over the drug profiling dataset by pooling all conditions together to reflect the relative cell type abundance. The cosine similarity is presented in Figure 5F.
3. (Half-data model) Gene program expression means and cross-correlation. The first- and second-order moments of distributions are their fundamental characteristics. We computed the Euclidean (Frobenius) norm of the difference between the mean (cross-correlation) of the held-out experimental data and the D-SPIN model and presented it in Figure S5G(i)(ii)
4. (Half-data model) Optimal transport (OT) distance between experimental data and D-SPIN model. OT offers a robust method to quantify the distance between two probability distributions, taking into account the underlying geometry of the data space. For computational tractability, we computed the entropy-regularized Sinkhorn OT distance with the Python package POT (pot.sinkhorn(reg=0.5)) between heldout experimental data and D-SPIN model distributions and presented in Figure S5G(iii). Specifically, as the samples of the D-SPIN model were obtained through MCMC, they were highly noisy at the single-state level. States that appeared only once or twice were likely by chance and led to skewed representations of the true distributions. This noise could be mitigated with neighborhood projection or distribution average in previous metrics, but had a large impact on state-level distribution comparisons, such as OT. Therefore, we filtered out states in the D-SPIN model that appeared fewer than 5 times, which were roughly half of the 10^6^ samples.
5. (Half-data model) Direct comparison of state probabilities. We further directly compared the probability of every single state, presented as scatter plots as in Figure S5H. The probability of the D-SPIN model also filtered out states that appeared fewer than 5 times, as in optimal transport distance computation.

### 2.15 Computing strength of immune inhibitors

The strength of immune inhibitors in the D-SPIN model is quantified by projecting each inhibitor’s relative response vector 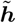 onto a principal direction, denoted by ***e*** with the expression 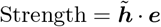. The principal direction, ***e***, is derived from the leading left singular vector of the matrix composed of stacked relative response vectors 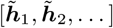 from all immune inhibitors, including strong inhibitors, weak inhibitors I, and weak inhibitors II. The leading singular vector, determined through singular value decomposition (SVD), captures the primary axis of variance of the inhibitor response vectors. It represents the consensus direction of gene program combinations that all immune inhibitors act on.

### 2.16 Categories of drug interactions on gene program level

We define 5 major types of drug interactions on the gene-program level: additive, subadditive, dominant, synergistic, and antagonistic. Denoting the response of two single drugs and drug combinations on a specific gene program as *h*_1_, *h*_2_, *h*_*c*_, we classify different drug interactions based on the sign and magnitude of the single drug and combination responses. We define a tolerance *δ* = 0.5, and the sign of a response *h* is defined as the sign after soft thresholding

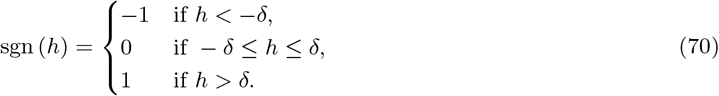

Additive interactions are defined as

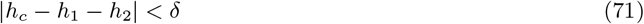

Otherwise, if the two drug responses have the same sign sgn (*h*_1_) sgn (*h*_2_) *>* 0, we assume *h*_1_ *>* 0, *h*_2_ *>* 0 for notation simplicity, and the situation of *h*1 *<* 0, *h*_2_ *<* 0 naturally follows. The interactions are defined by

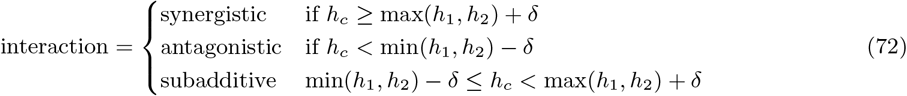

If the two drug responses have opposite signs sgn (*h*_1_) sgn (*h*_2_) *<* 0, we assume *h*_1_ *>* 0, *h*_2_ *<* 0, *h*_*c*_ *>*= 0 for notation simplicity. The interactions are defined by

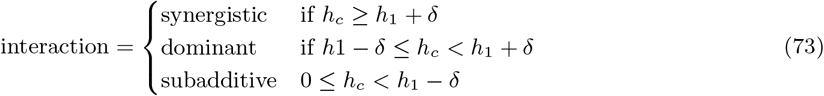

If the two drug responses both have sign 0, the interactions are defined by

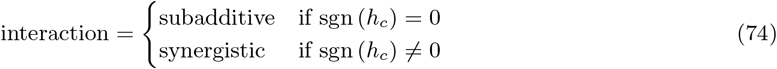

Lastly, if only one of the drug responses has sign 0, we assume *h*_1_ *>* 0, sgn (*h*_2_) = 0 for notational simplicity. The interactions are defined by

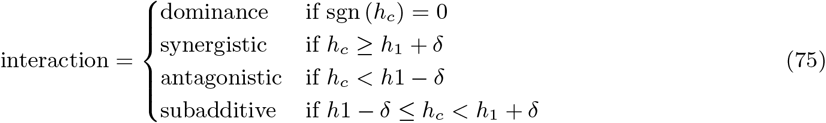

### 2.17 Predicting drug dosage combination response with D-SPIN

We model the combinatorial drug dosage response using a structured equation for each gene program in the form of

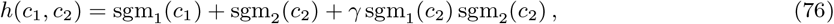

where *c*_1_, *c*_2_ are the log-dosage of the two drugs with base 10, sgm_1_, sgm_2_ are the sigmoid dosage response curve of two single drugs, and *γ* is the interaction strength of the program. Specifically, given a subset of dosage combination conditions, we trained the D-SPIN model on these combination conditions together with all the single-drug dosage response data, activated control, and resting control data. For each program, we fit the following equations from the response ***h***^(*n*)^ learned by D-SPIN.

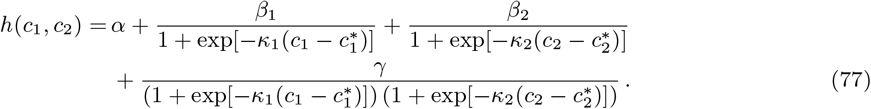

*β*_1_, *β*_2_ quantifies the effect of two single drugs and *γ* quantifies the strength of drug interactions. The fitting was implemented with scipy.optimize.curve fit. Zero drug dosages were replaced with 10^−6^ to avoid infinity. When none of the dosage combination data is provided, the last term is ill-defined, and we set *γ* = 0.

The predicted cell state distribution under unseen dosage combination conditions is evaluated in the same way as in Sec. 2.14. The dosage combination experiments included all pairwise combinations of dasatinib and halcinonide in the following dosages: dasatinib at 0.1, 1, 10, 100, and 1000 nM, and halcinonide at 0.01, 0.1, 1, 10, 100, and 1000 nM. Among the 30 dosage combination conditions (excluding single drug and control conditions), we sampled 10^6^ MCMC samples from D-SPIN using the regulatory network ***J*** learned only with given dosage combinations, and response vectors ***h***^(*n*)^ predicted by the structured equation. We projected the MCMC samples to the nearest neighbor cell of the dosage combination experiment to define a distribution on UMAP coordinates and cell subtypes (Leiden clusters). The predicted cell state distributions were compared with experimental data using UMAP embedding visualizations, cosine similarity of cell subtype distributions, program expression means, cross-correlations, and optimal-transport (OT) distances as described in detail in Sec. 2.14.

### 2.18 Phase diagrams of dosages of drug combination

The regulatory network of the D-SPIN model is able to convert response vectors into distributions of cell states, therefore enabling the prediction and interpolation of cell state transitions between experimental conditions. In the dosage combination experiments, we observed the transition between activated macrophage states, monocyte-like resting states, and combinatorial states of hyper-inhibited M2 macrophages. We used D-SPIN to depict the transitions between the different cell states induced by the change in drug combination dosages.

We constructed a phase diagram of cell states by estimating the average log-likelihood of major cell subtypes under every drug dosage combination. In the analysis, we identified five myeloid states of interest as target states, including (a) Activated macrophage: activated control; (b) resting monocyte: resting control, (c) M2 macrophage: 1000 nM halcinonide; (d) inhibited monocyte: 1000 nM dasatinib, and (e) hyper-inhibited M2 macrophage: 1000 nM dasatinib plus 1000 nM halcinonide. For any given drug combination concentration (*c*_1_, *c*_2_), we interpolated the response vector ***ĥ***(*c*_1_, *c*_2_) (MATLAB fit ‘thinplateinterp’) and evaluated the score *u* of the target cell subtype as the average log-likelihood of individual states {***s***}_Target_ as

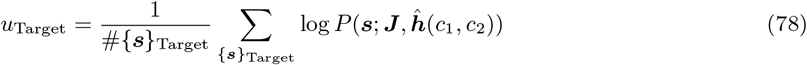

Then, we used softmax across all target cell subtypes to compute the estimated likelihood of each state

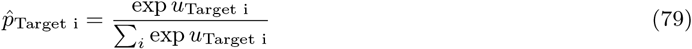

and rendered the phase diagram by the target state with the largest probability. For the D-SPIN model trained with single-drug responses supplemented with one or three dosage combination conditions, the prediction of the phase diagram follows the same procedure after replacing the response vector with the predicted structural equation in Sec. 2.17.

## References

[1] Dennis Bray. Protein molecules as computational elements in living cells. Nature, 376(6538):307–312, 1995.

[2] Aviv Regev and Ehud Shapiro. The π-calculus as an abstraction for biomolecular systems. In Modelling in Molecular Biology, pages 219–266. Springer, 2004.

[3] Aviv Regev and Ehud Shapiro. Cellular abstractions: Cells as computation. Nature, 419(6905):343–343, 2002.

[4] David A Sivak and Matt Thomson. Environmental statistics and optimal regulation. PLoS computational biology, 10(9):e1003826, 2014.

[5] Cameron Sokolik, Yanxia Liu, David Bauer, Jade McPherson, Michael Broeker, Graham Heimberg, Lei S Qi, David A Sivak, and Matt Thomson. Transcription factor competition allows embryonic stem cells to distinguish authentic signals from noise. Cell systems, 1(2):117–129, 2015.

[6] Hao Yuan Kueh, Ameya Champhekar, Stephen L Nutt, Michael B Elowitz, and Ellen V Rothenberg. Positive feedback between pu. 1 and the cell cycle controls myeloid differentiation. Science, 341(6146):670–673, 2013.

[7] Nir Yosef, Alex K Shalek, Jellert T Gaublomme, Hulin Jin, Youjin Lee, Amit Awasthi, Chuan Wu, Katarzyna Karwacz, Sheng Xiao, Marsela Jorgolli, et al. Dynamic regulatory network controlling th17 cell differentiation. Nature, 496(7446):461–468, 2013.

[8] Charlotte Bunne, Yusuf Roohani, Yanay Rosen, Ankit Gupta, Xikun Zhang, Marcel Roed, Theo Alexandrov, Mohammed AlQuraishi, Patricia Brennan, Daniel B Burkhardt, et al. How to build the virtual cell with artificial intelligence: Priorities and opportunities. Cell, 187(25):7045–7063, 2024.

[9] Graham Heimberg, Tony Kuo, Daryle J DePianto, Omar Salem, Tobias Heigl, Nathaniel Diamant, Gabriele Scalia, Tommaso Biancalani, Shannon J Turley, Jason R Rock, et al. A cell atlas foundation model for scalable search of similar human cells. Nature, pages 1–3, 2024.

[10] Haotian Cui, Chloe Wang, Hassaan Maan, Kuan Pang, Fengning Luo, Nan Duan, and Bo Wang. scgpt: toward building a foundation model for single-cell multi-omics using generative ai. Nature Methods, 21(8):1470–1480, 2024.

[11] Eric H Davidson. The regulatory genome: gene regulatory networks in development and evolution. Elsevier, 2010.

[12] Eric H Davidson. Emerging properties of animal gene regulatory networks. Nature, 468(7326):911–920, 2010.

[13] Eric H Davidson and Douglas H Erwin. Gene regulatory networks and the evolution of animal body plans. Science, 311(5762):796–800, 2006.

[14] Lacramioara Bintu, Nicolas E Buchler, Hernan G Garcia, Ulrich Gerland, Terence Hwa, Jané Kondev and Rob Phillips. Transcriptional regulation by the numbers: models. Current opinion in genetics & development, 15(2):116–124, 2005.

[15] Jacques Monod. The growth of bacterial cultures. Annual review of microbiology, 3(1):371–394, 1949.

[16] Arthur B Pardee, François Jacob, and Jacques Monod. The genetic control and cytoplasmic expression of “inducibility” in the synthesis of β-galactosidase by e. coli. Journal of Molecular Biology, 1(2):165– 178, 1959.

[17] Shai S Shen-Orr, Ron Milo, Shmoolik Mangan, and Uri Alon. Network motifs in the transcriptional regulation network of escherichia coli. Nature genetics, 31(1):64–68, 2002.

[18] Ron Milo, Shai Shen-Orr, Shalev Itzkovitz, Nadav Kashtan, Dmitri Chklovskii, and Uri Alon. Network motifs: simple building blocks of complex networks. Science, 298(5594):824–827, 2002.

[19] Günter P Wagner, Mihaela Pavlicev, and James M Cheverud. The road to modularity. Nature Reviews Genetics, 8(12):921–931, 2007.

[20] Tong Ihn Lee, Nicola J Rinaldi, François Robert, Duncan T Odom, Ziv Bar-Joseph, Georg K Gerber, Nancy M Hannett, Christopher T Harbison, Craig M Thompson, Itamar Simon, et al. Transcriptional regulatory networks in saccharomyces cerevisiae. science, 298(5594):799–804, 2002.

[21] Denes Hnisz, Brian J Abraham, Tong Ihn Lee, Ashley Lau, Violaine Saint-André, Alla A Sigova, Heather A Hoke, and Richard A Young. Super-enhancers in the control of cell identity and disease. Cell, 155(4):934–947, 2013.

[22] Christopher T Harbison, D Benjamin Gordon, Tong Ihn Lee, Nicola J Rinaldi, Kenzie D Macisaac, Timothy W Danford, Nancy M Hannett, Jean-Bosco Tagne, David B Reynolds, Jane Yoo, et al. Transcriptional regulatory code of a eukaryotic genome. Nature, 431(7004):99–104, 2004.

[23] Sebastian J Maerkl and Stephen R Quake. A systems approach to measuring the binding energy landscapes of transcription factors. Science, 315(5809):233–237, 2007.

[24] Zhanzhi Hu, Patrick J Killion, and Vishwanath R Iyer. Genetic reconstruction of a functional transcriptional regulatory network. Nature genetics, 39(5):683–687, 2007.

[25] Linda S Huang and Paul W Sternberg. Genetic dissection of developmental pathways. Methods in cell biology, 48:97–122, 1995.

[26] Edwin L Ferguson, Paul W Sternberg, and H Robert Horvitz. A genetic pathway for the specification of the vulval cell lineages of caenorhabditis elegans. Nature, 326(6110):259–267, 1987.

[27] Chiou-Hwa Yuh, Hamid Bolouri, and Eric H Davidson. Genomic cis-regulatory logic: experimental and computational analysis of a sea urchin gene. Science, 279(5358):1896–1902, 1998.

[28] Atray Dixit, Oren Parnas, Biyu Li, Jenny Chen, Charles P Fulco, Livnat Jerby-Arnon, Nemanja D Marjanovic, Danielle Dionne, Tyler Burks, Raktima Raychowdhury, et al. Perturb-seq: dissecting molecular circuits with scalable single-cell rna profiling of pooled genetic screens. cell, 167(7):1853– 1866, 2016.

[29] Joseph M Replogle, Reuben A Saunders, Angela N Pogson, Jeffrey A Hussmann, Alexander Lenail, Alina Guna, Lauren Mascibroda, Eric J Wagner, Karen Adelman, Gila Lithwick-Yanai, et al. Mapping information-rich genotype-phenotype landscapes with genome-scale perturb-seq. Cell, 2022.

[30] Christopher S McGinnis, David M Patterson, Juliane Winkler, Daniel N Conrad, Marco Y Hein, Vasudha Srivastava, Jennifer L Hu, Lyndsay M Murrow, Jonathan S Weissman, Zena Werb, et al. Multi-seq: sample multiplexing for single-cell rna sequencing using lipid-tagged indices. Nature methods, 16(7):619–626, 2019.

[31] Jase Gehring, Jong Hwee Park, Sisi Chen, Matthew Thomson, and Lior Pachter. Highly multiplexed single-cell rna-seq by dna oligonucleotide tagging of cellular proteins. Nature Biotechnology, 38(1):35– 38, 2020.

[32] Sanjay R Srivatsan, José L McFaline-Figueroa, Vijay Ramani, Lauren Saunders, Junyue Cao, Jonathan Packer, Hannah A Pliner, Dana L Jackson, Riza M Daza, Lena Christiansen, et al. Massively multiplex chemical transcriptomics at single-cell resolution. Science, 367(6473):45–51, 2020.

[33] Jesse Zhang, Airol A Ubas, Richard de Borja, Valentine Svensson, Nicole Thomas, Neha Thakar, Ian Lai, Aidan Winters, Umair Khan, Matthew G Jones, et al. Tahoe-100m: A giga-scale single-cell perturbation atlas for context-dependent gene function and cellular modeling. bioRxiv, pages 2025–02, 2025.

[34] Vân Anh Huynh-Thu, Alexandre Irrthum, Louis Wehenkel, and Pierre Geurts. Inferring regulatory networks from expression data using tree-based methods. PloS one, 5(9):e12776, 2010.

[35] Thomas Moerman, Sara Aibar Santos, Carmen Bravo González-Blas, Jaak Simm, Yves Moreau, Jan Aerts, and Stein Aerts. Grnboost2 and arboreto: efficient and scalable inference of gene regulatory networks. Bioinformatics, 35(12):2159–2161, 2019.

[36] Thalia E Chan, Michael PH Stumpf, and Ann C Babtie. Gene regulatory network inference from single-cell data using multivariate information measures. Cell systems, 5(3):251–267, 2017.

[37] Aditya Pratapa, Amogh P Jalihal, Jeffrey N Law, Aditya Bharadwaj, and TM Murali. Benchmarking algorithms for gene regulatory network inference from single-cell transcriptomic data. Nature methods, 17(2):147–154, 2020.

[38] John J Hopfield. Neural networks and physical systems with emergent collective computational abilities. Proceedings of the national academy of sciences, 79(8):2554–2558, 1982.

[39] Elad Schneidman, Michael J Berry, Ronen Segev, and William Bialek. Weak pairwise correlations imply strongly correlated network states in a neural population. Nature, 440(7087):1007–1012, 2006.

[40] William Bialek, Andrea Cavagna, Irene Giardina, Thierry Mora, Edmondo Silvestri, Massimiliano Viale, and Aleksandra M Walczak. Statistical mechanics for natural flocks of birds. Proceedings of the National Academy of Sciences, 109(13):4786–4791, 2012.

[41] Narayana P Santhanam and Martin J Wainwright. Information-theoretic limits of selecting binary graphical models in high dimensions. IEEE Transactions on Information Theory, 58(7):4117–4134, 2012.

[42] Alex H Lang, Hu Li, James J Collins, and Pankaj Mehta. Epigenetic landscapes explain partially reprogrammed cells and identify key reprogramming genes. PLoS computational biology, 10(8):e1003734, 2014.

[43] Andrew E Teschendorff and Andrew P Feinberg. Statistical mechanics meets single-cell biology. Nature Reviews Genetics, 22(7):459–476, 2021.

[44] Adam A Margolin, Ilya Nemenman, Katia Basso, Chris Wiggins, Gustavo Stolovitzky, Riccardo Dalla Favera, and Andrea Califano. Aracne: an algorithm for the reconstruction of gene regulatory networks in a mammalian cellular context. In BMC bioinformatics, volume 7, pages 1–15. Springer, 2006.

[45] Florian Markowetz and Rainer Spang. Inferring cellular networks–a review. BMC bioinformatics, 8(6):S5, 2007.

[46] Julian Besag. Spatial interaction and the statistical analysis of lattice systems. Journal of the Royal Statistical Society: Series B (Methodological), 36(2):192–225, 1974.

[47] Erik Aurell and Magnus Ekeberg. Inverse ising inference using all the data. Physical review letters, 108(9):090201, 2012.

[48] Pradeep Ravikumar, Martin J Wainwright, and John D Lafferty. High-dimensional ising model selection using ℓ1-regularized logistic regression. The Annals of Statistics, pages 1287–1319, 2010.

[49] H Chau Nguyen, Riccardo Zecchina, and Johannes Berg. Inverse statistical problems: from the inverse ising problem to data science. Advances in Physics, 66(3):197–261, 2017.

[50] Assen Roguev, Sourav Bandyopadhyay, Martin Zofall, Ke Zhang, Tamas Fischer, Sean R Collins, Hongjing Qu, Michael Shales, Han-Oh Park, Jacqueline Hayles, et al. Conservation and rewiring of functional modules revealed by an epistasis map in fission yeast. science, 322(5900):405–410, 2008.

[51] Eran Segal, Michael Shapira, Aviv Regev, Dana Pe’er, David Botstein, Daphne Koller, and Nir Friedman. Module networks: identifying regulatory modules and their condition-specific regulators from gene expression data. Nature genetics, 34(2):166–176, 2003.

[52] Xiaoqiao Chen, Sisi Chen, and Matt Thomson. Minimal gene set discovery in single-cell mrna-seq datasets with activesvm. Nature Computational Science, 2(6):387–398, 2022.

[53] Seungjin Choi. Algorithms for orthogonal nonnegative matrix factorization. In 2008 ieee international joint conference on neural networks (ieee world congress on computational intelligence), pages 1828– 1832. IEEE, 2008.

[54] Stephen A Vavasis. On the complexity of nonnegative matrix factorization. SIAM Journal on Optimization, 20(3):1364–1377, 2010.

[55] Graham Heimberg, Rajat Bhatnagar, Hana El-Samad, and Matt Thomson. Low dimensionality in gene expression data enables the accurate extraction of transcriptional programs from shallow sequencing. Cell systems, 2(4):239–250, 2016.

[56] Sisi Chen, Paul Rivaud, Jong H Park, Tiffany Tsou, Emeric Charles, John R Haliburton, Flavia Pichiorri, and Matt Thomson. Dissecting heterogeneous cell populations across drug and disease conditions with popalign. Proceedings of the National Academy of Sciences, 117(46):28784–28794, 2020.

[57] Jan Krumsiek, Carsten Marr, Timm Schroeder, and Fabian J Theis. Hierarchical differentiation of myeloid progenitors is encoded in the transcription factor network. PloS one, 6(8):e22649, 2011.

[58] Judea Pearl. Reverend bayes on inference engines: A distributed hierarchical approach. In Probabilistic and Causal Inference: The Works of Judea Pearl, pages 129–138. ACM, 2022.

[59] Judea Pearl. Evidential reasoning using stochastic simulation of causal models. Artificial intelligence, 32(2):245–257, 1987.

[60] Barry C Arnold and S James Press. Compatible conditional distributions. Journal of the American Statistical Association, 84(405):152–156, 1989.

[61] Leland H Hartwell, John J Hopfield, Stanislas Leibler, and Andrew W Murray. From molecular to modular cell biology. Nature, 402(Suppl 6761):C47–C52, 1999.

[62] Jialong Jiang, David A Sivak, and Matt Thomson. Active learning of spin network models. arXiv preprint 1903.10474, 2019.

[63] Mark Newman. Networks. Oxford university press, 2018.

[64] Albert-László Barabási. Scale-free networks: a decade and beyond. science, 325(5939):412–413, 2009.

[65] Anna D Broido and Aaron Clauset. Scale-free networks are rare. Nature communications, 10(1):1017, 2019.

[66] Brad T Sherman, Ming Hao, Ju Qiu, Xiaoli Jiao, Michael W Baseler, H Clifford Lane, Tomozumi Imamichi, and Weizhong Chang. David: a web server for functional enrichment analysis and functional annotation of gene lists (2021 update). Nucleic Acids Res, 10, 2022.

[67] Maxim V Kuleshov, Matthew R Jones, Andrew D Rouillard, Nicolas F Fernandez, Qiaonan Duan, Zichen Wang, Simon Koplev, Sherry L Jenkins, Kathleen M Jagodnik, Alexander Lachmann, et al. Enrichr: a comprehensive gene set enrichment analysis web server 2016 update. Nucleic acids research, 44(W1):W90–W97, 2016.

[68] Damian Szklarczyk, Annika L Gable, Katerina C Nastou, David Lyon, Rebecca Kirsch, Sampo Pyysalo, Nadezhda T Doncheva, Marc Legeay, Tao Fang, Peer Bork, et al. The string database in 2021: customizable protein–protein networks, and functional characterization of user-uploaded gene/measurement sets. Nucleic acids research, 49(D1):D605–D612, 2021.

[69] Melanie de Almeida, Matthias Hinterndorfer, Hanna Brunner, Irina Grishkovskaya, Kashish Singh, Alexander Schleiffer, Julian Jude, Sumit Deswal, Robert Kalis, Milica Vunjak, et al. Akirin2 controls the nuclear import of proteasomes in vertebrates. Nature, 599(7885):491–496, 2021.

[70] Daniel Schraivogel, Andreas R Gschwind, Jennifer H Milbank, Daniel R Leonce, Petra Jakob, Lukas Mathur, Jan O Korbel, Christoph A Merten, Lars Velten, and Lars M Steinmetz. Targeted perturb-seq enables genome-scale genetic screens in single cells. Nature methods, 17(6):629–635, 2020.

[71] Kenji Kamimoto, Blerta Stringa, Christy M Hoffmann, Kunal Jindal, Lilianna Solnica-Krezel, and Samantha A Morris. Dissecting cell identity via network inference and in silico gene perturbation. Nature, 614(7949):742–751, 2023.

[72] Robert J Prill, Daniel Marbach, Julio Saez-Rodriguez, Peter K Sorger, Leonidas G Alexopoulos, Xiaowei Xue, Neil D Clarke, Gregoire Altan-Bonnet, and Gustavo Stolovitzky. Towards a rigorous assessment of systems biology models: the dream3 challenges. PloS one, 5(2):e9202, 2010.

[73] Deniz Seçilmiş, Thomas Hillerton, Andreas Tjärnberg, Sven Nelander, Torbjörn EM Nordling, and Erik LL Sonnhammer. Knowledge of the perturbation design is essential for accurate gene regulatory network inference. Scientific reports, 12(1):16531, 2022.

[74] Carmen Bravo González-Blas, Seppe De Winter, Gert Hulselmans, Nikolai Hecker, Irina Matetovici, Valerie Christiaens, Suresh Poovathingal, Jasper Wouters, Sara Aibar, and Stein Aerts. Scenic+: single-cell multiomic inference of enhancers and gene regulatory networks. Nature methods, 20(9):1355–1367, 2023.

[75] Trevor Hastie, Robert Tibshirani, Jerome H Friedman, and Jerome H Friedman. The elements of statistical learning: data mining, inference, and prediction, volume 2. Springer, 2009.

[76] Vincent A Traag, Ludo Waltman, and Nees Jan Van Eck. From louvain to leiden: guaranteeing well-connected communities. Scientific reports, 9(1):5233, 2019.

[77] Andrea Schüler, Maike Schwieger, Afra Engelmann, Kristoffer Weber, Stefan Horn, Ursula Müller, Michael A Arnold, Eric N Olson, and Carol Stocking. The mads transcription factor mef2c is a pivotal modulator of myeloid cell fate. Blood, The Journal of the American Society of Hematology, 111(9):4532–4541, 2008.

[78] Zeinab Albadry M Zahran, Xiaorong Gu, Babal K Jha, and Yogenthiran Saunthararajah. Bcr-abl1 dislocates npm1, pu. 1 and klf1 into cytoplasm to thereby skew granulo-monocytic and impede erythroid differentiation. Blood, 142:1375, 2023.

[79] Chiara Milanese, Cíntia R Bombardieri, Sara Sepe, Sander Barnhoorn, César Payán-Goméz, Donatella Caruso, Matteo Audano, Silvia Pedretti, Wilbert P Vermeij, Renata MC Brandt, et al. Dna damage and transcription stress cause atp-mediated redesign of metabolism and potentiation of anti-oxidant buffering. Nature communications, 10(1):4887, 2019.

[80] Miaomiao Yang, Yanming Lu, Weilan Piao, and Hua Jin. The translational regulation in mtor pathway. Biomolecules, 12(6):802, 2022.

[81] Daniella M Schwartz, Yuka Kanno, Alejandro Villarino, Michael Ward, Massimo Gadina, and John J O’Shea. Jak inhibition as a therapeutic strategy for immune and inflammatory diseases. Nature reviews Drug discovery, 16(12):843–862, 2017.

[82] Wen Zhou, Mary A Yui, Brian A Williams, Jina Yun, Barbara J Wold, Long Cai, and Ellen V Rothenberg. Single-cell analysis reveals regulatory gene expression dynamics leading to lineage commitment in early t cell development. Cell systems, 9(4):321–337, 2019.

[83] Bram Van de Sande, Joon Sang Lee, Euphemia Mutasa-Gottgens, Bart Naughton, Wendi Bacon, Jonathan Manning, Yong Wang, Jack Pollard, Melissa Mendez, Jon Hill, et al. Applications of single-cell rna sequencing in drug discovery and development. Nature Reviews Drug Discovery, pages 1–25, 2023.

[84] C Domínguez Conde, C Xu, LB Jarvis, DB Rainbow, SB Wells, T Gomes, SK Howlett, O Suchanek, K Polanski, HW King, et al. Cross-tissue immune cell analysis reveals tissue-specific features in humans. Science, 376(6594):eabl5197, 2022.

[85] Edwin T Jaynes. Information theory and statistical mechanics. Physical review, 106(4):620, 1957.

[86] Thibaut Desgeorges, Giorgio Caratti, Rémi Mounier, Jan Tuckermann, and Bénédicte Chazaud. Glucocorticoids shape macrophage phenotype for tissue repair. Frontiers in immunology, 10:1591, 2019.

[87] Yan-Cun Liu, Xian-Biao Zou, Yan-Fen Chai, and Yong-Ming Yao. Macrophage polarization in inflammatory diseases. International journal of biological sciences, 10(5):520, 2014.

[88] Ctibor Skuta, Martin Popr, Tomas Muller, Jindrich Jindrich, Michal Kahle, David Sedlak, Daniel Svozil, and Petr Bartunek. Probes & drugs portal: an interactive, open data resource for chemical biology. Nature methods, 14(8):759–760, 2017.

[89] David S Wishart, Craig Knox, An Chi Guo, Savita Shrivastava, Murtaza Hassanali, Paul Stothard, Zhan Chang, and Jennifer Woolsey. Drugbank: a comprehensive resource for in silico drug discovery and exploration. Nucleic acids research, 34(uppl 1):D668–D672, 2006.

[90] Gerard Manning, David B Whyte, Ricardo Martinez, Tony Hunter, and Sucha Sudarsanam. The protein kinase complement of the human genome. Science, 298(5600):1912–1934, 2002.

[91] Mark J Chen, Jack E Dixon, and Gerard Manning. Genomics and evolution of protein phosphatases. Science signaling, 10(474):eaag1796, 2017.

[92] Clifford A Lowell. Src-family kinases: rheostats of immune cell signaling. Molecular immunology, 41(6-7):631–643, 2004.

[93] Roland Lang, Michael Hammer, and Jörg Mages. Dusp meet immunology: dual specificity mapk phosphatases in control of the inflammatory response. The Journal of Immunology, 177(11):7497– 7504, 2006.

[94] Ian Gans, Ellen I Hartig, Shusen Zhu, Andrea R Tilden, Lucie N Hutchins, Nathaniel J Maki, Joel H Graber, and James A Coffman. Klf9 is a key feedforward regulator of the transcriptomic response to glucocorticoid receptor activity. Scientific reports, 10(1):11415, 2020.

[95] Heng Yang, Lin Xia, Jian Chen, Shuqing Zhang, Vincent Martin, Qingqing Li, Shangqing Lin, Jinfeng Chen, Joseph Calmette, Min Lu, et al. Stress–glucocorticoid–tsc22d3 axis compromises therapy-induced antitumor immunity. Nature medicine, 25(9):1428–1441, 2019.

[96] Hwijin Kim. The transcription factor mafb promotes anti-inflammatory m2 polarization and cholesterol efflux in macrophages. Scientific reports, 7(1):7591, 2017.

[97] Yin Gao, Peng Fang, Wen-Jin Li, Jian Zhang, Guang-Ping Wang, Duan-Feng Jiang, and Fang-Ping Chen. Lncrna neat1 sponges mir-214 to regulate m2 macrophage polarization by regulation of b7-h3 in multiple myeloma. Molecular immunology, 117:20–28, 2020.

[98] Jessica Hoppstädter and Alaina J Ammit. Role of dual-specificity phosphatase 1 in glucocorticoid-driven anti-inflammatory responses. Frontiers in immunology, 10:1446, 2019.

[99] Andrew R Clark, Joana RS Martins, and Carmen R Tchen. Role of dual specificity phosphatases in biological responses to glucocorticoids. Journal of Biological Chemistry, 283(38):25765–25769, 2008.

[100] Lian Wang, Yanghui Zhu, Nan Zhang, Yali Xian, Yu Tang, Jing Ye, Fekrazad Reza, Gu He, Xiang Wen, and Xian Jiang. The multiple roles of interferon regulatory factor family in health and disease. Signal Transduction and Targeted Therapy, 9(1):282, 2024.

[101] Daniel Romaus-Sanjurjo, Junmi M Saikia, Hugo J Kim, Kristen M Tsai, Geneva Q Le, and Binhai Zheng. Overexpressing eukaryotic elongation factor 1 alpha (eef1a) proteins to promote corticospinal axon repair after injury. Cell Death Discovery, 8(1):390, 2022.

[102] Chunhong Yan, Dan Lu, Tsonwin Hai, and Douglas D Boyd. Activating transcription factor 3, a stress sensor, activates p53 by blocking its ubiquitination. The EMBO journal, 24(13):2425–2435, 2005.

[103] Jianfeng Shu, Jinni Jiang, Xiaofang Wang, Xuejie Yang, Guofang Zhao, and Ting Cai. Mdm2 provides top2 poison resistance by promoting proteolysis of top2βcc in a p53-independent manner. Cell Death & Disease, 15(1):83, 2024.

[104] Sara Zaccara, Toma Tebaldi, C Pederiva, Yari Ciribilli, Alessandra Bisio, and Alberto Inga. p53-directed translational control can shape and expand the universe of p53 target genes. Cell Death & Differentiation, 21(10):1522–1534, 2014.

[105] Constance Qiao Xin Yeo, Irina Alexander, Zhaoru Lin, Shuhui Lim, Obed Akwasi Aning, Ramesh Kumar, Kanda Sangthongpitag, Vishal Pendharkar, Vincent HB Ho, and Chit Fang Cheok. p53 maintains genomic stability by preventing interference between transcription and replication. Cell reports, 15(1):132–146, 2016.

[106] Yung-Chih Cheng, Andrew Snavely, Lee B Barrett, Xuefei Zhang, Crystal Herman, Devlin J Frost, Priscilla Riva, Ivan Tochitsky, Riki Kawaguchi, Bhagat Singh, et al. Topoisomerase i inhibition and peripheral nerve injury induce dna breaks and atf3-associated axon regeneration in sensory neurons. Cell reports, 36(10), 2021.

[107] Naama Geva-Zatorsky, Erez Dekel, Ariel A Cohen, Tamar Danon, Lydia Cohen, and Uri Alon. Protein dynamics in drug combinations: a linear superposition of individual-drug responses. Cell, 140(5):643– 651, 2010.

[108] Yanhua Yang, Shujun Xia, Lu Zhang, Wenhan Wang, Lin Chen, and Weiwei Zhan. Mir-324-5p/ptprd/cebpd axis promotes papillary thyroid carcinoma progression via microenvironment alteration. Cancer Biology & Therapy, 21(6):522–532, 2020.

[109] Yu Shi, Bin Zhang, Jian Zhu, Wu Huang, Bin Han, Qilong Wang, Chunjian Qi, Minghai Wang, and Fang Liu. mir-106b-5p inhibits irf1/ifn-β signaling to promote m2 macrophage polarization of glioblastoma. OncoTargets and therapy, pages 7479–7492, 2020.

[110] Michael A Cannarile, Martin Weisser, Wolfgang Jacob, Anna-Maria Jegg, Carola H Ries, and Dominik Rüttinger. Colony-stimulating factor 1 receptor (csf1r) inhibitors in cancer therapy. Journal for immunotherapy of cancer, 5(1):53, 2017.

[111] Ruo-Yu Ma, Annabel Black, and Bin-Zhi Qian. Macrophage diversity in cancer revisited in the era of single-cell omics. Trends in immunology, 43(7):546–563, 2022.

[112] Lacramioara Bintu, Nicolas E Buchler, Hernan G Garcia, Ulrich Gerland, Terence Hwa, Jane Kondev, Thomas Kuhlman, and Rob Phillips. Transcriptional regulation by the numbers: applications. Current opinion in genetics & development, 15(2):125–135, 2005.

[113] Jeremy Gunawardena. A linear framework for time-scale separation in nonlinear biochemical systems. PloS one, 7(5):e36321, 2012.

[114] Mohammad Lotfollahi, F Alexander Wolf, and Fabian J Theis. scgen predicts single-cell perturbation responses. Nature methods, 16(8):715–721, 2019.

[115] Mohammad Lotfollahi, Anna Klimovskaia Susmelj, Carlo De Donno, Leon Hetzel, Yuge Ji, Ignacio L Ibarra, Sanjay R Srivatsan, Mohsen Naghipourfar, Riza M Daza, Beth Martin, et al. Predicting cellular responses to complex perturbations in high-throughput screens. Molecular Systems Biology, page e11517, 2023.

[116] Yusuf Roohani, Kexin Huang, and Jure Leskovec. Predicting transcriptional outcomes of novel multigene perturbations with gears. Nature Biotechnology, pages 1–9, 2023.

[117] Mingze Dong, Bao Wang, Jessica Wei, Antonio H de O. Fonseca, Curtis J Perry, Alexander Frey, Feriel Ouerghi, Ellen F Foxman, Jeffrey J Ishizuka, Rahul M Dhodapkar, et al. Causal identification of single-cell experimental perturbation effects with cinema-ot. Nature Methods, 20(11):1769–1779, 2023.

[118] Haim Sompolinsky and Annette Zippelius. Relaxational dynamics of the edwards-anderson model and the mean-field theory of spin-glasses. Physical Review B, 25(11):6860, 1982.

[119] Daniel S Fisher and David A Huse. Nonequilibrium dynamics of spin glasses. Physical Review B, 38(1):373, 1988.

[120] Roy J Glauber. Time-dependent statistics of the ising model. Journal of mathematical physics, 4(2):294– 307, 1963.

## References

[1] David Van Dijk, Roshan Sharma, Juozas Nainys, Kristina Yim, Pooja Kathail, Ambrose J Carr, Cassandra Burdziak, Kevin R Moon, Christine L Chaffer, Diwakar Pattabiraman, et al. Recovering gene interactions from single-cell data using data diffusion. Cell, 174(3):716–729, 2018.

[2] Christopher S McGinnis, David M Patterson, Juliane Winkler, Daniel N Conrad, Marco Y Hein, Vasudha Srivastava, Jennifer L Hu, Lyndsay M Murrow, Jonathan S Weissman, Zena Werb, et al. Multi-seq: sample multiplexing for single-cell rna sequencing using lipid-tagged indices. Nature methods, 16(7):619–626, 2019.

[3] H Chau Nguyen, Riccardo Zecchina, and Johannes Berg. Inverse statistical problems: from the inverse ising problem to data science. Advances in Physics, 66(3):197–261, 2017.

[4] Pradeep Ravikumar, Martin J Wainwright, and John D Lafferty. High-dimensional ising model selection using ℓ1-regularized logistic regression. The Annals of Statistics, pages 1287–1319, 2010

[5] Narayana P Santhanam and Martin J Wainwright. Information-theoretic limits of selecting binary graphical models in high dimensions. IEEE Transactions on Information Theory, 58(7):4117–4134, 2012.

[6] Jialong Jiang, David A Sivak, and Matt Thomson. Active learning of spin network models. arXiv preprint arXiv:1903.10474, 2019.

[7] Julian Besag. Spatial interaction and the statistical analysis of lattice systems. Journal of the Royal Statistical Society: Series B (Methodological), 36(2):192–225, 1974.

[8] Erik Aurell and Magnus Ekeberg. Inverse ising inference using all the data. Physical review letters, 108(9):090201, 2012.

[9] Jan Krumsiek, Carsten Marr, Timm Schroeder, and Fabian J Theis. Hierarchical differentiation of myeloid progenitors is encoded in the transcription factor network. PloS one, 6(8):e22649, 2011.

[10] Judea Pearl. Reverend bayes on inference engines: A distributed hierarchical approach. In Probabilistic and Causal Inference: The Works of Judea Pearl, pages 129–138. ACM, 2022.

[11] Judea Pearl. Evidential reasoning using stochastic simulation of causal models. Artificial intelligence, 32(2):245–257, 1987.

[12] Barry C Arnold and S James Press. Compatible conditional distributions. Journal of the American Statistical Association, 84(405):152–156, 1989.

[13] Kenji Kamimoto, Blerta Stringa, Christy M Hoffmann, Kunal Jindal, Lilianna Solnica-Krezel, and Samantha A Morris. Dissecting cell identity via network inference and in silico gene perturbation. Nature, 614(7949):742–751, 2023.

[14] Carmen Bravo González-Blas, Seppe De Winter, Gert Hulselmans, Nikolai Hecker, Irina Matetovici, Valerie Christiaens, Suresh Poovathingal, Jasper Wouters, Sara Aibar, and Stein Aerts. Scenic+: single-cell multiomic inference of enhancers and gene regulatory networks. Nature methods, 20(9):1355–1367, 2023.

[15] Sisi Chen, Paul Rivaud, Jong H Park, Tiffany Tsou, Emeric Charles, John R Haliburton, Flavia Pichiorri, and Matt Thomson. Dissecting heterogeneous cell populations across drug and disease conditions with popalign. Proceedings of the National Academy of Sciences, 117(46):28784–28794, 2020.

[16] Cameron Sokolik, Yanxia Liu, David Bauer, Jade McPherson, Michael Broeker, Graham Heimberg, Lei S Qi, David A Sivak, and Matt Thomson. Transcription factor competition allows embryonic stem cells to distinguish authentic signals from noise. Cell systems, 1(2):117–129, 2015.

[17] Judea Pearl, Madelyn Glymour, and Nicholas P Jewell. Causal inference in statistics: A primer. John Wiley & Sons, 2016.

[18] Joseph M Replogle, Reuben A Saunders, Angela N Pogson, Jeffrey A Hussmann, Alexander Lenail, Alina Guna, Lauren Mascibroda, Eric J Wagner, Karen Adelman, Gila Lithwick-Yanai, et al. Mapping information-rich genotype-phenotype landscapes with genome-scale perturb-seq. Cell, 2022.

[19] Seungjin Choi. Algorithms for orthogonal nonnegative matrix factorization. In 2008 ieee international joint conference on neural networks (ieee world congress on computational intelligence), pages 1828–1832. IEEE, 2008.

[20] Stephen A Vavasis. On the complexity of nonnegative matrix factorization. SIAM Journal on Optimization, 20(3):1364–1377, 2010.

[21] Brad T Sherman, Ming Hao, Ju Qiu, Xiaoli Jiao, Michael W Baseler, H Clifford Lane, Tomozumi Imamichi, and Weizhong Chang. David: a web server for functional enrichment analysis and functional annotation of gene lists (2021 update). Nucleic Acids Res, 10, 2022.

[22] Maxim V Kuleshov, Matthew R Jones, Andrew D Rouillard, Nicolas F Fernandez, Qiaonan Duan, Zichen Wang, Simon Koplev, Sherry L Jenkins, Kathleen M Jagodnik, Alexander Lachmann, et al. Enrichr: a comprehensive gene set enrichment analysis web server 2016 update. Nucleic acids research, 44(W1):W90– W97, 2016.

[23] Damian Szklarczyk, Annika L Gable, Katerina C Nastou, David Lyon, Rebecca Kirsch, Sampo Pyysalo, Nadezhda T Doncheva, Marc Legeay, Tao Fang, Peer Bork, et al. The string database in 2021: customizable protein–protein networks, and functional characterization of user-uploaded gene/measurement sets. Nucleic acids research, 49(D1):D605–D612, 2021.

[24] Dana Pe er, Aviv Regev, Gal Elidan, and Nir Friedman. Inferring subnetworks from perturbed expression profiles. BIOINFORMATICS-OXFORD-, 17:S215–S224, 2001.

[25] Trevor Hastie, Robert Tibshirani, Jerome H Friedman, and Jerome H Friedman. The elements of statistical learning: data mining, inference, and prediction, volume 2. Springer, 2009.

[26] Lisa Holthaus, Daniel Lamp, Anita Gavrisan, Virag Sharma, Anette-Gabriele Ziegler, Martin Jastroch, and Ezio Bonifacio. Cd4+ t cell activation, function, and metabolism are inhibited by low concentrations of dmso. Journal of immunological methods, 463:54–60, 2018.

[27] Romain Lopez, Jeffrey Regier, Michael B Cole, Michael I Jordan, and Nir Yosef. Deep generative modeling for single-cell transcriptomics. Nature methods, 15(12):1053–1058, 2018.

[28] C Domínguez Conde, C Xu, LB Jarvis, DB Rainbow, SB Wells, T Gomes, SK Howlett, O Suchanek, K Polanski, HW King, et al. Cross-tissue immune cell analysis reveals tissue-specific features in humans. Science, 376(6594):eabl5197, 2022.

[29] Aditya Pratapa, Amogh P Jalihal, Jeffrey N Law, Aditya Bharadwaj, and TM Murali. Benchmarking algorithms for gene regulatory network inference from single-cell transcriptomic data. Nature methods, 17(2):147–154, 2020.

[30] Robert J Prill, Daniel Marbach, Julio Saez-Rodriguez, Peter K Sorger, Leonidas G Alexopoulos, Xiaowei Xue, Neil D Clarke, Gregoire Altan-Bonnet, and Gustavo Stolovitzky. Towards a rigorous assessment of systems biology models: the dream3 challenges. PloS one, 5(2):e9202, 2010.

[31] Deniz Seçilmiş, Thomas Hillerton, Andreas Tjärnberg, Sven Nelander, Torbjörn EM Nordling, and Erik LL Sonnhammer. Knowledge of the perturbation design is essential for accurate gene regulatory network inference. Scientific reports, 12(1):16531, 2022.

[32] Vincent A Traag, Ludo Waltman, and Nees Jan Van Eck. From louvain to leiden: guaranteeing well-connected communities. Scientific reports, 9(1):5233, 2019.

[33] David S Wishart, Craig Knox, An Chi Guo, Savita Shrivastava, Murtaza Hassanali, Paul Stothard, Zhan Chang, and Jennifer Woolsey. Drugbank: a comprehensive resource for in silico drug discovery and exploration. Nucleic acids research, 34(uppl 1):D668–D672, 2006.

